# Metaeffector interactions modulate the type III effector-triggered immunity load of *Pseudomonas syringae*

**DOI:** 10.1101/2021.11.18.469137

**Authors:** Alexandre Martel, Bradley Laflamme, Clare Breit-McNally, Fabien Lonjon, Darrell Desveaux, David S. Guttman

**Author notes:** These authors contributed equally (Alexandre Martel, Bradley Laflamme). These authors contributed equally and appear alphabetically (Darrell Desveaux, David S. Guttman).

## Abstract

The bacterial plant pathogen *Pseudomonas syringae* requires type III secreted effectors (T3SEs) for pathogenesis. However, a major facet of plant immunity entails the recognition of a subset of *P. syringae*’s T3SEs by intracellular host receptors in a process called Effector-Triggered Immunity (ETI). Prior work has shown that ETI-eliciting T3SEs are pervasive in the *P. syringae* species complex raising the question of how *P. syringae* mitigates its ETI load to become a successful pathogen. While pathogens can evade ETI by T3SE mutation, recombination, or loss, there is increasing evidence that effector-effector (a.k.a., metaeffector) interactions can suppress ETI. To study the ETI-suppression potential of *P. syringae* T3SE repertoires, we compared the ETI-elicitation profiles of two genetically divergent strains: *P. syringae* pv. *tomato* DC3000 (PtoDC3000) and *P. syringae* pv. *maculicola* ES4326 (PmaES4326), which are both virulent on *Arabidopsis thaliana* but harbour largely distinct effector repertoires. Of the 529 T3SE alleles screened on *A. thaliana* Col-0 from the *P. syringae* T3SE compendium (PsyTEC) [1], 69 alleles from 21 T3SE families elicited ETI in at least one of the two strain backgrounds, while 50 elicited ETI in both backgrounds, resulting in 19 differential ETI responses including two novel ETI-eliciting families: AvrPto1 and HopT1. Although most of these differences were quantitative, three ETI responses were completely absent in one of the pathogenic backgrounds. We performed ETI suppression screens to test if metaeffector interactions contributed to these ETI differences, and found that HopQ1a suppressed AvrPto1m-mediated ETI, while HopG1c and HopF1g suppressed HopT1b-mediated ETI. Overall, these results show that *P. syringae* strains leverage metaeffector interactions and ETI suppression to overcome the ETI load associated with their native T3SE repertoires.

## INTRODUCTION

The bacterial plant pathogen *Pseudomonas syringae* infects hundreds of plant species and is responsible for dozens of agronomically important crop diseases [2–6]. Central to the pathogenesis of *P. syringae* is the type III secretion system (T3SS), which the pathogen uses to inject virulence proteins termed type III secreted effectors (T3SEs or simply effectors) directly into host cells [6]. *P. syringae* T3SEs disrupt a variety of host cellular processes, with many characterized T3SEs disrupting signaling cascades associated with the initial perception of microbes based on conserved molecular patterns (i.e., Pattern-Triggered Immunity, PTI) [7, 8]. Notably, effectors are also central to the plant defense strategy against *P. syringae*, as intracellular host receptors in the Nucleotide-Binding Leucine Rich Repeat (NLR) class can perceive effector activity and activate a powerful immune response termed Effector-Triggered Immunity (ETI), which is often associated with a programmed cell death response at the site of infection called the hypersensitive response (HR) [9].

In a recent study, we defined the *Arabidopsis thaliana – P. syringae* “ETI landscape” by screening for ETI using a compendium of *P. syringae* effectors representing their global diversity (*P. syringae* type III effector compendium, PsyTEC) [1]. The PsyTEC screen interrogated 529 representative T3SE alleles, delivered by *P. syringae* pv. *tomato* DC3000 (PtoDC3000) onto the *A. thaliana* Col-0 accession and identified 59 ETI-eliciting alleles dispersed among 19 distinct T3SE families. When we mapped the distribution of these ETI- elicitor orthologs onto a collection of 494 *P. syringae* strains, we found that 96.8% of strains carry at least one allele that has the potential to elicit ETI, including strains that are pathogenic on *A. thaliana*, which raised the question of how *P. syringae* manages its ETI load? We define ETI-load as the number of host-specific ETI-eliciting T3SEs per strain. ETI load is analogous to ‘mutation load’, which is the genetic burden caused by the accumulation of deleterious mutations in a population. Mutation load, and putatively ETI-load, can substantially reduce the fitness of individuals relative to what it would be without the compromising deleterious mutation or effector.

Microbial pathogens utilize several evolutionary strategies to modulate their ETI-load, with the most commonly invoked strategies being effector diversification via mutation or recombination, and gene loss (i.e., pseudogenization) [9, 10]. For example, *P. syringae* pv. *phaseolicola* 1302A is able to excise a genomic island containing the effector HopAR1 (formerly known as AvrPphB [11]), enabling it to remain virulent on bean plants which can mount a HopAR1-associated ETI response [12]. Likewise, PtoDC3000 carries a heavily truncated version of HopO1c, an effector recognized in *A. thaliana* by the ZAR1 NLR [1], suggesting that functionality of this effector has been sacrificed via a nonsense mutation to avoid ETI-elicitation. One limitation to this strategy is that it will only be generally effective for those effectors that share some functional redundancy with other effectors carried by the same strain. This strategy would be much less likely to be successful for conserved (“core”) effectors such as HopM1 and AvrE1 in *P. syringae*, which are required for virulence [13–15]. Indeed, the diversification or loss of essential and non-redundant effectors may result in significant fitness defects [13, 15, 16]. In such cases, the fine-tuning of effector dosage during infection is likely necessary to optimize virulence benefits and mitigate ETI-load, as may be the case with AvrE1 in PtoDC3000 given that ETI is only observed when AvrE1 is overexpressed [1].

An alternative evolutionary strategy that can be employed by bacteria such as *P. syringae* is to use its effector repertoire to suppress ETI-elicitation. This epistatic process has been termed effector-effector or “metaeffector” interactions [17]. Such interactions may entail a direct antagonistic interaction between the ETI-suppressing and ETI-eliciting effector [18], or they may be indirect interactions due to a shared host target or pathway between an ETI-eliciting and non-eliciting effector [19]. In either case, metaeffector interactions present an alternative path to ETI evasion which avoids the potential fitness detriment associated with gene loss. Given that *P. syringae* strains often deploy large repertoires of more than 30 effectors [20], it is plausible that many of these effectors modulate one another’s activities in a widespread manner.

Substantial work on metaeffector activity has been undertaken in several mammalian pathogens, notably *Legionella pneumophila* [21] and the *Salmonella* genus [22], demonstrating how metaeffector interactions can directly or indirectly coordinate effector activities. In *P. syringae*, metaeffector studies have mainly focused on investigating ETI-suppression. A seminal mechanistic study of *P. syringae* metaeffector interactions in *A. thaliana* is AvrRpm1-mediated ETI suppression by AvrRpt2. Both *P. syringae* effectors AvrRpm1 and AvrRpt2 interact with the *A. thaliana* protein RIN4. The phosphorylation of RIN4 by AvrRpm1 elicits an ETI mediated by the NLR RPM1, while the cleavage of RIN4 by AvrRpt2 elicits an ETI mediated by the NLR RPS2 [23–25]. In an *A. thaliana* line lacking RPS2, the cleavage of RIN4 by AvrRpt2 prevents AvrRpm1 from interacting with RIN4 and activating RPM1-mediated ETI [26, 27]. In turn, AvrRpt2-mediated ETI can be suppressed by HopF2, also through an interaction with RIN4 [19]. Jamir et al. [28] showed that five PtoDC3000 effectors could suppress the HopA1 (formerly HopPsyA) mediated HR in tobacco and *A. thaliana.* In the *P. syringae* pv. *phaseolicola*, AvrB1 (formerly AvrPphC) can suppress ETI associated with HopF1 (formerly AvrPphF) in bean [29]. More recently, a systematic investigation of ETI suppression in PtoDC3000 revealed effectors HopI1 and HopAB1 (formerly AvrPtoB) as major suppressors of ETI-associated cell death in *Nicotiana benthamiana* [30], and another study of the HopZ family found several effectors which suppress HopZ3-mediated ETI [18]. Additionally, many studies over the past few decades have identified ETI suppression events mediated by effectors in *P. syringae* [31–35]. These studies cumulatively point towards metaeffector ETI-suppression being a widespread component of *P. syringae* virulence, though the ETI suppression capacity of individual *P. syringae* strains is poorly understood.

In this study, we explored metaeffector interactions in two *P. syringae* strains, PtoDC3000 and *P. syringae* pv. *maculicola* ES4326 (PmaES4326). Although both strains are highly virulent on *A. thaliana*, they harbour largely distinct effector repertoires, which we hypothesized would produce divergent ETI-suppression profiles on *A. thaliana*. To test this, we screened PsyTEC in PmaES4326 and compared the output to our original PtoDC3000 screen [1]. We found that some effectors from PsyTEC only elicited ETI when expressed in one of the two pathogen backgrounds, leading us to screen for effectors that suppress the ETI response in the virulent background. These screens revealed several PtoDC3000 effectors which are novel suppressors of ETI in *A. thaliana*, despite these effectors mostly having previously defined roles in PTI suppression. Furthermore, our investigation of these differential ETI profiles revealed that the suppression of ETI primarily occurs along quantitative, rather than qualitative, lines. Our results contribute to a growing appreciation for the prominent immune suppression capacities of *P. syringae* effectors.

## METHODS

### Plant material, bacterial strains, and cloning

*A. thaliana* Col-0 plants were grown in Sunshine Mix 1 soil with a 12-hour photoperiod of 150 microeinsteins of light at a constant temperature of 22°C.

The PsyTEC collection of representative *P. syringae* T3SEs was described in [1]. 530 PsyTEC constructs in the MCS2 vector, including 529 T3SE alleles under their native promoters, with chaperones when appropriate, and an empty vector, were mated into PmaES4326 using a modification of a previously described tri-parental mating [1]. Briefly, a 96-pin pinner was used to pin the *Escherichia coli* donor (MCS2::PsyTEC), the *E. coli* helper (pRK600) and PmaES4326 strains onto LB and grown for one day at 28°C. Colonies were then pinned onto KB (streptomycin 100 μg/ml, kanamycin 50 μg/ml) and grown for one day at 28°C. Single colonies for each strain were streak purified on KB (streptomycin 100 μg/ml, kanamycin 50 μg/ml) and then stocked.

We produced pUCP20TC, a tetracycline-resistant Gateway-compatible broad-host-range vector. So do so, we took pUCP20 [36] and swapped its ampicillin resistance marker to tetracycline (Pauline Wang, personal communication). To produce a Gateway-compatible version of pUCP20TC, the Gateway-compatible cassette was excised from pBBR1-MCS-2 via EcoRV digest and cloned into the SmaI site in the polylinker of pUCP20TC. The resulting vector was cloned into *E. coli* DB3.1 and validated via PCR and Sanger sequencing. Putative suppression constructs were synthesized into pDONR207 as in [1] and were transferred to the aforementioned pUCP20TC vector by gateway cloning. Construct sequences are presented in Table S3. These pUCP20TC constructs were mated into *P. syringae* strains harboring ETI-eliciting T3SEs in the MCS2 vector by tri-parental mating [1], resulting in strains harboring three resistances: streptomycin (PmaES4326 genetic background), kanamycin (ETI elicitor in MCS2), and tetracycline (suppressor in pUCP20TC).

**Table 1.** PtoDC3000 and PmaES4326 native T3SE repertoires

| T3SE Allele * | PtoDC3000 | PmaES4326 | T3SE Allele * | PtoDC3000 | PmaES4326 |
| --- | --- | --- | --- | --- | --- |
| AvrE1f | 1 | 0 | HopC1a | 1 | 0 |
| AvrE1i | 0 | 1 | HopD1a | 1 | 1 |
| AvrPto1d | 1 | 0 | HopD1b | 1 | 0 |
| HopA1c | 1 | 0 | HopD1e | 1 | 1 |
| HopAA1b | 1 | 1 | HopD2b | 0 | 1 |
| HopAA1e | 1 | 0 | HopE1a | 1 | 0 |
| HopAA1k | 0 | 1 | HopF1g | 1 | 0 |
| HopAB1j | 1 | 0 | HopG1c | 1 | 0 |
| HopAB1p | 0 | 1 | HopH1a | 1 | 0 |
| HopAB1w | 0 | 1 | HopI1g | 0 | 1 |
| HopAD1a | 1 | 1 | HopI1i | 1 | 0 |
| HopAF1e | 1 | 0 | HopK1d | 1 | 0 |
| HopAF1h | 0 | 1 | HopL1a | 1 | 0 |
| HopAG1g | 2 | 0 | HopM1h | 1 | 0 |
| HopAH1a | 1 | 0 | HopM1v | 0 | 1 |
| HopAH1c | 1 | 0 | HopN1a | 1 | 0 |
| HopAH1i | 1 | 0 | HopO1a | 1 | 0 |
| HopAH1m | 0 | 1 | HopO1b | 1 | 1 |
| HopAH1s | 0 | 1 | HopO1c | 1 | 0 |
| HopAl1d | 1 | 0 | HopO1d | 1 | 0 |
| HopAL1b | 0 | 1 | HopO2f | 1 | 0 |
| HopAM1a | 2 | 0 | HopQ1a | 1 | 1 |
| HopAQ1a | 1 | 1 | HopR1b | 1 | 0 |
| HopAS1b | 2 | 0 | HopR1g | 0 | 2 |
| HopAS1e | 0 | 1 | HopS2a | 1 | 0 |
| HopAT1a | 2 | 0 | HopT1a | 1 | 1 |
| HopAZ1d | 0 | 1 | HopT1b | 1 | 0 |
| HopB1a | 1 | 0 | HopU1a | 1 | 0 |
| HopB2f | 2 | 0 | HopV1d | 1 | 1 |
| HopB2k | 0 | 1 | HopW1g | 0 | 1 |
| HopBD1a | 0 | 1 | HopX1c | 1 | 0 |
| HopBD1g | 0 | 1 | HopX1g | 0 | 1 |
| HopBG1a | 0 | 1 | HopY1c | 1 | 0 |
| HopBM1a | 2 | 0 | HopZ1b | 0 | 1 |
| HopBM1b | 0 | 1 | HopZ1c | 0 | 1 |
| HopBO1d | 0 | 1 | **Total** | **47** | **33** |
| HopC1a | 1 | 0 |  |  |  |

\*Shading depicts T3SE sequence similarity between strains. White: no overlap at the allele or family level. Light grey: both pathogens carry at least one allele of that effector family (e.g., AvrE1). Dark grey: both pathogens carry the same effector allele (e.g., HopAA1b).

### Immunoblotting

Validation of protein expression for strains in this study was performed as previously described [1]. Briefly, protein expression was induced by diluting *P.* syringae strains to an OD_600_ = 0.1 followed by overnight growth in *hrp*-inducing minimal media. 1.5 ml of culture was spun down and resuspended in SDS-loading buffer, followed by two 5-minute incubations at 95°C. These samples were used to run SDS-PAGE gels, followed by immunoblotting with 1:15 000 HA primary antibody and a 1:30 000 anti-rat secondary antibody.

### Bacterial infections, growth assays and hypersensitive response assays

Bacterial spray inoculations, bacterial growth assays and plant imaging were performed as previously described [37]. Briefly, bacterial suspensions (OD_600_ = 0.8 in 10 mM MgSO_4_ supplemented with 0.04% silwet L-77) were sprayed onto 4-week-old plants (approximately 2.5 ml of culture per plant) using Preval sprayers and immediately domed. For visual symptom quantification, plants were imaged periodically for 10 days post-infection (dpi) using a Nikon D5200 DSLR camera affixed with a Nikon 18 to 140 mm DX VR lens. Quantification using the PIDIQ ImageJ macro was performed as previously described [38].

Bacterial growth assays were performed 3 dpi. Four leaf discs (1 cm^2^) were homogenized in 1 ml 10 mM MgSO_4_ using a bead-beater. Following serial dilutions and overnight growth on KB agar (streptomycin 100 μg/ml for PmaES4326 or rifampicin 50 μg/ml for PtoDC3000) at 28°C, colony forming units were counted.

Hypersensitive response assays were performed on 4-5-week-old plants. A bacterial suspension (OD_600_ = 0.2 for PtoDC3000 and PmaES4326, OD_600_ = 0.5 for D36E, in 10 mM MgSO_4_) was infiltrated into the left side of each tested leaf with a blunt end syringe. Tissue collapse was assessed 18-22 hours post infiltration.

### Phylogenetic analysis

We used a previously generated *P. syringae* core genome phylogeny that included the 494 *P. syringae* strains used to generate PsyTEC [1, 39]. The annotated presence or absence of T3SEs within each *P. syringae* strain was determined using previously published metadata [1]. The alignments for the maximum likelihood T3SE family trees presented in Figure 5 were generated using PsyTEC amino acid T3SE allele sequences and MUSCLE [1].

## RESULTS

### Multiple T3SE alleles display strain specific ETI phenotypes

We previously screened PsyTEC for ETI elicitation on the Col-0 accession of *A. thaliana* using the PtoDC3000 pathovar. To determine how the effector repertoire of PtoDC3000 influenced the observed ETI profile, we screened PsyTEC for ETI elicitation using the divergent pathovar PmaES4326. PtoDC3000 is a tomato and *A. thaliana* pathogen found in phylogroup 1, while PmaES4326 is a radish and *A. thaliana* pathogen found in phylogroup 5. These strains are quite dissimilar, with d_N_ = 0.0339, d_S_ = 0.4735 and the two-way average nucleotide identity (ANI) of these two strains is 87.08% (±5.17% sd) from 16444 fragments [40]. The combined effector suites carried by PmaES4326 and PtoDC3000 capture 43 of the 70 *P. syringae* effector protein families, with 36 effector families in PtoDC3000 and 26 effector families in PmaES4326 (19 overlapping families, Jaccard similarity = 0.44). From the allelic perspective, PtoDC3000 and PmaES4326 carry 47 and 33 distinct effector alleles, respectively, with nine shared alleles (Jaccard similarity = 0.13) [20].

We mated the PsyTEC collection of 529 representative T3SE alleles into PmaES4326 and confirmed minimal media induced T3SE protein expression through immunoblotting, observing similar expression profiles to those from PtoDC3000 (Table S1, Figure S1). Overall, protein accumulation of 78% (412/529) of T3SEs was observed in PmaES4326, compared to 76% (402/529) in PtoDC3000, with only 11% (56/529) of T3SE alleles displaying variable expression phenotypes between the two strains (Table S1, Figure S1).

We performed a spray-inoculation ETI screen of all PsyTEC alleles delivered from PmaES4326 on *A. thaliana* Col-0 and quantified macroscopic disease symptoms for each strain using the ImageJ macro PIDIQ to quantify percent yellow (i.e., diseased) tissue [38]. We normalized the percent yellow tissue values within the bounds of our controls, resulting in disease scores ranging between 0.0 (positive ETI control, PmaES4326::HopZ1a) and 1.0 (negative ETI control, PmaES4326::empty vector). Similar to our previous PtoDC3000 screen [1], we defined an ETI disease score threshold that captured the eight well-characterized ETI-eliciting T3SE alleles in this pathosystem (HopZ1a, HopF1r [HopF2a], HopK1a [AvrRps4], AvrRpm1d, AvrRpt2b, AvrB1b, and HopAR1h [AvrPphB]) to determine which PsyTEC alleles elicited ETI (Table S2). This approach resulted in a 0.45 (45%) disease score threshold, as previously determined for PtoDC3000 (Table S2) [1].

Of the 529 PsyTEC effector alleles screened, 69 alleles from 21 T3SE families elicited ETI in at least one strain background, 59 elicited ETI when expressed in the PtoDC3000 background, 60 elicited ETI in the PmaES4326 background, and 50 elicited ETI in both backgrounds, resulting in 19 differential ETI responses and an ETI elicitation Jaccard similarity of 0.72. The T3SE disease scores obtained from our PmaES4326 screen were largely consistent with those of our PtoDC3000 screen, with an average delta disease score of 0.00066 (Table S2, Figure 1, Figure S2, Figure S3). Although the majority of T3SEs obtained similar disease scores in both screens, 19 T3SE alleles from ten T3SE families displayed differential ETI responses across the two strains (Figure 1A). These 19 T3SE alleles were responsible for most of the variation in disease score profiles observed between the two screens (Figure 1B, Figure S3). Ten T3SE alleles from five T3SE families (AvrE1aa, AvrE1o, AvrE1x, AvrPto1m, AvrPto1k, AvrRpt2a, HopO2a, HopO1b, HopT1b and HopT1d) passed our disease score threshold when delivered by PmaES4326 but not when delivered by PtoDC3000, including AvrPto1 and HopT1, two T3SE families that have not previously been described to elicit ETI in *A. thaliana* Col-0 (Figure 1A).

**Figure 1.**
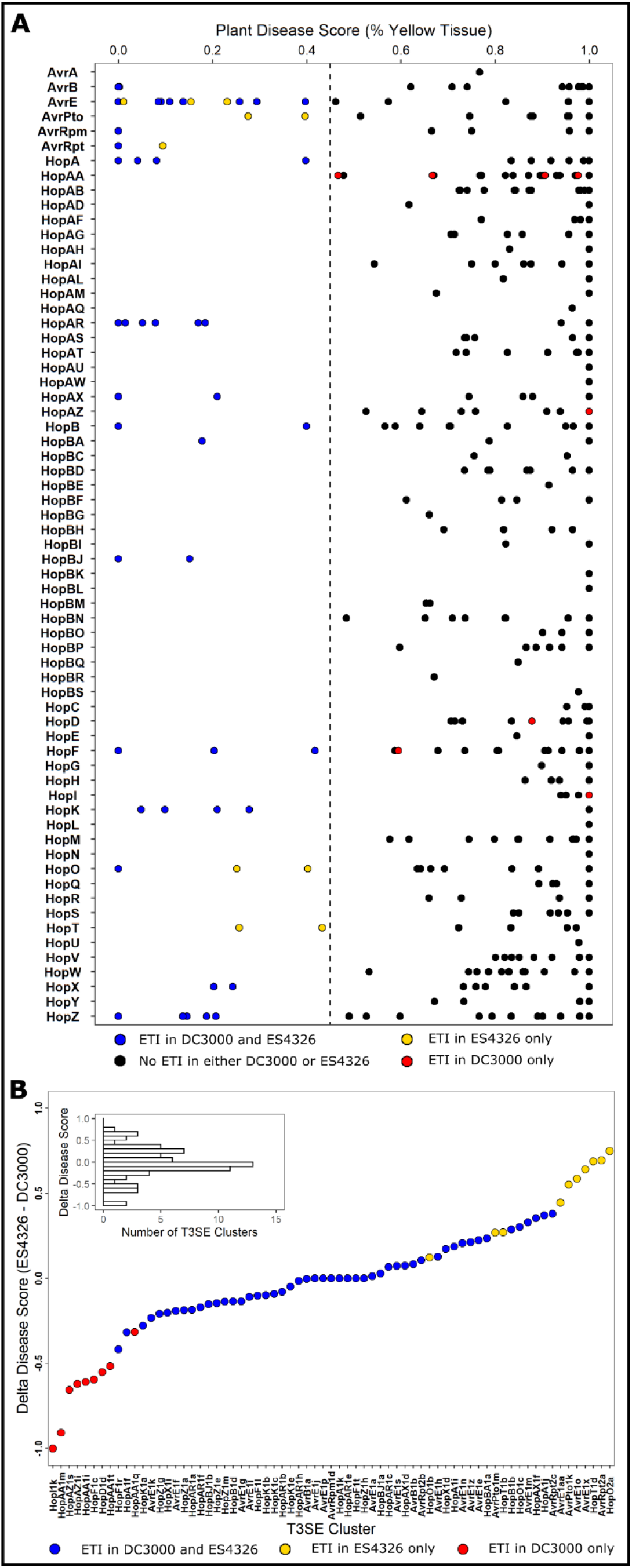
ETI potential of *P. syringae* T3SEs delivered from PmaES4326. **(A)** Plant disease scores of PmaES4326 strains carrying each of the 529 PsyTEC T3SE alleles from 65 families after spray infection of *A. thaliana* accession Col-0. Plant disease scores were determined by assessing the percent of yellow tissue [38] within the bounds of the positive ETI control (HopZ1a, disease score = 0.0) and the negative ETI control (EV, disease score = 1.0). A disease score below 0.45 was used as the threshold to classify ETI responses. Color coding provides a comparison of the current PmaES4326 data to published PtoDC3000 data [1]: black - no ETI when delivered from either genetic background; blue - ETI when delivered from either PtoDC3000 or PmaES4326; red - ETI observed only when delivered from PtoDC3000; and yellow - ETI observed only when delivered from PmaES4326. Detailed data is presented in Table S2. **(B)** The distribution of delta disease scores (PmaES4326 disease score - PtoDC3000 disease score) for all T3SE alleles that elicit ETI in at least one pathogenic background.

The PtoDC3000 AvrPto1 allele (AvrPto1d), which has previously been associated with immunity in certain *A. thaliana* accessions, was not captured in our screen [41]. Further, nine T3SE alleles from five additional T3SE families (HopD1d, HopF1c, HopI1k, HopAA1m, HopAA1t, HopAA1q, HopAA1i, HopAZ1i and HopAZ1s), which were previously identified as ETI-elicitors in the PtoDC3000 screen, did not pass the ETI disease score threshold when delivered from PmaES4326. As such 27.5% (19/69) of T3SE alleles display variable ETI phenotypes, highlighting the significant impact of pathogen genotype on ETI potential.

### Differences in ETI potential are largely quantitative, rather than qualitative

We performed comparative bacterial growth assays between each of the 19 differentially ETI-eliciting T3SE alleles on *A. thaliana* Col-0 to validate our screening results (Figure 2, Figure S4). All 19 T3SEs compromised bacterial growth, with 16/19 T3SEs reducing bacterial growth in both *P. syringae* genetic backgrounds, and only three T3SEs displaying statistically significant strain-specific reductions despite being expressed in both strains (Figure S1): AvrPto1m and HopT1b reduced bacterial growth solely when delivered from PmaES4326, while HopI1k only reduced PtoDC3000 growth (Figure S1; Figure 2, Figure S4). While there were only three qualitative ETI differences, significantly more quantitative differences in ETI intensity were observed between strains, which correlated with the strain-specific results obtained from our symptom-based primary screen (Figure 2). These results highlight that strain-specific differences in ETI potential are predominantly quantitative rather than qualitative.

**Figure 2.**
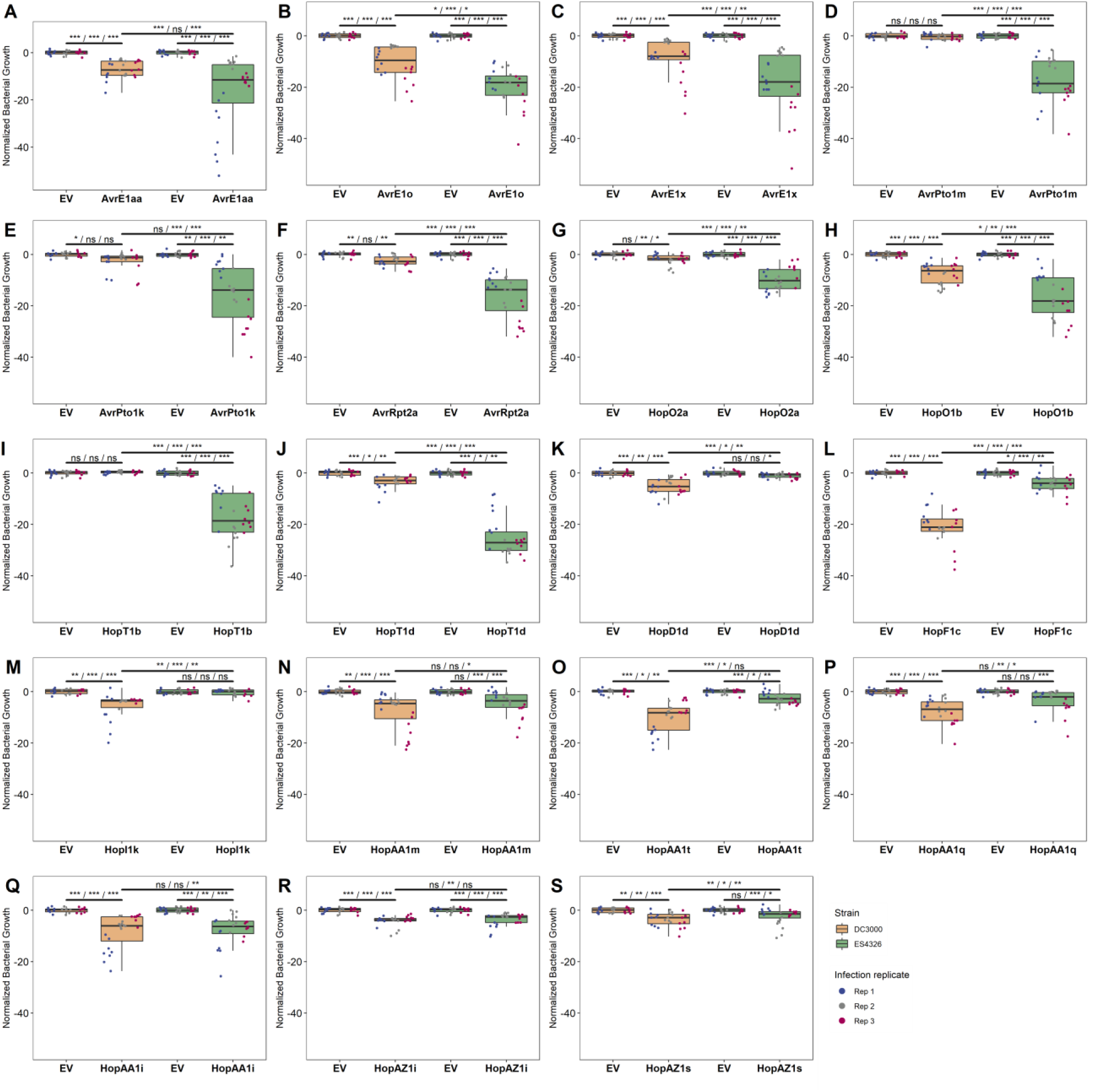
Strain-specific ETI differences are predominantly quantitative. Z-score normalized bacterial growth data for all 19 T3SE alleles that display differential ETI phenotypes when delivered from either **(A-J)** PmaES4326 or **(K-S)** PtoDC3000. Asterisks indicate treatments determined to be statistically significantly different based on pairwise T-tests (* P < 0.5, ** P < 0.1, *** P < 0.01). Spray inoculation growth assays were performed 3 days post infection. Raw growth assay data is presented in Figure S4.

### NLR-requirements and HR-induction of new ETI-eliciting T3SEs

We tested the new ETI-eliciting alleles uncovered in this study to determine if the ETI response was dependent on the same NLR required by other alleles in the same family. We confirmed that AvrRpt2a requires RPS2, HopO2a and HopO1b require ZAR1 and ZRK3, and AvrE1x, AvrE1o, and AvrE1aa require CAR1 (Figure S5A-C) [1, 37, 42]. We screened T-DNA knockouts of eight NLRs canonically associated with *P. syringae* ETI responses in an attempt to identify NLRs associated with AvrPto1 and HopT1 ETI (using AvrPto1m and HopT1b as representative alleles), but were unable to identify any candidates (Figure S5D-E) [43]. We also determined the ability of new ETI-eliciting T3SE alleles to induce a hypersensitive response (HR) and found that neither AvrPto1m nor HopT1b elicit HR on *A. thaliana* Col-0 when delivered from either PmaES4326 or the effectorless derivative of PtoDC3000 D36E (Figure S6A-B) [44]. Thus, the number of HR-independent ETI responses is further increased to 13/21 T3SE families recognized by *A. thaliana* Col-0 [1]. The three PmaES4326 ETI-eliciting alleles AvrRpt2a, HopO1b, and HopO2a elicit a weak ETI in PtoDC3000. These alleles are a part of previously characterized ETI-eliciting T3SE families that are associated with HR. When delivered from PtoDC3000, all three newly identified ETI-eliciting alleles trigger HR, although less consistently than their previously characterized homologs (Figure S6C-D).

### Distribution of the ETI load across the P. syringae phylogeny

Mapping the presence of newly identified ETI-eliciting T3SEs onto the *P. syringae* core genome phylogeny reveals 90 additional putative cases in which strains harbor T3SE orthologs that may be recognized by *A. thaliana* Col-0 (Figure 3A). Given that 96.8% of *P. syringae* strains were previously found to carry putative ETI-elicitors, it is unsurprising that we identified only two new strains that have ETI potential on *A. thaliana* Col-0, bringing the total to 97.2% (480/494). However, the number of strains carrying multiple ETI-eliciting T3SEs increased from 70.7% (349/494) to 74.3% (367/494), shifting the mean distribution of ETI-eliciting T3SEs per strain from 2.31 to 2.58, with the maximum number of putative ETI-elicitors carried by a single strain increasing from six to eight (Figure 3B). Notably, the novel ETI-eliciting alleles from AvrPto1 and HopT1 (only identified in the PmaES4326 screen) are both found predominantly in phylogroup 1 (Figure 3A). Given that PtoDC3000 is a phylogroup 1 strain, this suggests that members of this phylogroup may have developed effective methods to overcome the AvrPto1 and HopT1 immune responses.

**Figure 3.**
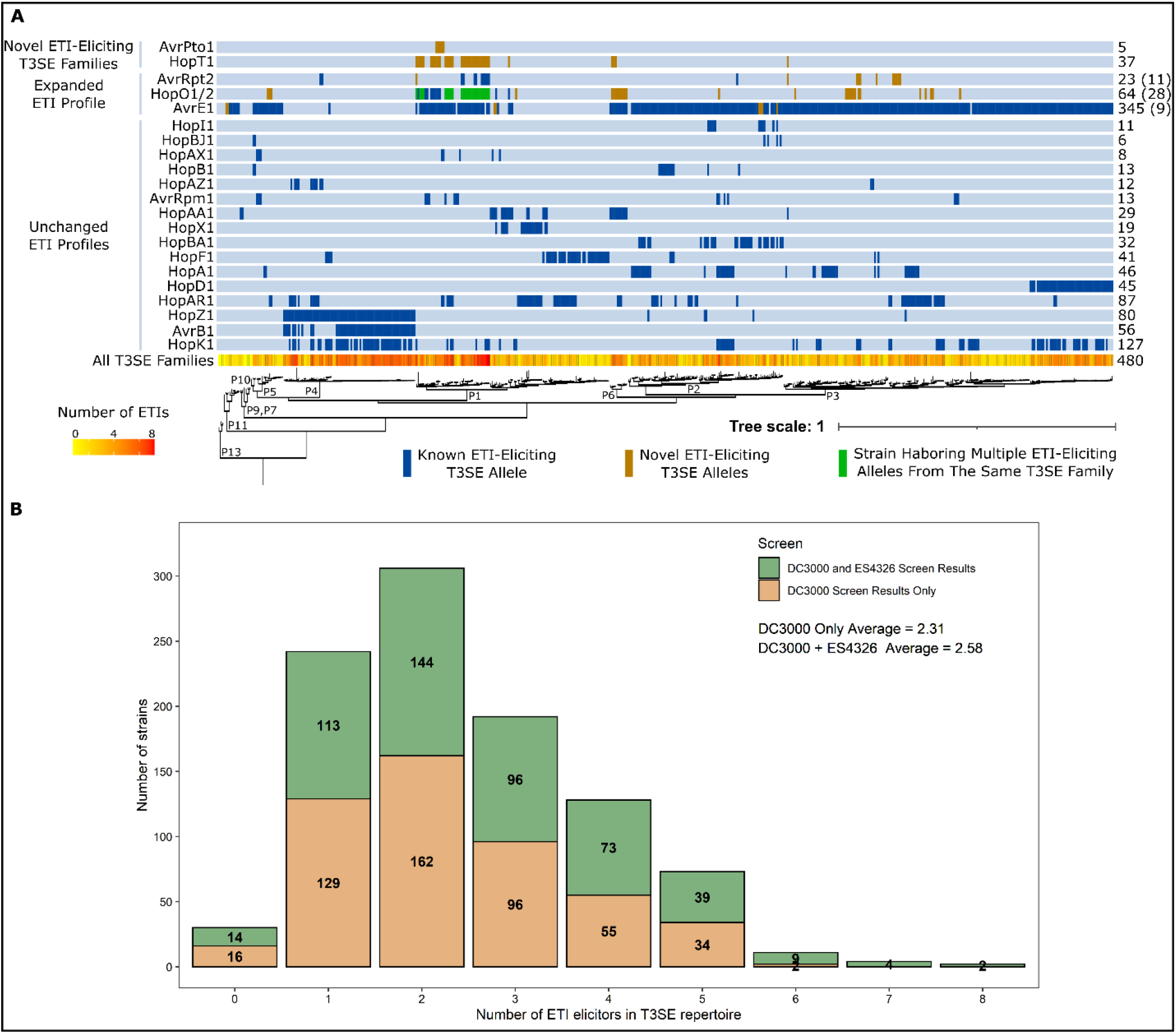
The *A. thaliana* ETI load of *P. syringae*. **(A)** Predicted orthologs of ETI elicitors carried by each of the 494 *P. syringae* strains used to generate the PsyTEC collection mapped onto a previously generated *P. syringae* core genome phylogeny with phylogroup designations indicated by P1-P13 [1, 39]. Putatively ETI-eliciting homologs are color coded as follows: blue - previously elucidated ETI-eliciting homolog; orange - newly identified putatively ETI-eliciting homologs originating from the PmaES4326 PsyTEC screen; and green - strain harboring previously and newly identified putatively ETI-eliciting homologs from the same T3SE family. Numbers on the right indicate the total number of strains that harbor putatively ETI-eliciting alleles from each T3SE family. For T3SE families with expanded ETI profiles, the number in parenthesis indicates the number of new strains harboring putatively ETI-eliciting T3SE alleles within that T3SE family. The total number of ETI-eliciting T3SE alleles carried by each strain is also mapped onto the phylogeny, ranging from 0 (yellow) to 8 (red). **(B)** Distribution of the number of putative ETI-eliciting T3SE alleles carried by each of the 494 *P. syringae* strains included in this study when using results from only the PtoDC3000 screen or from both the PtoDC300 and PmaES4326 screens [1].

### Multiple T3SEs from PtoDC3000 specifically suppress ETI

We hypothesized that PtoDC3000 carries one or more T3SE(s) that suppressed the AvrPto1m- and HopT1b-associated ETIs, thereby giving rise to the strain-specific ETI patterns observed. We therefore tested whether T3SEs native to PtoDC3000’s repertoire could suppress AvrPto1m or HopT1b ETI responses when expressed in PmaES4326. To this end, we developed a two-vector system to express ETI-elicitors (from pBBR1-MCS2 [45], kanamycin-resistant) alongside putative ETI-suppressors (from pUCP20TC, tetracycline-resistant, see methods) in the same strain, which we could then test for ETI using our disease-based screening assay [1, 38]. We validated this system using the well-studied AvrRpm1 / AvrRpt2 suppression system [26, 27]. Our assay showed that AvrRpt2 suppressed AvrRpm1 ETI in a *rps2* mutant background when delivered from either PtoDC3000 or PmaES4326 (Figure S7A-B). Further, both effectors were appropriately expressed from their respective vectors in PtoDC3000 and PmaES4326 (Figure S7C-D). Using this two-vector system, we generated a collection of PmaES4326 strains expressing AvrPto1m or HopT1b in combination with the 36 well-expressed T3SE alleles from the PtoDC3000 repertoire [44] (Table S3). We confirmed that the presence of an empty ‘suppressor’ vector did not impact the corresponding ETI responses by quantifying disease symptom progression (Figure S8).

We performed spray inoculation of *A. thaliana* Col-0 with our ETI-elicitor / putative suppressor strains and quantified macroscopic disease symptoms as described above for our ETI screen. We normalized percent yellow tissue values within the bounds of our controls, resulting in disease scores ranging between 0.0 (positive ETI control) and 1.0 (no-ETI control). Three PtoDC3000 T3SE alleles provided partial ETI suppression phenotypes: HopQ1a suppressed AvrPto1m (45% disease score), and HopG1c and HopF1g suppressed HopT1b (39% and 57% disease score, respectively) (Figure 4A, Table S4). These increases in disease symptoms were not due to lack of ETI-elicitor expression (Figure S9).

**Figure 4.**
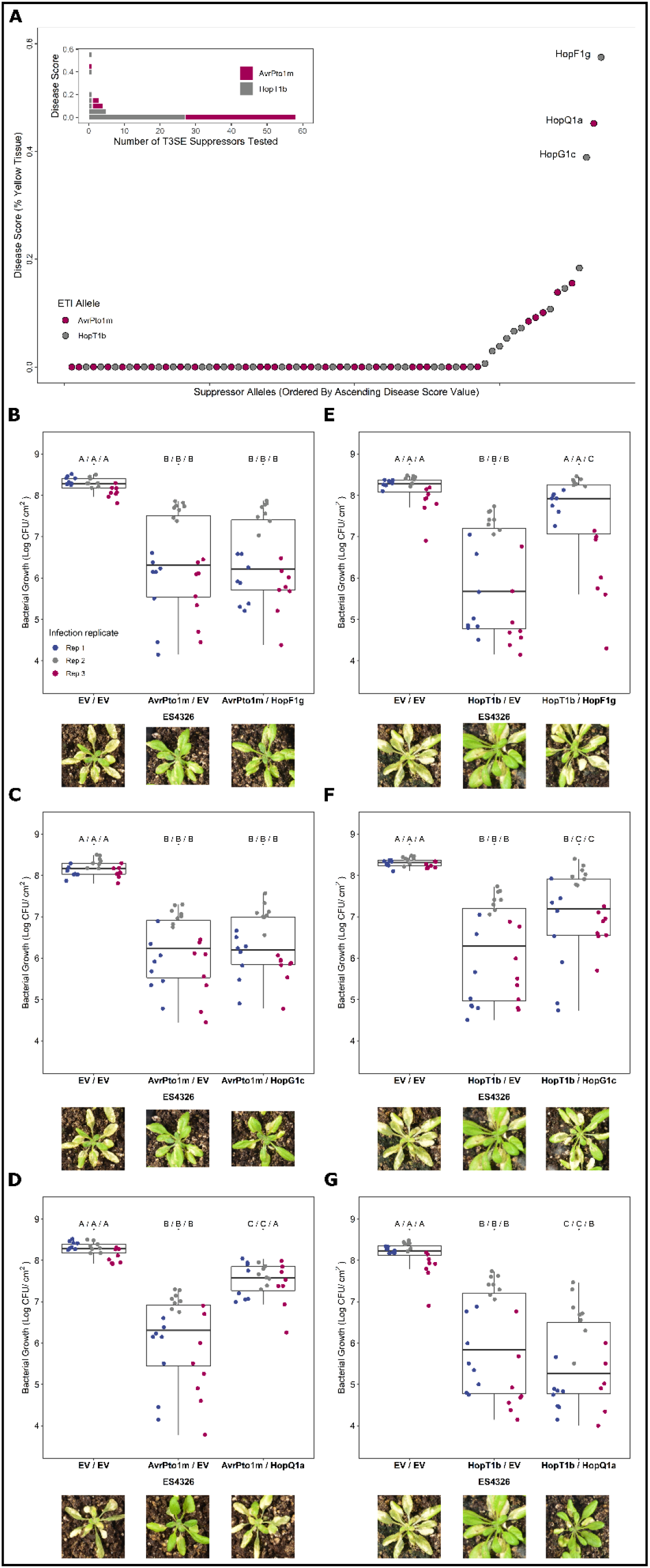
Identification of PtoDC3000 ETI suppressing effectors. **(A)** Plant disease scores obtained from strains used to identify suppressors of AvrPto1m- and HopT1b-associated ETIs. PmaES4326 strains harboring the ETI elicitor (on the MCS2 vector) and the PtoDC3000 candidate suppressor allele (on the pUCP20TC vector) were used for spray inoculation. Plant disease scores were determined by plotting percent chlorosis [38] within the bounds of the positive ETI control (MCS2::ETI pUCP20TC::EV, disease score = 0.0) and the negative ETI control (MCS2::EV pUCP20TC::EV, disease score = 1.0). Validation of identified ETI suppression phenotypes of **(B-D)** AvrPto1m and **(E-G)** HopT1b through bacterial growth assays. Letters represent statistically significant differences (Tukey’s HSD, P < 0.05). Plant images display representative visual phenotypes following spray-inoculation assays 5-7 days post infection.

We validated the observed ETI suppression phenotypes through bacterial growth assays, testing all three identified suppressor alleles (HopF1g, HopG1c and HopQ1a) on both AvrPto1m and HopT1b. Consistent with our screening results, we observed partial restorations in bacterial growth when either HopF1g or HopG1c was co-expressed with HopT1b, but not AvrPto1m, and when HopQ1a was co-expressed with AvrPto1m, but not HopT1b (Figure 4B-G). As observed with disease symptoms, no single suppressor restored empty-vector levels of bacterial growth in all three replicates, highlighting the partial contribution of these ETI suppression phenotypes. Overall, our screen identified three partial and specific suppressors of ETI: HopF1g and HopG1c suppressed HopT1b ETI, whereas HopQ1a suppressed AvrPto1m ETI.

We also assessed the ETI potential of AvrPto1m and HopT1b in PtoDC3000 lacking their associated ETI-supressing T3SEs using bacterial growth assays. Although we observed a weak reduction in bacterial growth using AvrPto1m in a subset of replicates in wild-type PtoDC3000, a greater reduction was observed (>10-fold) when delivered from the PtoDC3000 ΔHopQ1a background [44] (Figure 5A). Similarly, HopT1b only reduced bacterial growth in the ΔHopF1g [28] and ΔHopG1c PtoDC3000 strains [46] (Figure 5C, E). Complementation of these suppressor knockout PtoDC3000 strains with their corresponding T3SEs significantly increased bacterial growth in all cases (Figure 5B, D, F). Growth of the HopQ1a knock out strain was only partially restored by complementation, suggesting that additional factors, such relative T3SE expression levels, may contribute to the magnitude of ETI suppression. Overall, our results demonstrate that the native PtoDC3000 T3SEs HopQ1a, HopF1g and HopG1c can suppress the ETI responses of AvrPto1m or HopT1b.

**Figure 5.**
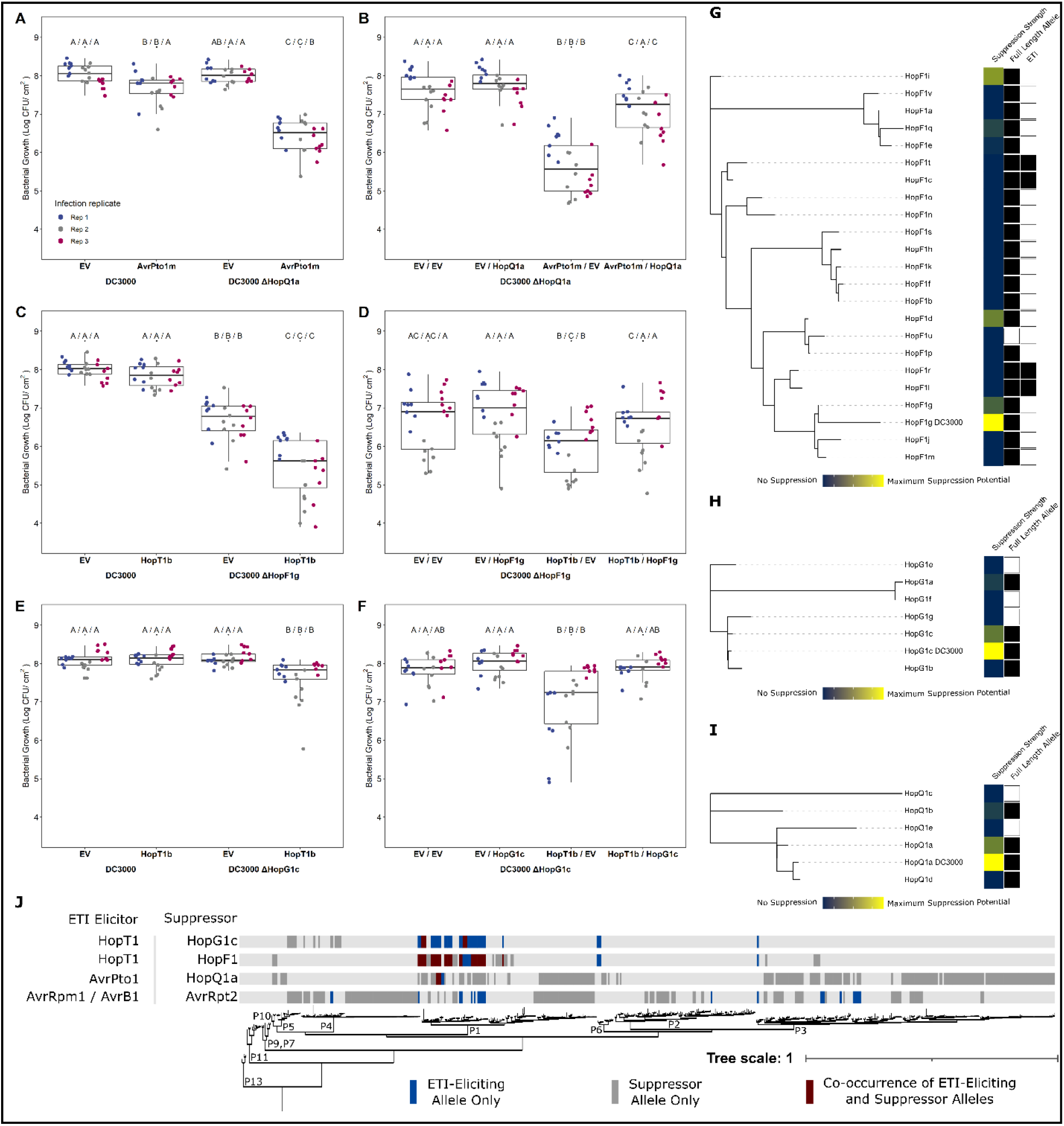
Modulation of ETI load in *P. syringae*. Comparative bacterial growth assays and subsequent complementation of **(A-B)** AvrPto1m in PtoDC3000 vs. PtoDC3000 ΔHopQ1a, **(C-D)** HopT1b in PtoDC3000 vs. PtoDC3000 ΔHopF1g, and **(E-F)** PtoDC3000 ΔHopG1c. Letters represent statistically significant differences (Tukey’s HSD, P < 0.05). Maximum likelihood PsyTEC representative allele T3SE family trees for **(G)** HopF1, **(H)** HopG1, and **(I)** HopQ1, including the PtoDC3000 allele. The first metadata column shows relative ETI suppression strength using the DC3000 allele as reference for maximum suppression (yellow to blue represent high to low suppression). The second metadata column identifies full length alleles, with filled squares representing alleles whose amino acid length is at least 75% the family average. The third metadata column show HopF1 ETI elicitors (filled squares). **(J)** Co-occurrence of ETI elicitor - suppressor pairs across the *P. syringae* core genome phylogeny [1, 39]. The presence of solely an ETI elicitor allele from an ETI elicitor - suppressor pair is denoted in blue, the presence of solely the suppressor is denoted in grey, and co-occurrence of both components of an ETI - suppressor pair is denoted in burgundy.

### Diversification of ETI suppression across T3SE families

We next investigated the extent to which ETI suppression is conserved across T3SE subfamilies. We used PsyTEC representatives of the HopF1 (23 alleles), HopG1 (7 alleles) and HopQ1 (6 alleles) T3SE subfamilies, co-expressed them with their corresponding ETI suppression target in PmaES4326, and determined a normalized disease score. Certain members of the HopF1 family are known to trigger ZAR1-dependent ETI [1, 47] and were thus tested for suppression on *zar1-1* plants. For all three T3SE families, the PtoDC3000 identical T3SE allele displayed the strongest suppression phenotype (Figure 5G-I). In the HopF1 family, the PsyTEC representatives of HopF1g, HopF1i, HopF1d and HopF1q displayed moderate suppression of the HopT1b ETI (Figure 5G). Across the HopG1 and HopQ1 PsyTEC representatives, one member of each family partially suppressed the HopT1b or AvrPto1m ETI. Across all three suppressing T3SE families, only full-length alleles were capable of ETI suppression. These results show that ETI suppression phenotypes are not broadly distributed across T3SE families and display allele specificity.

## DISCUSSION

In this study we investigated how pathogen genotype, in particular the T3SE repertoire, influences the ETI potential on a particular host. We built on the ETI landscape that we previously established using PtoDC3000 on the model plant *Arabidopsis thaliana* (Col-0), by repeating the PsyTEC ETI screen with PmaES4326 as the pathogenic background. This screen uncovered ten novel ETI-eliciting T3SE families, including two new T3SE families that had not been identified as elicitors when expressed from PtoDC3000 (Figure 1). Summing together the ETI landscapes identified using both PtoDC3000 and PmaES4326, the prominence of ETI in the *A. thaliana* - *P. syringae* pathosystem is further strengthened, with 69/529 (13.0%) PsyTEC effectors representing 21/70 (30.0%) T3SE families capable of eliciting ETI on one accession of *A. thaliana*. Importantly, 27.5% (19/69) of these ETI eliciting alleles displayed pathogen strain-specific differences highlighting the important contribution of pathogen genotypes in modulating ETI responses. When the ETI load inferred from both screens are mapped onto the *P. syringae* phylogeny together, 97.2% (480/494) of *P. syringae* strains possess an ortholog of a putative ETI-eliciting T3SE allele. 74.3% of strains possess more than one putative ETI-elicitor, with up to eight ETI responses being predicted for certain strains of *P. syringae* (Figure 3). These findings further cement ETI as a pervasive component of broad-spectrum resistance against *P. syringae* in *A. thaliana* and a significant selective load that must be mitigated by *P. syringae* strains.

### ETI load mitigation by P. syringae

The phytopathogen *P. syringae* strain PtoDC3000 has one of the best characterized effector repertoires of any plant pathogen. These effectors successfully promote virulence in *A. thaliana* and tomato, despite possessing at least one T3SE that can trigger ETI in *A. thaliana*. Namely, our original PsyTEC screen [1] identified AvrE1 as an ETI-elicitor in *A. thaliana* despite it being native to PtoDC3000. AvrE1 is required to promote water soaking in a functionally redundant manner with HopM1 [15]. However, expression of plasmid-borne AvrE1 in PtoDC3000 triggers CAR1-mediated ETI in *A. thaliana* [1]. We hypothesize that PtoDC3000 must carefully regulate the expression of AvrE1 in order to realize virulence benefits but avoid activating ETI. In another example of ETI avoidance, PtoDC3000 possesses a heavily truncated allele of HopO1c that is not recognized by the ZAR1 NLR in *A. thaliana*, unlike the full-length HopO1c PsyTEC allele from PsyUSA007 [1]. In this study we have shown that metaeffector interactions can also suppress ETI responses in PtoDC3000, which possesses at least 3 T3SEs that can dampen the ETI responses elicited by AvrPto1 and HopT1. When combined with previous studies, at least 33 PtoDC3000 T3SEs can suppress ETI and/or ETI-associated HR to various extents across multiple species, emphasizing ETI-suppression as a major function of its T3SE repertoire [19, 28, 30, 44, 48, 49].

We find multiple cases of co-occurrence of ETI elicitor - ETI suppressor pairs within *P. syringae* strains, particularly in phylogroup 1 (Figure 5J). This is not the case for the classic example of ETI suppression of AvrRpm1 or AvrB1 by AvrRpt2, where no instances of co-occurrence are seen among the 494 strains we have used for our studies (Figure 5J). We do not rule out the possibilities that strains harboring both AvrRpm1/AvrB1 and AvrRpt2 exist but have not been sampled, or that horizontal gene transfer could result in such strains arising in the future. However, we highlight that the co-occurrence of ETI elicitor - suppressor pairs in multiple *P. syringae* strains identified in this study suggests that these T3SE interactions are likely biologically significant. Supporting the biological relevance of our ETI-suppression events, we found that AvrPto1m and HopT1b are primarily native to phylogroup 1 strains of *P. syringae*, alongside PtoDC3000 (Figure 3A). Given that these immune responses were only identified when the effectors were expressed in PmaES4326, it is likely that effectors particular to phylogroup 1 strains, such as PtoDC3000, can naturally mask these immune responses. Lending further support to this idea, we found multiple instances of co-occurrence between AvrPto1m or HopT1b and their suppressors (Figure 5J). The natural co-occurrence of elicitors and suppressors not only suggests that these suppression events have applicability to other *P. syringae* strains beyond the model PtoDC3000, but also that certain inferences of ETI based on our PsyTEC-wide screens are likely to be at least partially suppressed in a natural context. These examples are likely to be significant underestimates of the actual scale of ETI suppression that exists across the diverse *P. syringae* effector pool.

### The quantitative nature of ETI across the *P. syringae* pan-genome

Although the ETI-elicitation profiles of PtoDC3000 and PmaES4326 were largely overlapping, the 19/69 differential elicitors highlight that pathogen genotype has a major impact on ETI elicitation (Figure 1, Figure 2). Remarkably, these differences in ETI elicitation between pathogen backgrounds were mostly quantitative, rather than qualitative. Only three of 19 differential ETI responses were completely abolished in one background (Figure 2). Indeed, the other 16 ETI responses could be described as ETI elicitors when expressed in either background, though their impact on bacterial virulence is weak enough in one background to not be captured consistently by our primary screening method (Figure 1, Figure 2, Figure S2). Quantitative variation in ETI can therefore skew towards extreme weakness – to the point where traditional macroscopic phenotypes for monitoring ETI, such as plant health (Figure S2) and the hypersensitive response (Figure S6), are unable to reliably capture it. It is only when performing sensitive measurements of bacterial load that some weaker ETI responses can be validated. For example, HopT1d was not captured in our PtoDC3000 screen and does not elicit HR, however, sensitive bacterial load measurements confirm that it weakly but significantly impacts PtoDC3000’s fitness on Col-0 (Figure 2, Figure S6) [1]. While we still classified ETI-elicitors based on the disease scores obtained in both screens for consistency, we undoubtedly missed some weaker ETI responses that didn’t pass our disease score threshold. Further, T3SE alleles that were classified as ETI-elicitors in both screens may likely also display quantitative variation based on the pathogenic background, although their immune phenotypes remained strong enough to be captured in both primary screens.

### ETI suppression capability is not widely conserved in T3SE families

Using the effector repertoire of PtoDC3000, we found three effector alleles capable of suppressing either AvrPto1m-triggered or HopT1b-triggered immunity (HopF1g, HopG1c, and HopQ1a). Interestingly, these ETI-suppression phenotypes were not broadly distributed across these effector families: only one member from each of the HopG1 and HopQ1 families, and four members from the highly diverse HopF1 family showed suppression phenotypes (Figure 5G-I). The lack of widespread ETI suppression in these effector families undoubtedly reflects functional divergence, particularly given that effector orthologs can have amino acid sequence identity lower than 50% [20]. The specificity of ETI-suppression within families, as well as the fact that no effectors appeared to have a general capacity to suppress immunity (Figure 4B-G), suggest that these suppression events entail the disruption of specific immune pathways similar to what was observed for AvrRpm1/AvrRpt2. Alternatively, direct interactions between specific effectors, as observed for *L. pneumophila*, may modulate effector activities and corresponding ETI-eliciting capacities [21].

The *P. syringae* ETI-suppression activities that we identified did not completely restore virulence, and therefore are partial or quantitative. The lack of complete (i.e., qualitative) ETI suppression was not an experimental artifact since our two-vector system was able to recapitulate the complete suppression of AvrRpm1 ETI by AvrRpt2 (Figure S7). This accords with much of the previous literature on metaeffector interactions in *P. syringae*, where partial ETI suppression is more common than complete suppression [18, 30, 44, 49–51]. Although HopF1 has previously been reported to suppress ETI in *A. thaliana* [19, 28, 48], this is the first report of *A. thaliana* ETI-suppression for HopG1 and HopQ1. Outside of *A. thaliana*, all three of these PtoDC3000 T3SEs have been reported to suppress or delay ETI to various extents. In *Nicotiana benthamiana,* HopQ1 and HopF1 can block HopAD1-associated HR [52], while HopG1 can partially suppress or delay multiple HRs [30, 52]. Further, HopF1 carried by Pta11528 can suppress HR on *N. benthamiana* [49], HopF1 carried by Pph1449B can suppress HR on bean [53] and HopQ1 native to *P. syringae* pv. *actinidiae* strains can suppress multiple HRs on *N. benthamiana* [52]. In addition to ETI-suppression, HopF1 dampens PTI by targeting several PTI components, including RIN4 [19], FLS2 [54], and MAP kinase kinases [55, 56], while HopG1 modulates the structure of the actin cytoskeleton to promote disease [46]. While the specifics of HopQ1 function are unknown, the effector does bind to 14-3-3 proteins [57], which have been implicated in effective ETI signaling [58–61]. It is possible that HopQ1 functions primarily in *A. thaliana* as a suppressor of ETI, though certainly more functional characterization of this effector is necessary to make that conclusion. Thus, ETI suppression, rather than being the primary role of these effectors, is likely a by-product of their functional promiscuity.

## ACKNOWLEDGMENTS

We thank members of the Desveaux and Guttman labs for helpful feedback throughout this project. We would also like to thank Brad Day (Michigan State University) for providing the PtoDC3000 ΔHopG1 strain and Alan Collmer (Cornell University) for providing the PtoDC3000 ΔHopQ1 strain. This project was supported by Natural Sciences and Engineering Research Council of Canada (NSERC) postgraduate awards (AM and BL), NSERC Discovery grants (DSG and DD) and a Canada Research Chair in Plant-Microbe Systems Biology (DD).

## AUTHOR CONTRIBUTIONS

AM, BL, DSG and DD designed experiments. AM and BL generated and validated the PmaES4326 PsyTEC collection and performed the primary ETI screen. AM, BL, CBM and FL performed growth assay validations. AM and BL generated and validated the suppression strains and performed the suppression screen. AM, BL and CBM validated suppression phenotypes and performed HRs. AM, BL, DSG and DD wrote the manuscript.

## SUPPLEMENTAL DATA

**Figure S1. Compiled immunoblots depicting expression of PsyTEC in PmaES4326**

**Figure S2. Compiled images of primary ETI screening of PsyTEC in PmaES4326**

**Table S1. Summary protein expression of the PsyTEC in PmaES4326.**

**Table S2. Summary primary ETI screening data of PsyTEC in PmaES4326.**

**Table S3. PtoDC3000 T3SE alleles used for the ETI suppression screen**

**Table S4. Summary ETI suppression screen data.**

## SUPPLEMENTAL FIGURES

**(Multi-page figure included in supplemental data)**

**Figure S1.** Compiled immunoblots depicting expression of PsyTEC in PmaES4326 Immunoblots against the HA tag of each of the 529 PsyTEC representative alleles in PmaES4326 following overnight growth in *hrp*-inducing minimal media. An empty-vector (EV) and positive expression control (HopZ1a) is included in each immunoblot. A ponceau staining of the membrane is presented under each immunoblot to display equal loading. An individual asterisk (*) indicates T3SEs for which we could not detect expression. Two asterisks (**) indicates that expression was detected, but at a different size than expected. NR indicates that this sample is not relevant to this study. Numbers to the left of the immunoblots describe the molecular weight of the ladder, in kDa.

**(Multi-page figure included in supplemental data)**

**Figure S2. Compiled images of primary ETI screening of PsyTEC in PmaES4326**

Images of *A. thaliana* Col-0 plants spray inoculated with PmaES4326 harboring representative PsyTEC T3SE alleles 5 - 10 days post infection that were used to determine percent chlorosis and disease score metrics presented in Figure 1. T3SE allele names are presented above the images, while the associated disease score for each treatment is described below. A positive ETI control (“P” PmaES4326 harboring HopZ1a, with a disease score of 0) and a negative ETI control (“N” PmaES4326 harboring an empty vector, with a disease score of 1) were included on each flat. T3SE descriptions are color coded based on their ETI classification using a disease score threshold of 0.45, where blue - ETI positive T3SE and red - ETI negative T3SE. Shaded- out plant columns are not relevant to this study.

---

**Figure S3.**
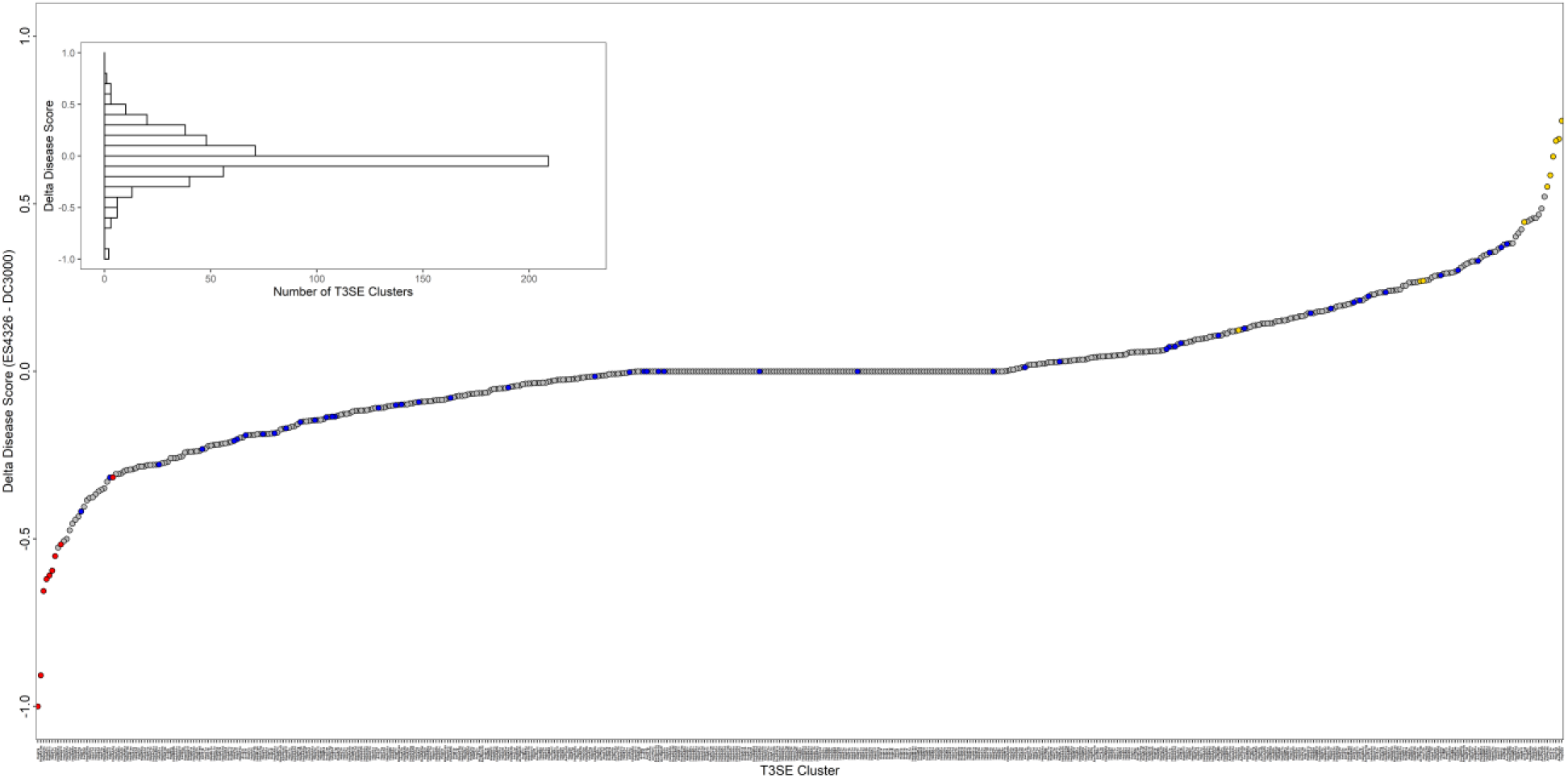
Delta disease score between PmaES4326 and PtoDC3000 ETI screens for all PsyTEC alleles. Comparison of disease scores obtained for all 529 PsyTEC T3SE alleles between the PmaES4326 and PtoDC3000 PsyTEC ETI screens [1]. Delta disease score is calculated as (Disease score obtained in PmaES4326 ETI screen) - (Disease score obtained in the PtoDC3000 ETI screen) for each individual T3SE allele. A histogram presenting the distribution of obtained delta disease score values is also included. Individual T3SE allele datapoints are color coded as follows: grey - no ETI when delivered from either genetic background, blue - ETI when delivered from either PtoDC3000 or PmaES4326, red - ETI observed only when delivered from PtoDC3000 and yellow - ETI observed only when delivered from PmaES4326.

**Figure S4.**
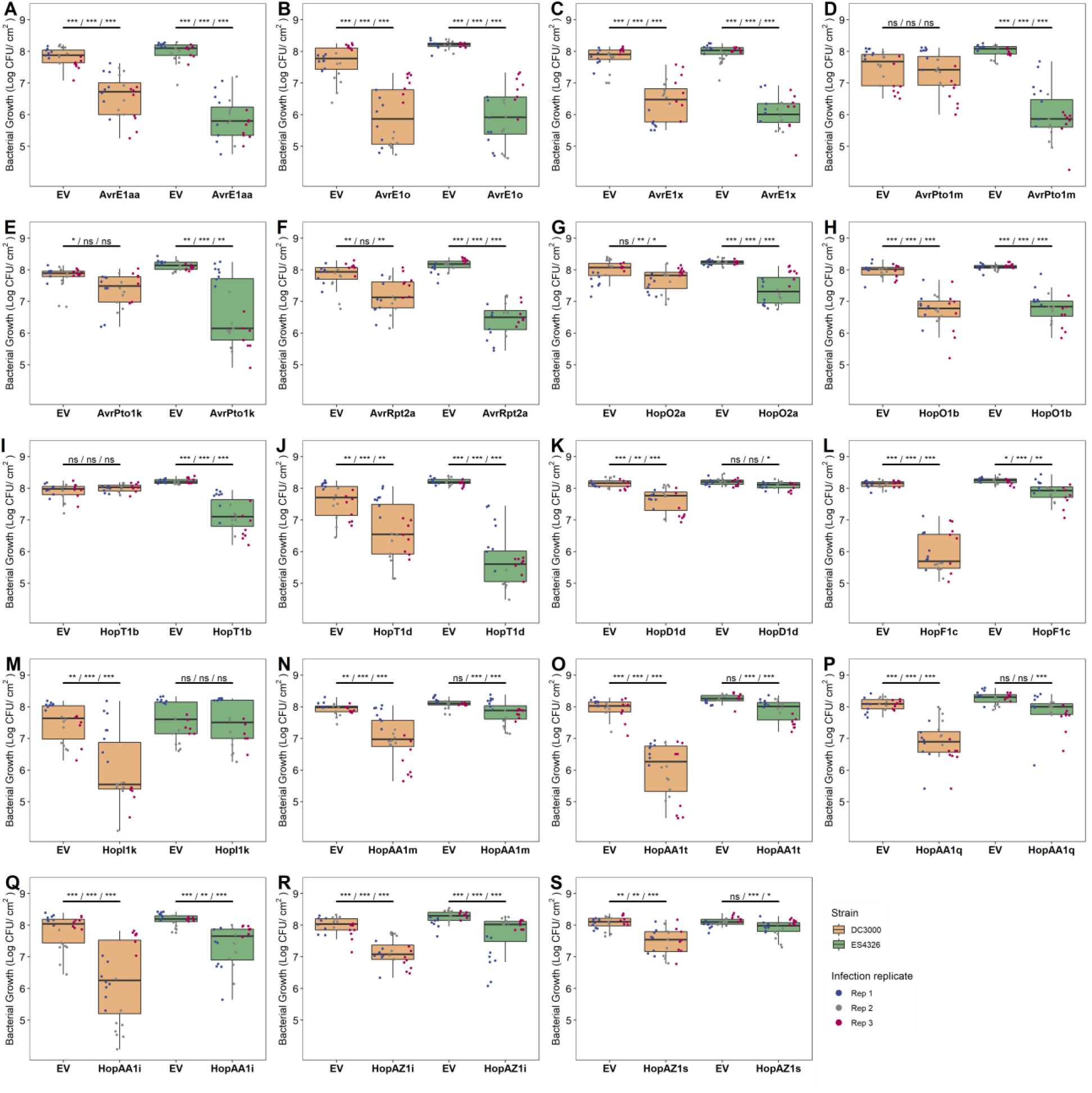
Comparative bacterial growth assays of 19 T3SEs in PtoDC3000 and PmaES4326 on *A. thaliana* Col-0. **(A-J)** Bacterial growth data for all 19 T3SE alleles that displayed differential ETI phenotypes when delivered from either PmaES4326 or **(K-S)** PtoDC3000. Asterisks indicate treatments determined to be statistically significantly different based on pairwise T-tests (* P < 0.5, ** P < 0.1, *** P < 0.01). Growth assays were performed 3 days post infection.

**Figure S5.**
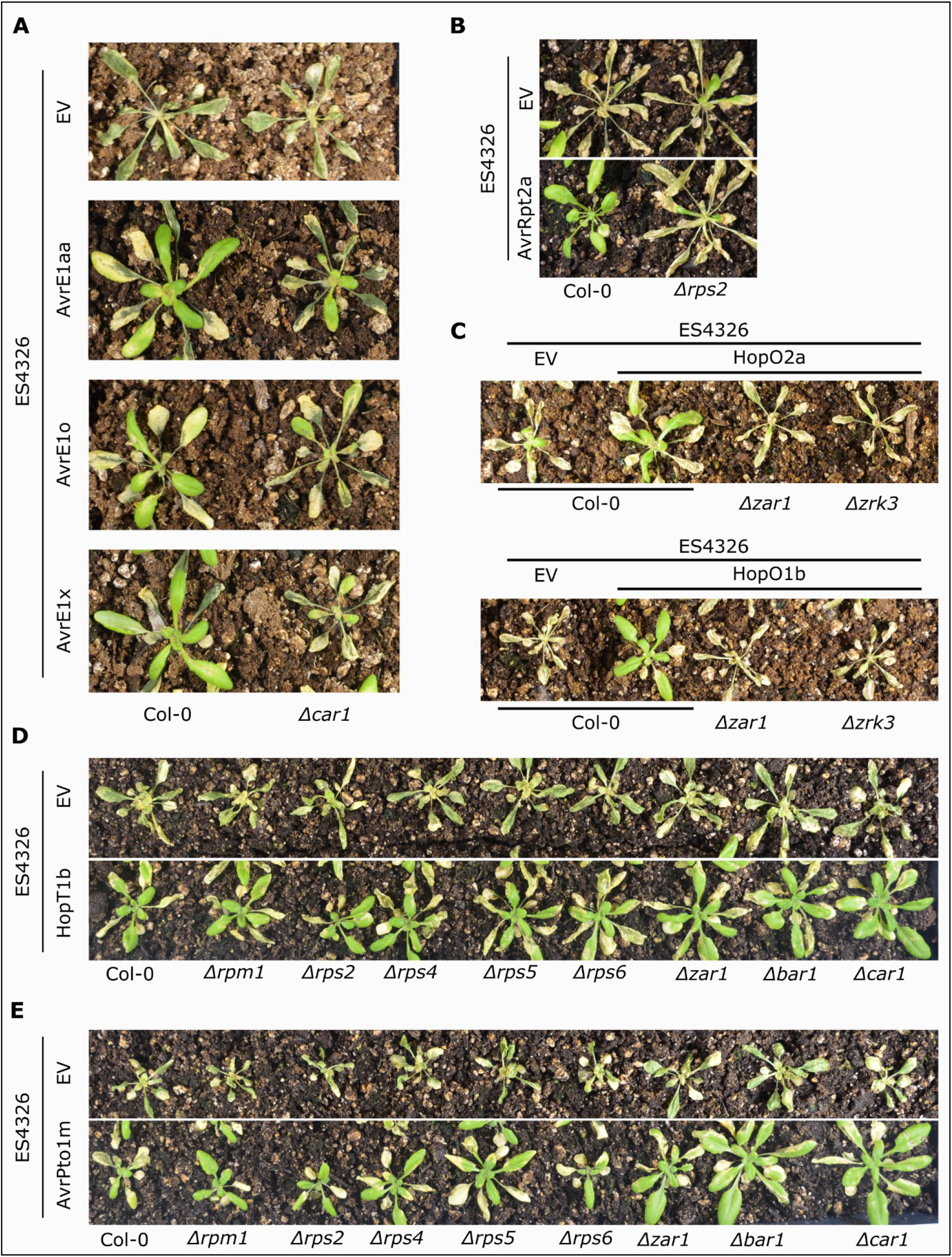
Validation of R gene requirements for newly identified ETI-eliciting T3SE alleles. Host genetic requirements of newly identified ETI-eliciting T3SE families belonging to T3SE families with previously characterized host genetic components was confirmed, when delivered from PmaES4326. **(A)** AvrE1x, AvrE1o and AvrE1aa were spray-inoculated on Col-0 and *Δcar1* (*car1-1*), **(B)** AvrRpt2a was spray inoculated onto Col-0 and Δrps2 and **(C)** HopO2a and HopO1b were spray-inoculated on Col-0, Δzar1 (*zar1-1*) and Δzrk3 (*zrk3-1*). One allele from each newly identified ETI-eliciting T3SE family, **(D)** AvrPto1m and **(E)** HopT1b, was spray inoculated onto a collection of 8 knockout lines representing each characterized NLR mediating *P. syringae* T3SE ETIs in *A. thaliana*.

**Figure S6.**
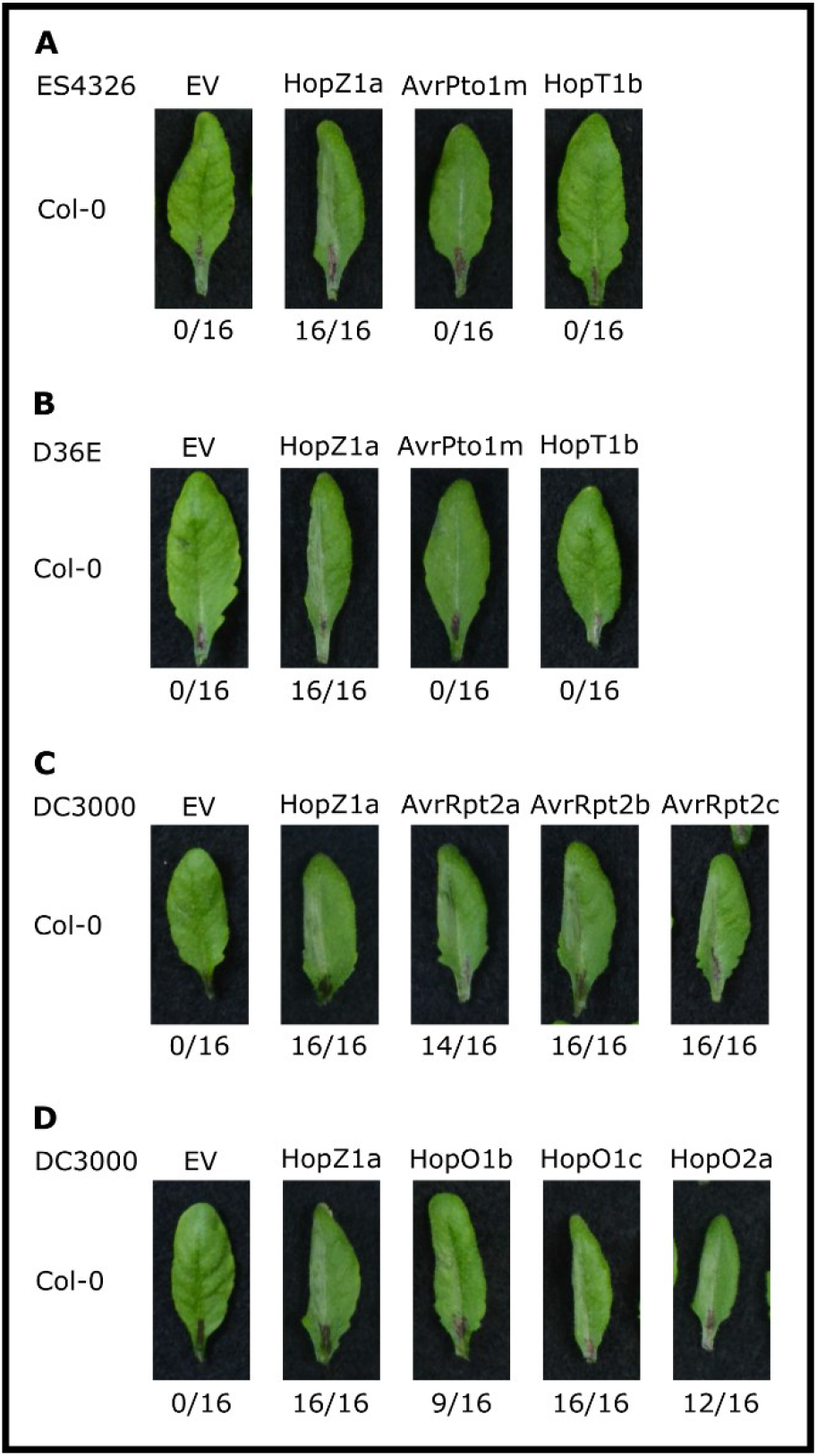
Validation of HR phenotypes for newly identified ETI-eliciting T3SE alleles. Hypersensitive response (HR) assays delivering an empty-vector (EV), HopZ1a, AvrPto1m and HopT1b from **(A)** PmaES4326 and from **(B)** the effectorless PtoDC3000 derivative D36E into the left side of *A. thaliana* Col-0 leaves. HR assays delivering all ETI-eliciting alleles from the **(C)** AvrRpt2 and the **(D)** HopO1/2 T3SE families, in conjunction with an empty-vector (EV) and HopZ1a control, delivered from PtoDC3000 into the left side of *A. thaliana* Col-0 leaves. Numbers below the representative leaf images represent the total number of observed macroscopic tissue collapses observed out of 16. These experiments were repeated 3 times with similar results.

**Figure S7.**
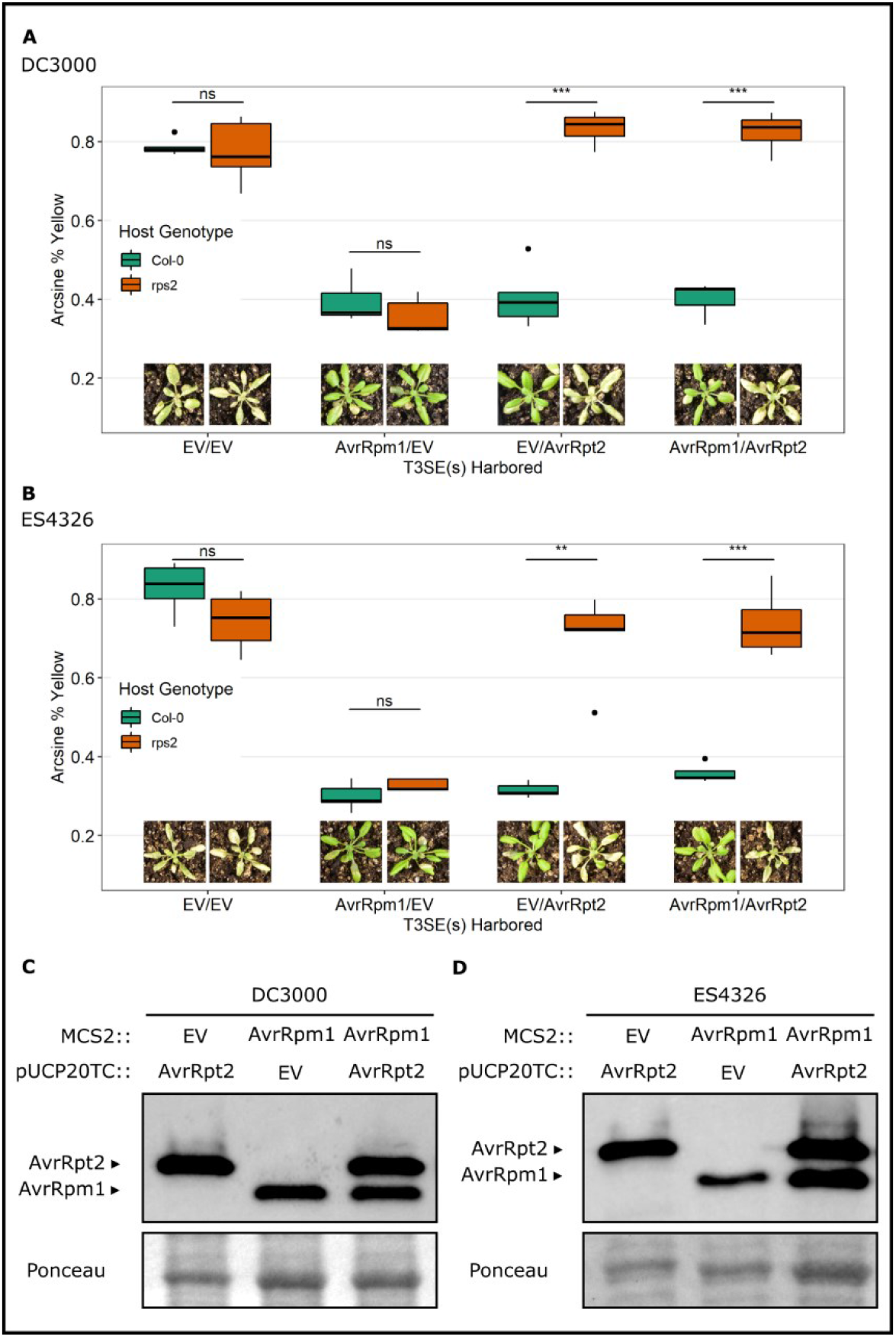
Validation of disease symptom-based suppression system. Validation that disease symptom quantification effectively captures the well characterized AvrRpm1/AvrRpt2 ETI suppression example using our two-vector system in both **(A)** PtoDC3000 and **(B)** PmaES4326. Asterisks indicate treatments determined to be statistically significantly different based on pairwise T-tests (* P < 0.5), whereas ns indicated no significant differences. Representative plant images for each treatment are included. Validation of protein accumulation through immunoblotting for all T3SEs in **(C)** PtoDC3000 and **(D)** PmaES4326 is presented, with associated Ponceau load controls below.

**Figure S8.**
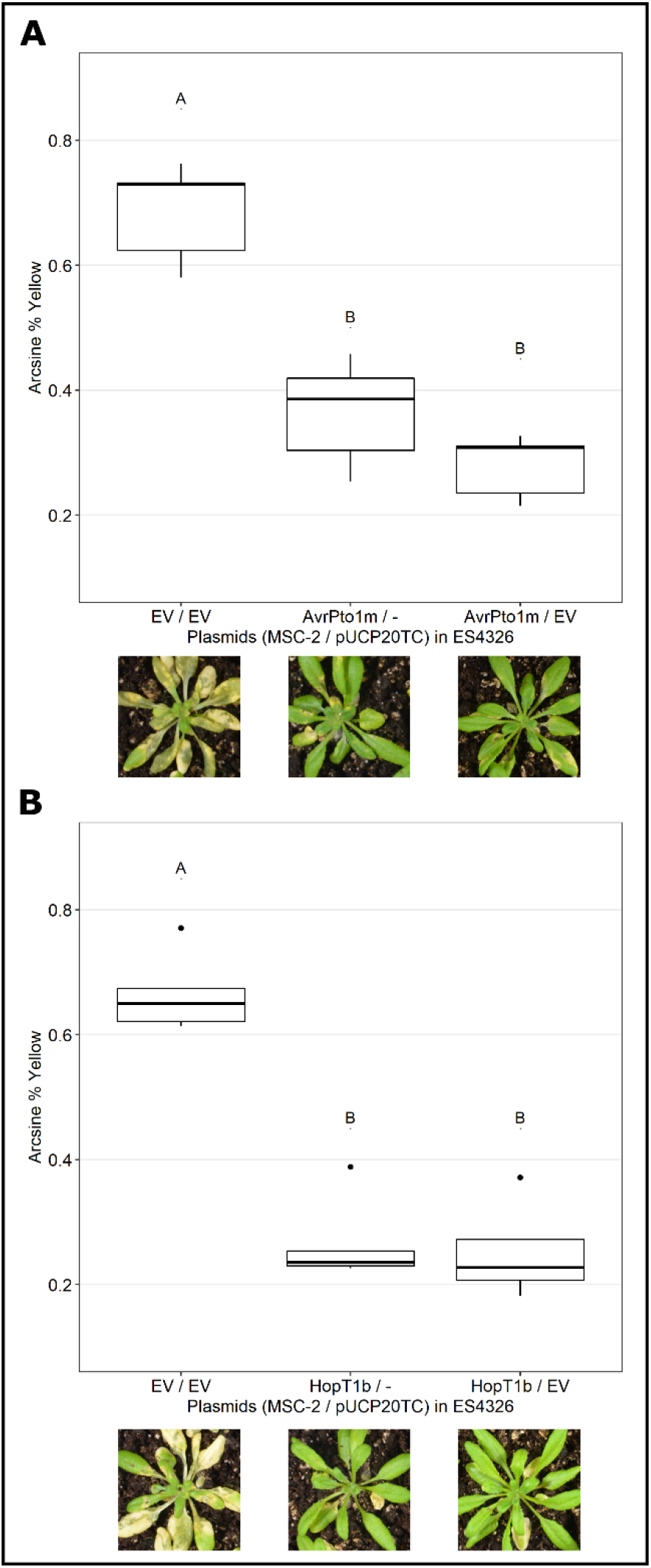
Validation of ETI phenotype in strains carrying multiple vectors. Visual disease symptom quantification through PIDIQ [38] validating similar ETI phenotypes for strains carrying only the ETI elicitor on MSC2 and those carrying an ETI elicitor on MCS2 and an empty vector pUCP20TC. Representative plant images for each treatment are included.

**Figure S9.**
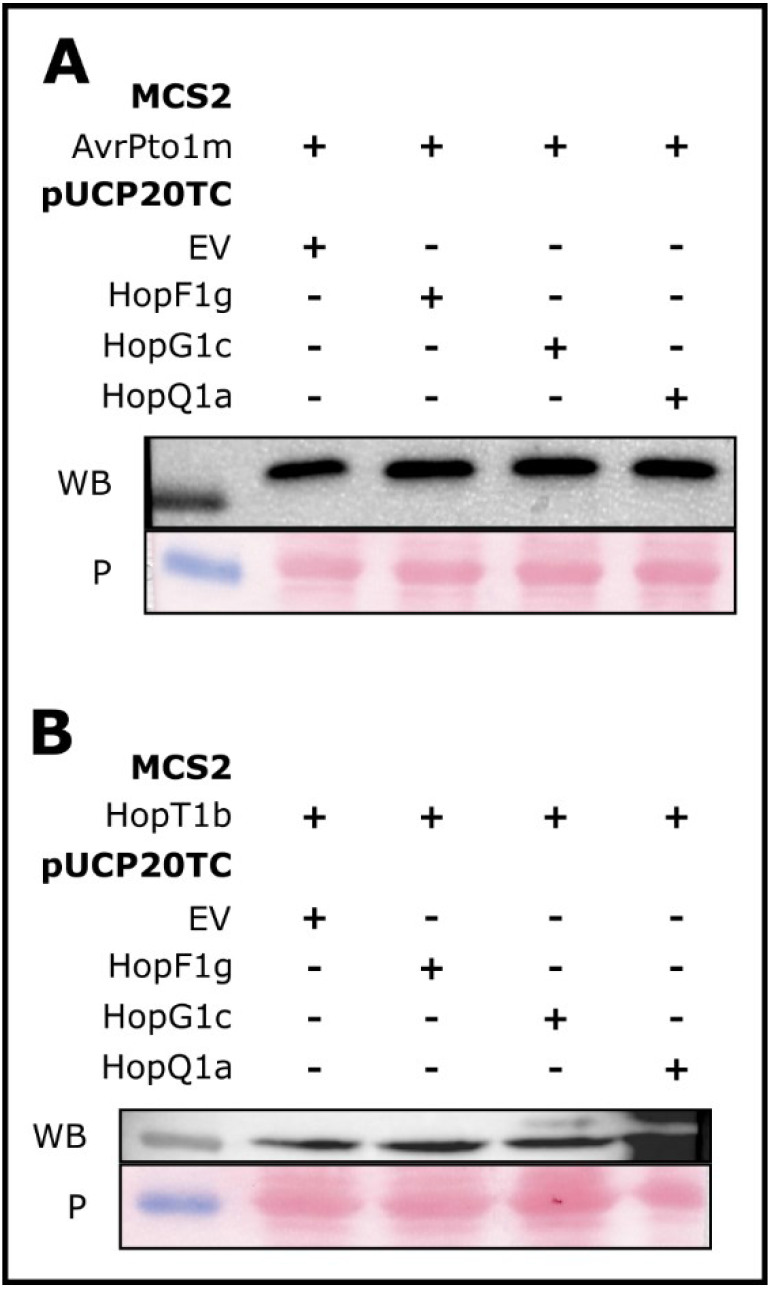
Validation of ETI-elicitor expression in suppression strains. Immunoblots confirming the minimal media-induced expression of **(A)** AvrPto1m and **(B)** HopT1b in the suppression strains harboring HopF1g, HopG1c, HopQ1a or an empty vector pUCP20TC plasmid. WB depicts the immunoblot against the ETI elicitor of interest probing for the fused HA tag; P depicts the associated Ponceau stained membrane confirming equal protein loading.

