## Supplementary figures and images for "Metaeffector interactions modulate the type III effector-triggered immunity load of *Pseudomonas syringae*"

### Fig S1

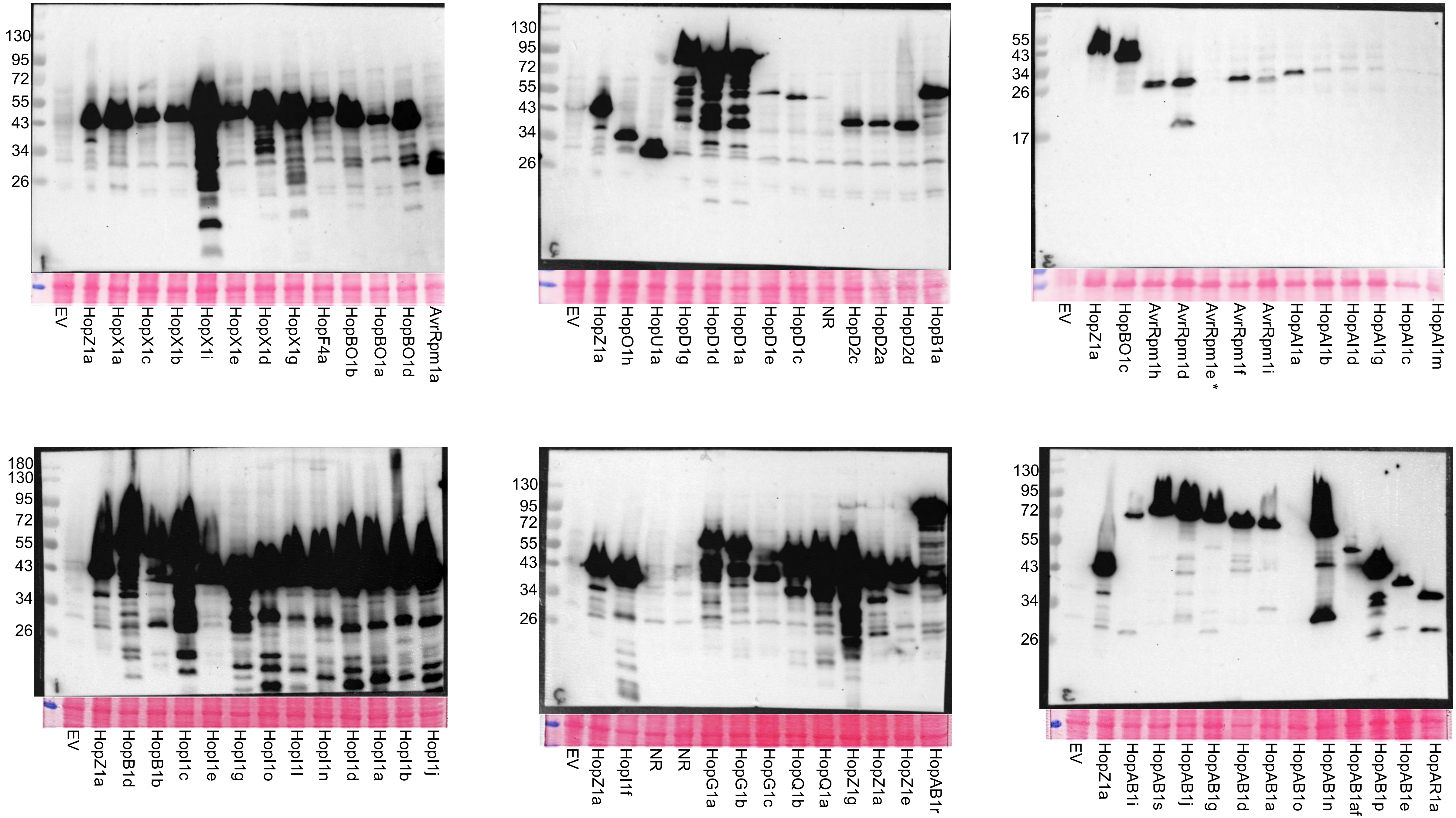

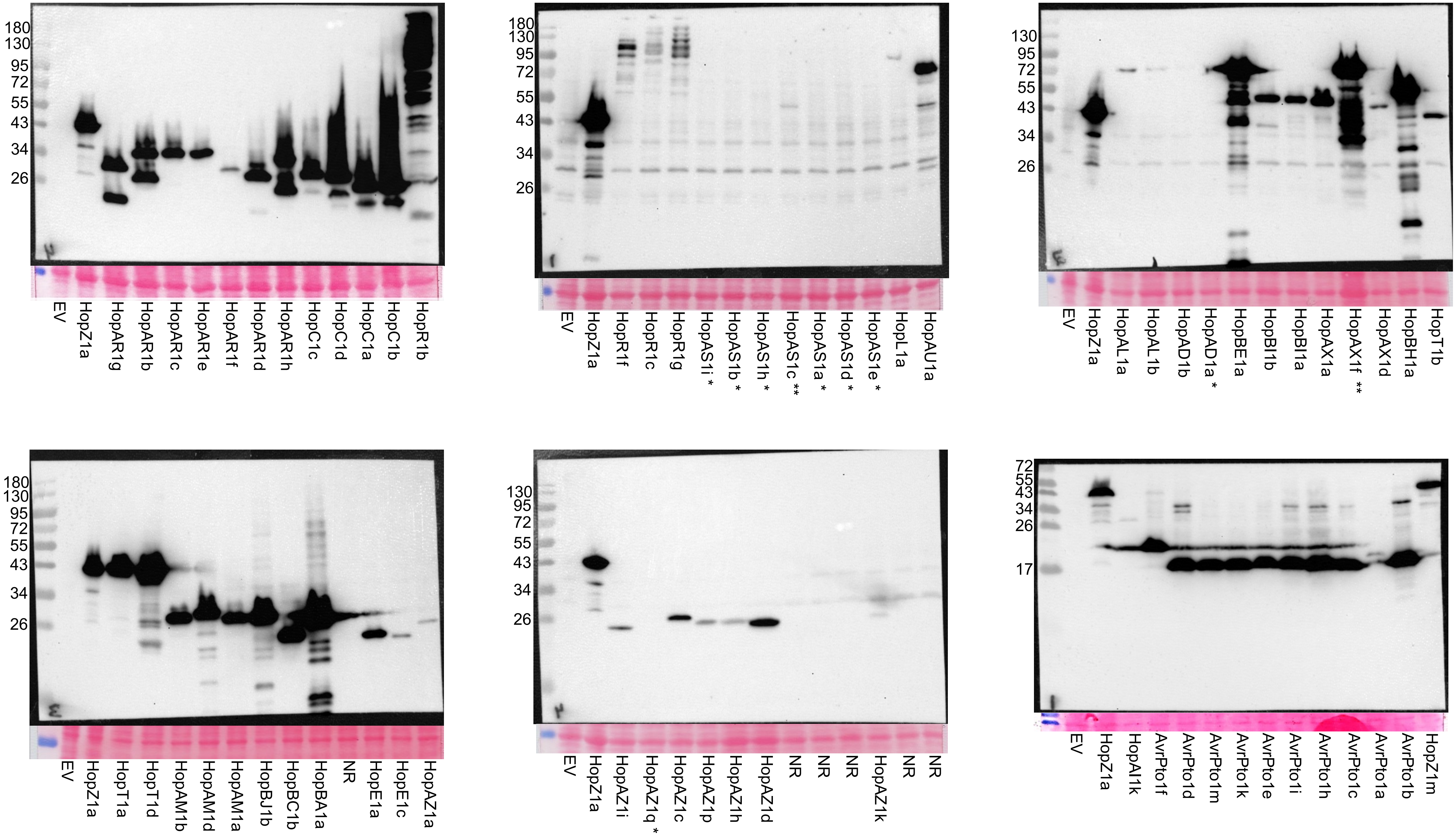

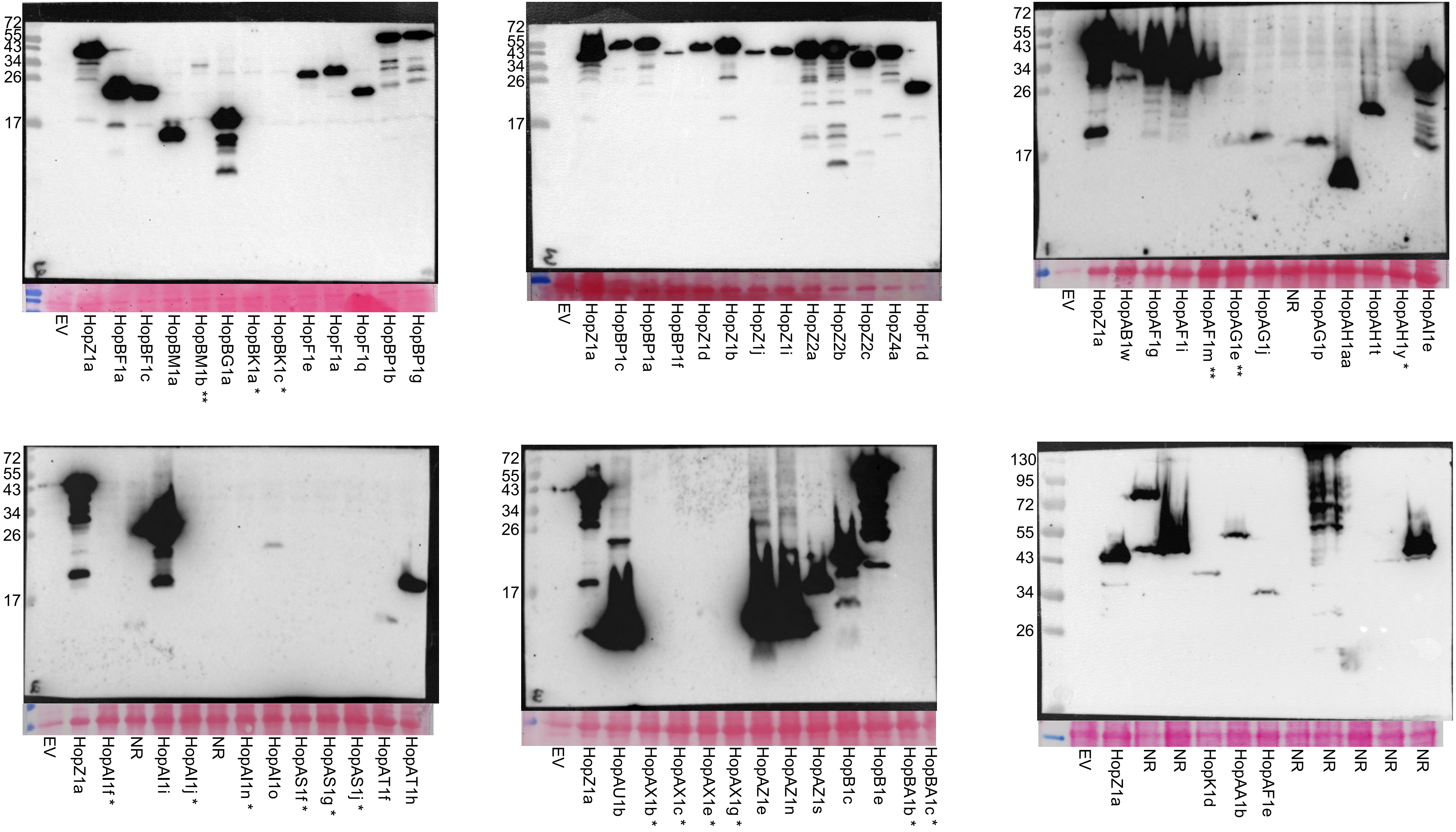

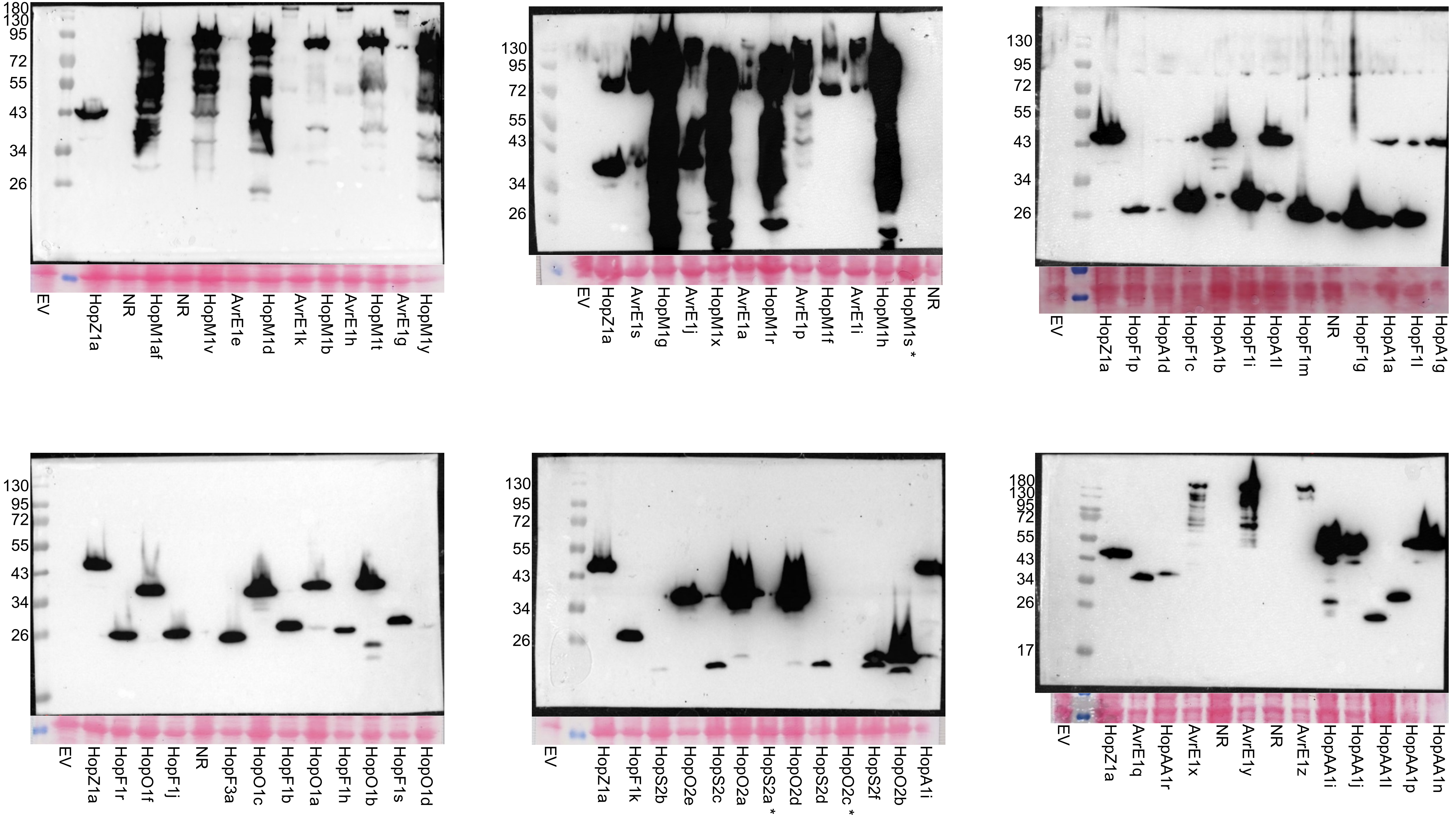

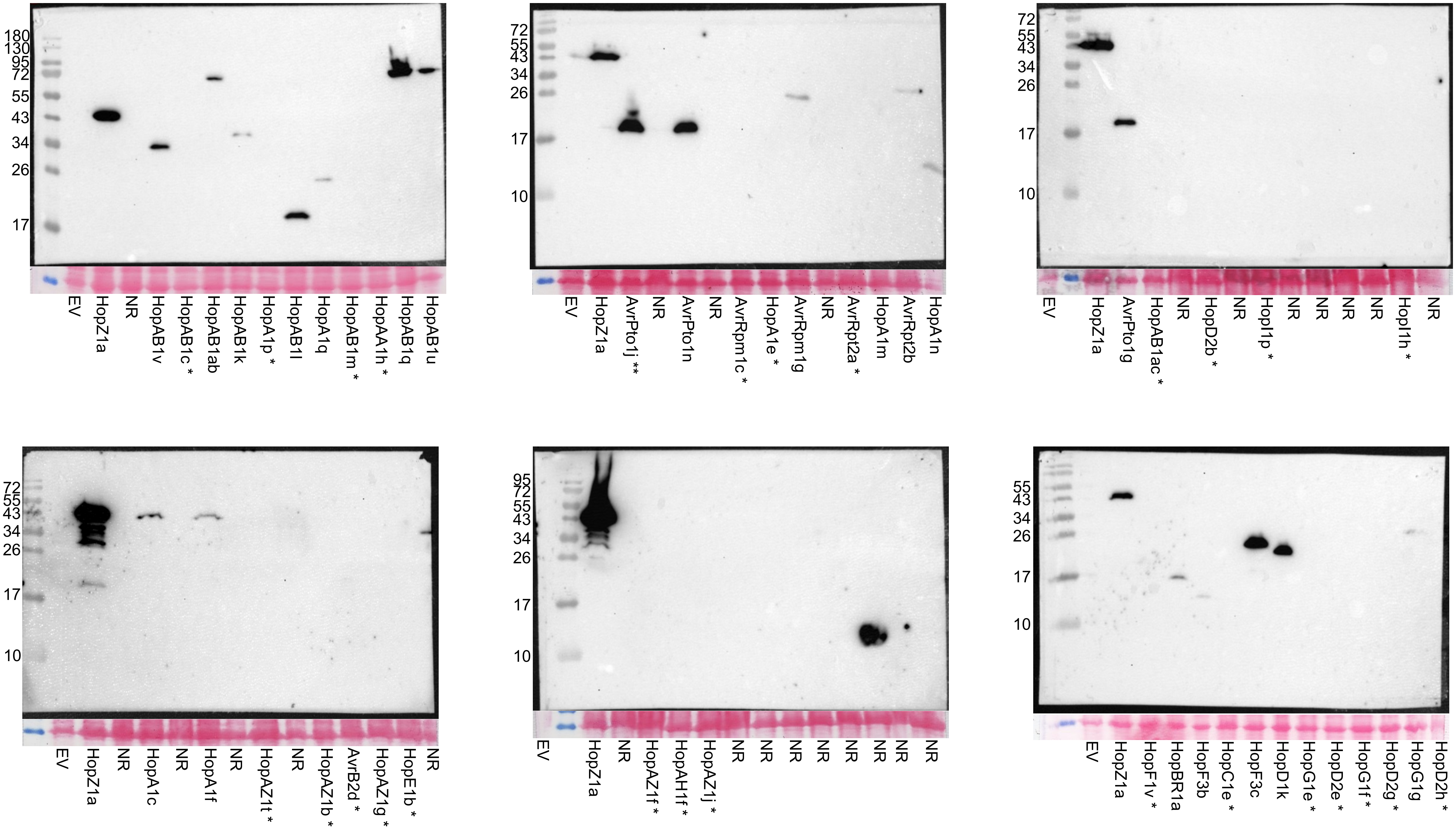

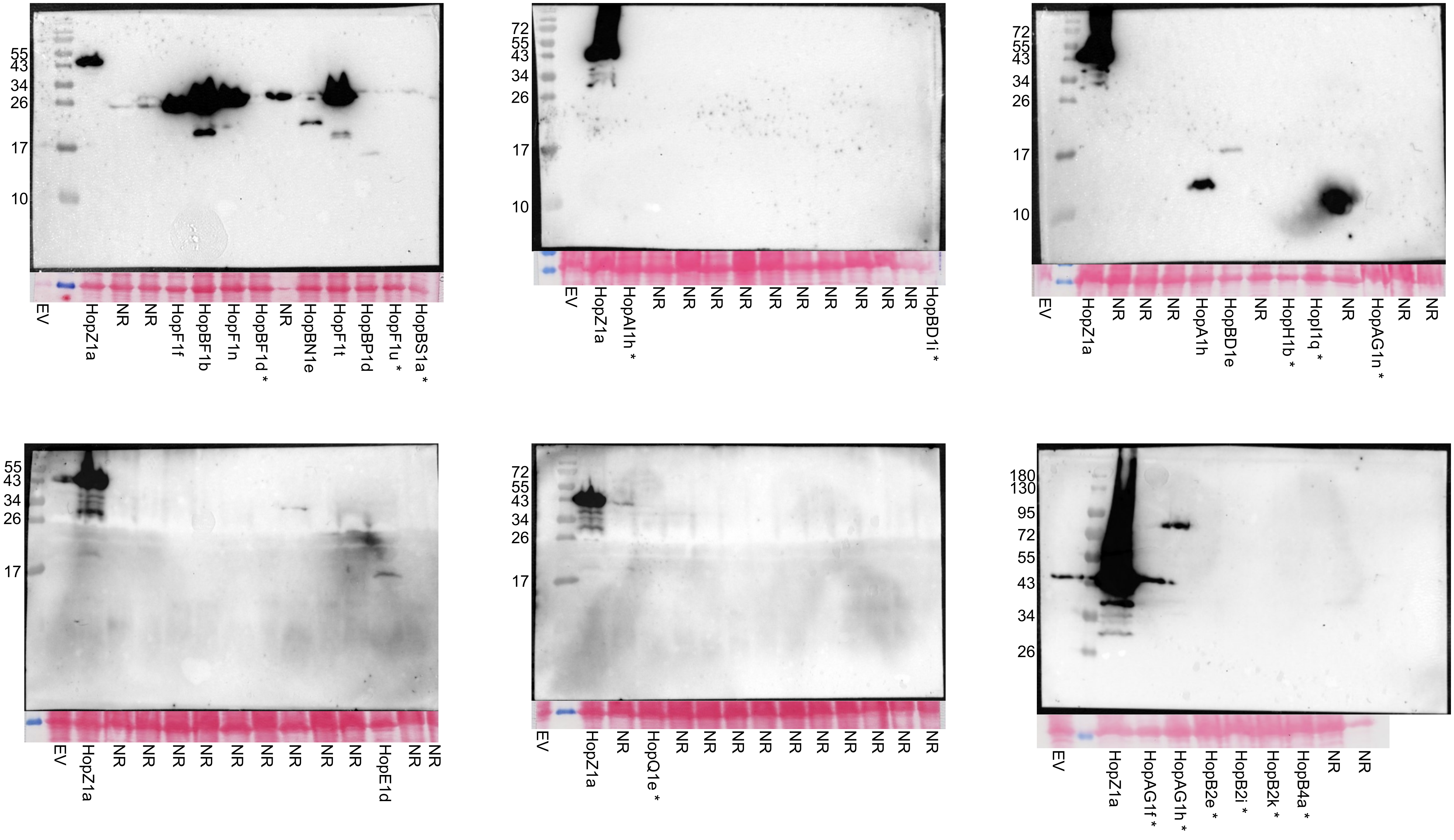

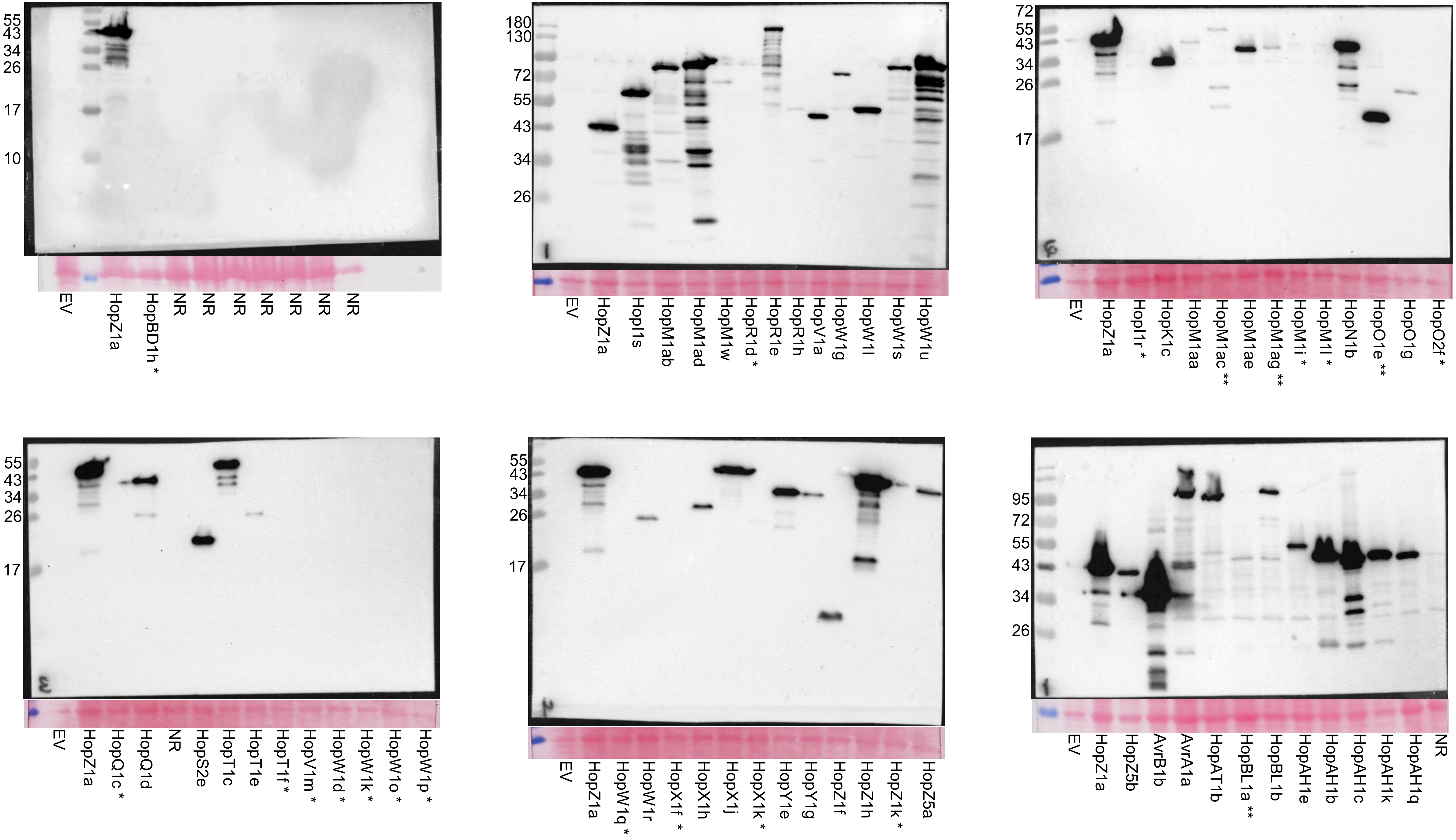

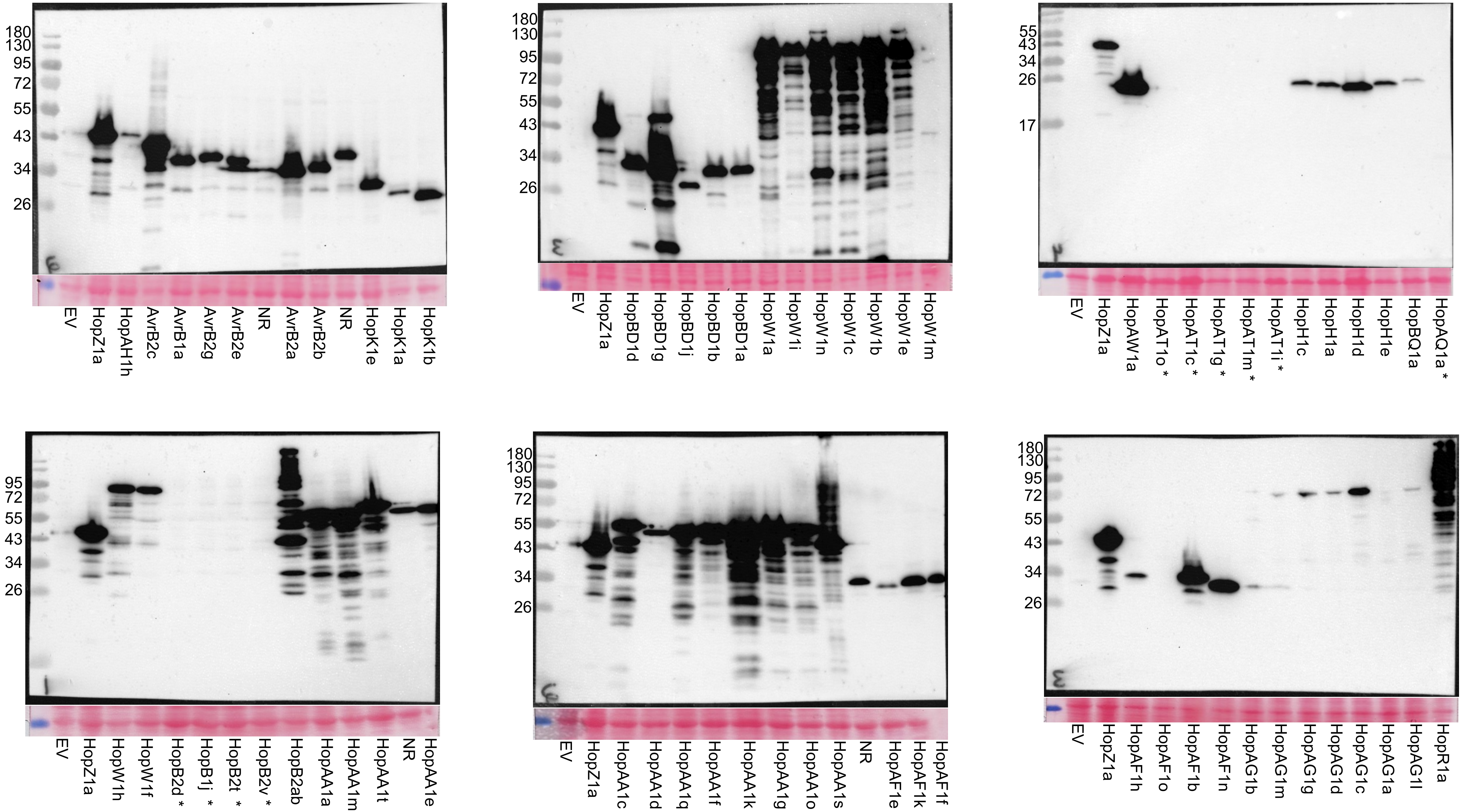

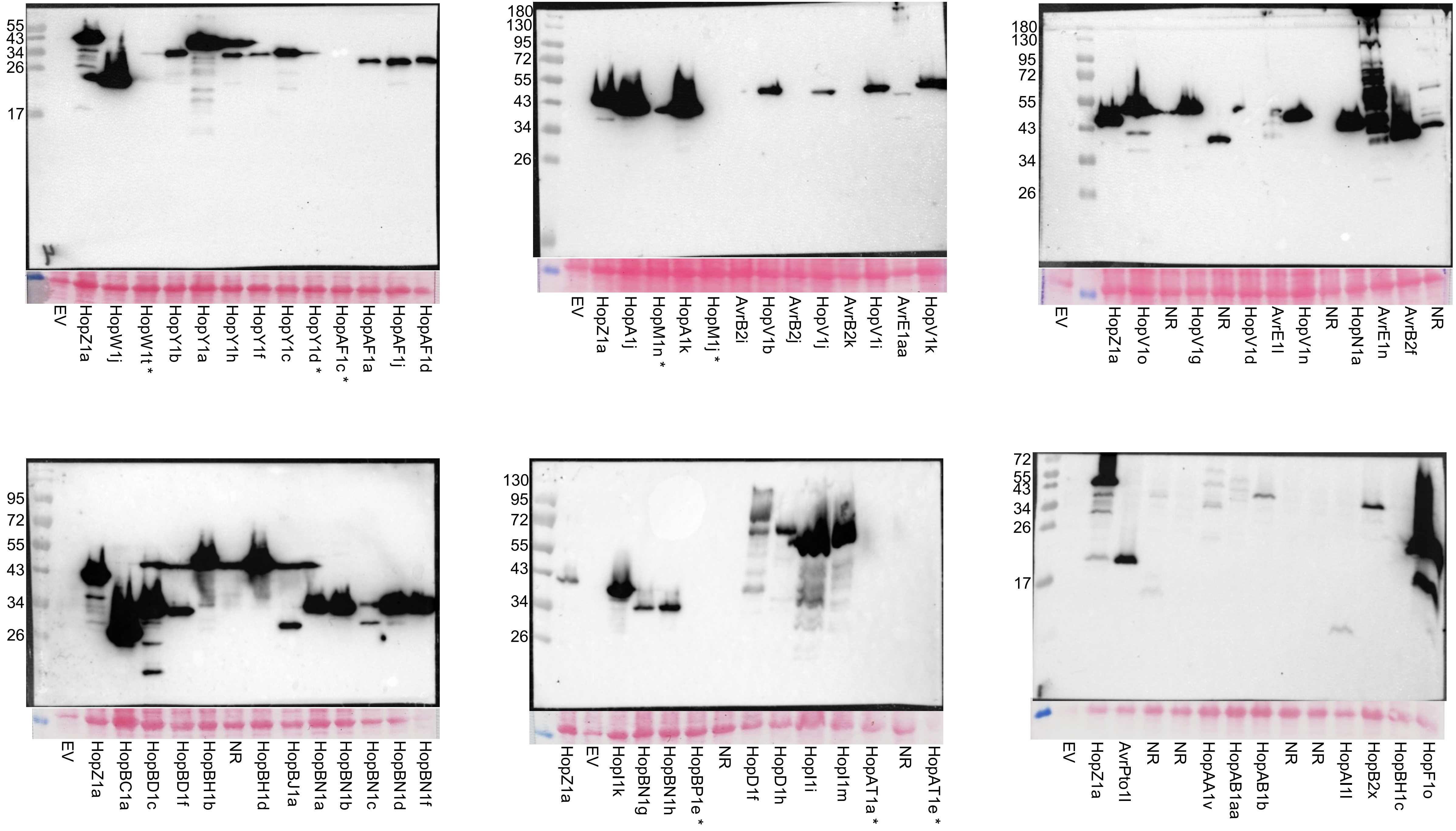

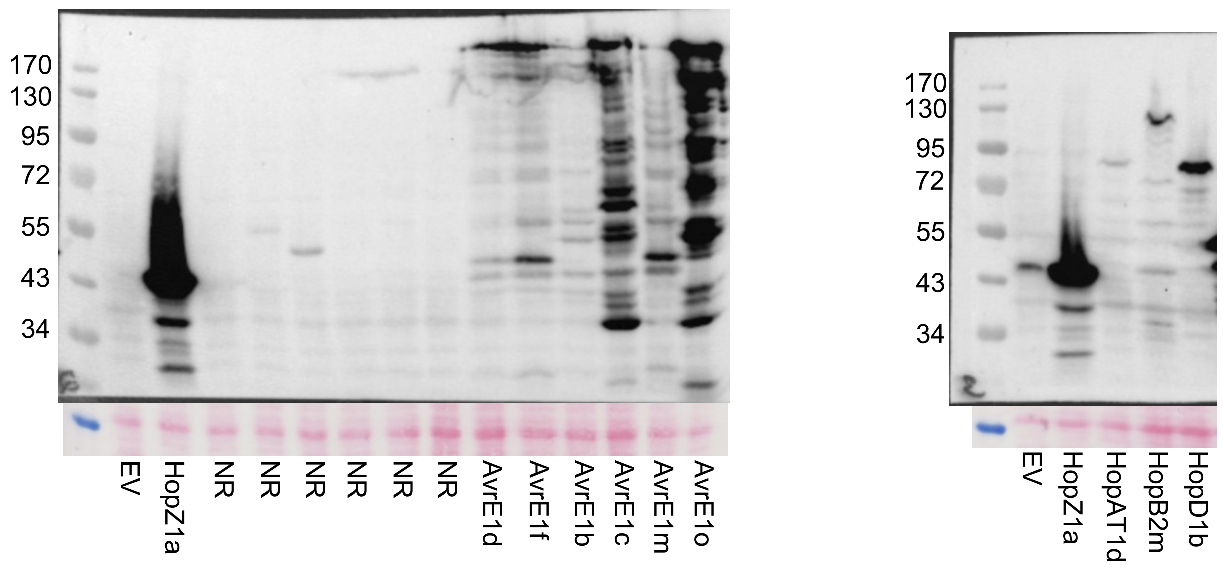
