## Supplementary material for "Metaeffector interactions modulate the type III effector-triggered immunity load of *Pseudomonas syringae*": Fig S2

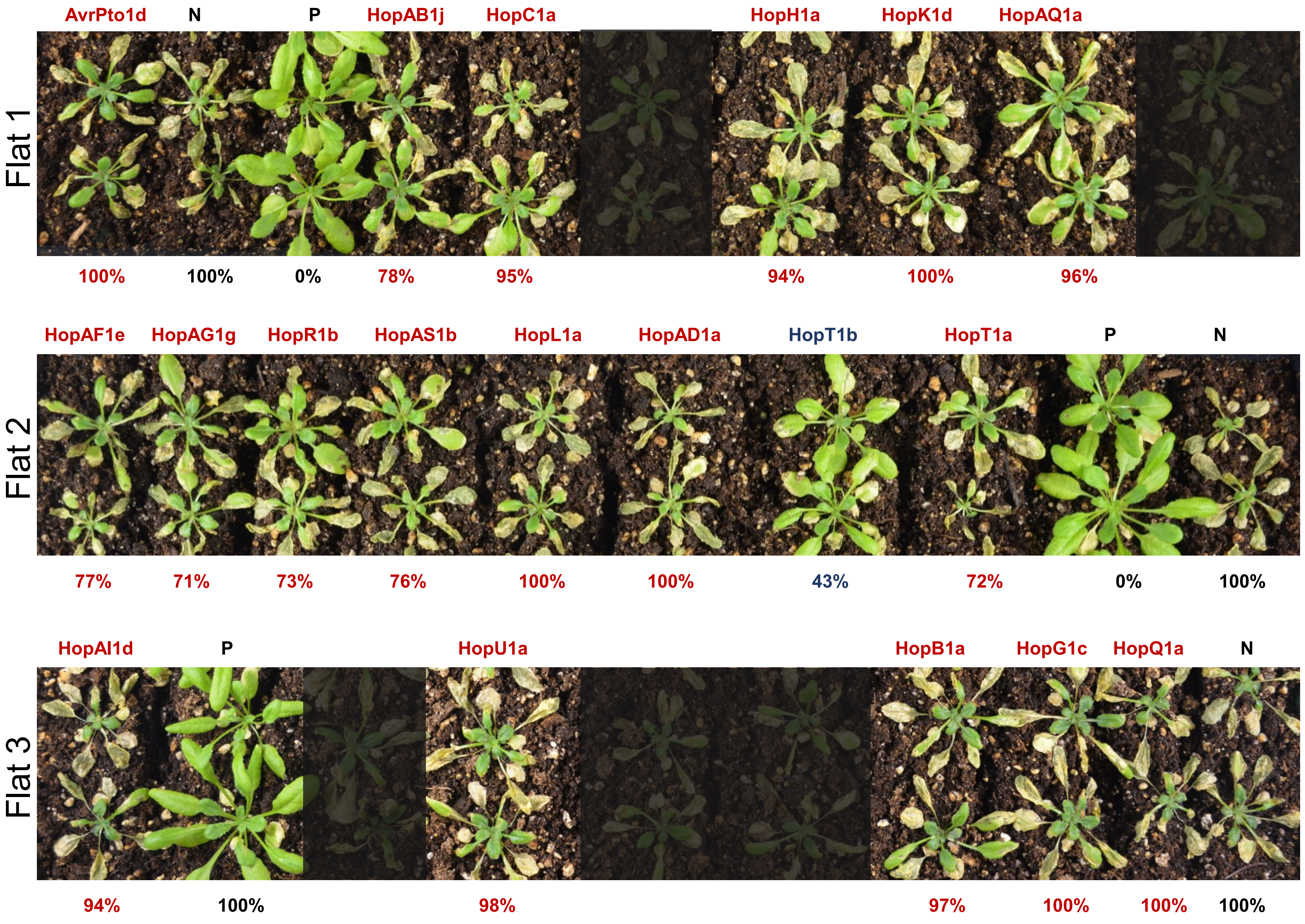

Flat 1
AvrPto1d
N
P
HopAB1j
HopC1a
HopH1a
HopK1d
HopAQ1a
100%
100%
0%
78%
95%
94%
100%
96%
Flat 2
HopAF1e
HopAG1g
HopR1b
HopAS1b
HopL1a
HopAD1a
HopT1b
HopT1a
P
N
77%
71%
73%
76%
100%
100%
43%
72%
0%
100%
Flat 3
HopAI1d
P
HopU1a
HopB1a
HopG1c
HopQ1a
N
94%
100%
98%
97%
100%
100%
100%

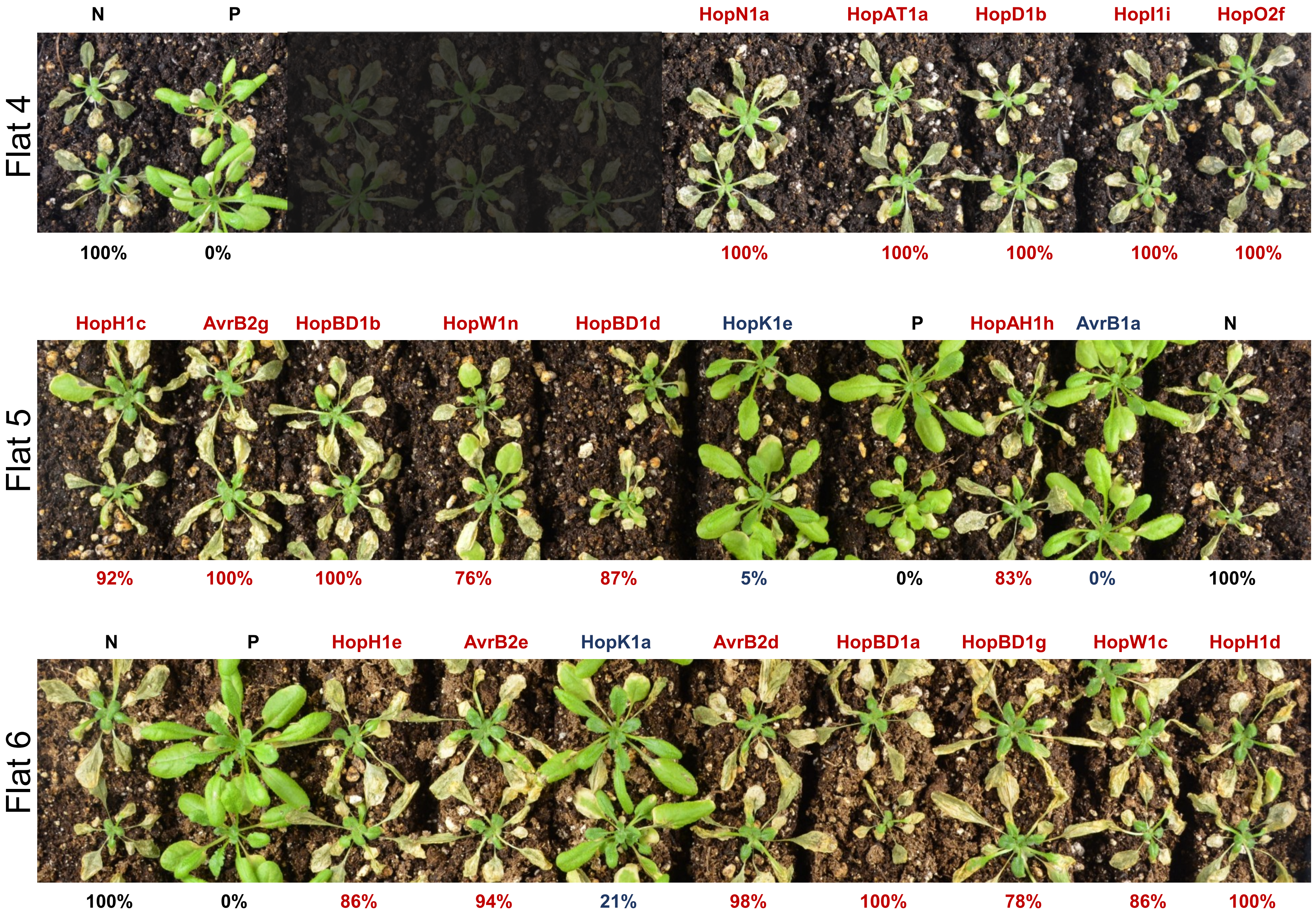

Flat 4
N
P
HopN1a
HopAT1a
HopD1b
HopI1i
HopO2f
100%
0%
100%
100%
100%
100%
100%
Flat 5
HopH1c
AvrB2g
HopBD1b
HopW1n
HopBD1d
HopK1e
P
HopAH1h
AvrB1a
N
92%
100%
100%
76%
87%
5%
0%
83%
0%
100%
Flat 6
N
P
HopH1e
AvrB2e
HopK1a
AvrB2d
HopBD1a
HopBD1g
HopW1c
HopH1d
100%
0%
86%
94%
21%
98%
100%
78%
86%
100%

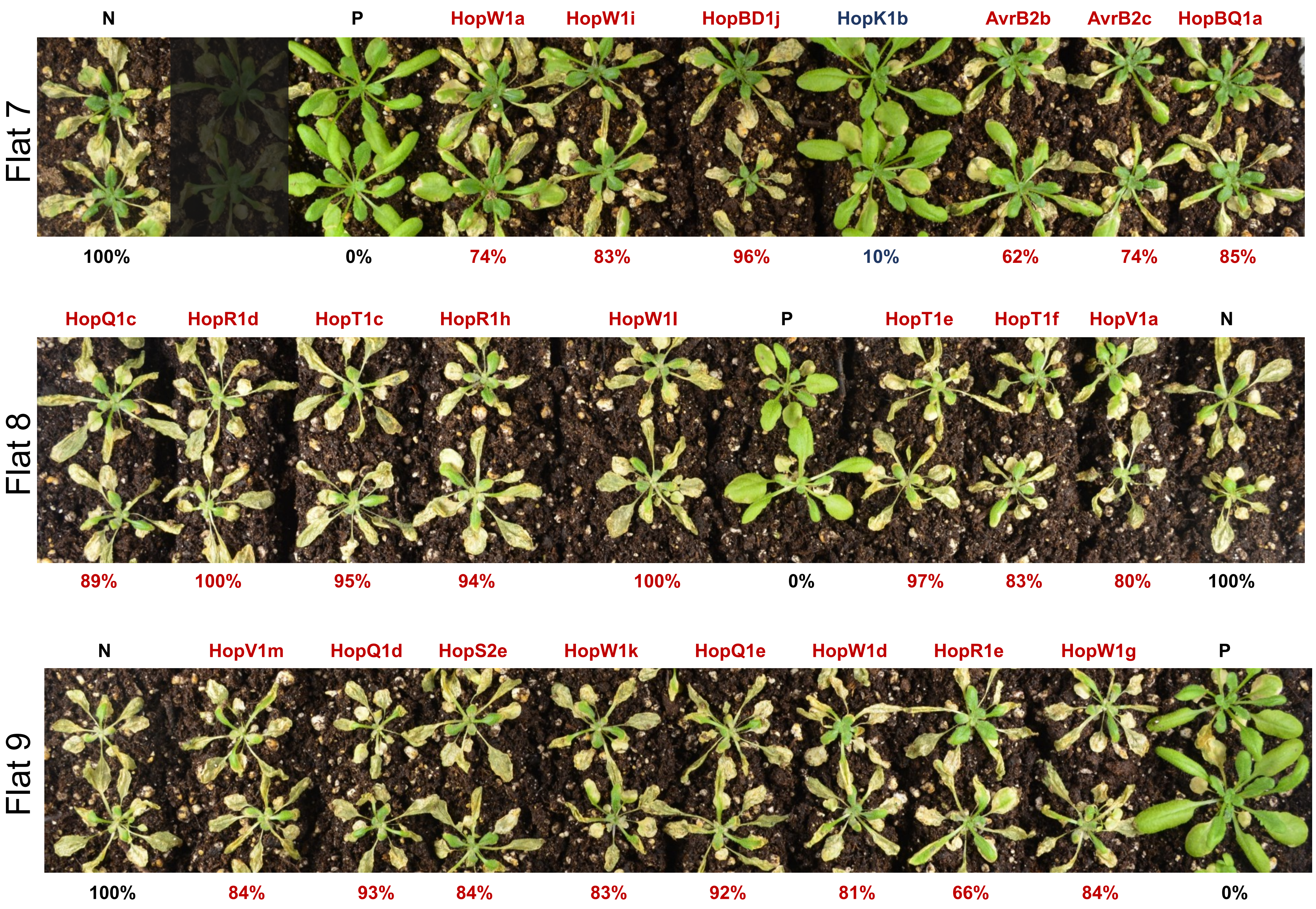
Flat 7
N
P
HopW1a
HopW1i
HopBD1j
HopK1b
AvrB2b
AvrB2c
HopBQ1a
100%
0%
74%
83%
96%
10%
62%
74%
85%
Flat 8
HopQ1c
HopR1d
HopT1c
HopR1h
HopW1l
P
HopT1e
HopT1f
HopV1a
N
89%
100%
95%
94%
100%
0%
97%
83%
80%
100%
Flat 9
N
HopV1m
HopQ1d
HopS2e
HopW1k
HopQ1e
HopW1d
HopR1e
HopW1g
P
100%
84%
93%
84%
83%
92%
81%
66%
84%
0%

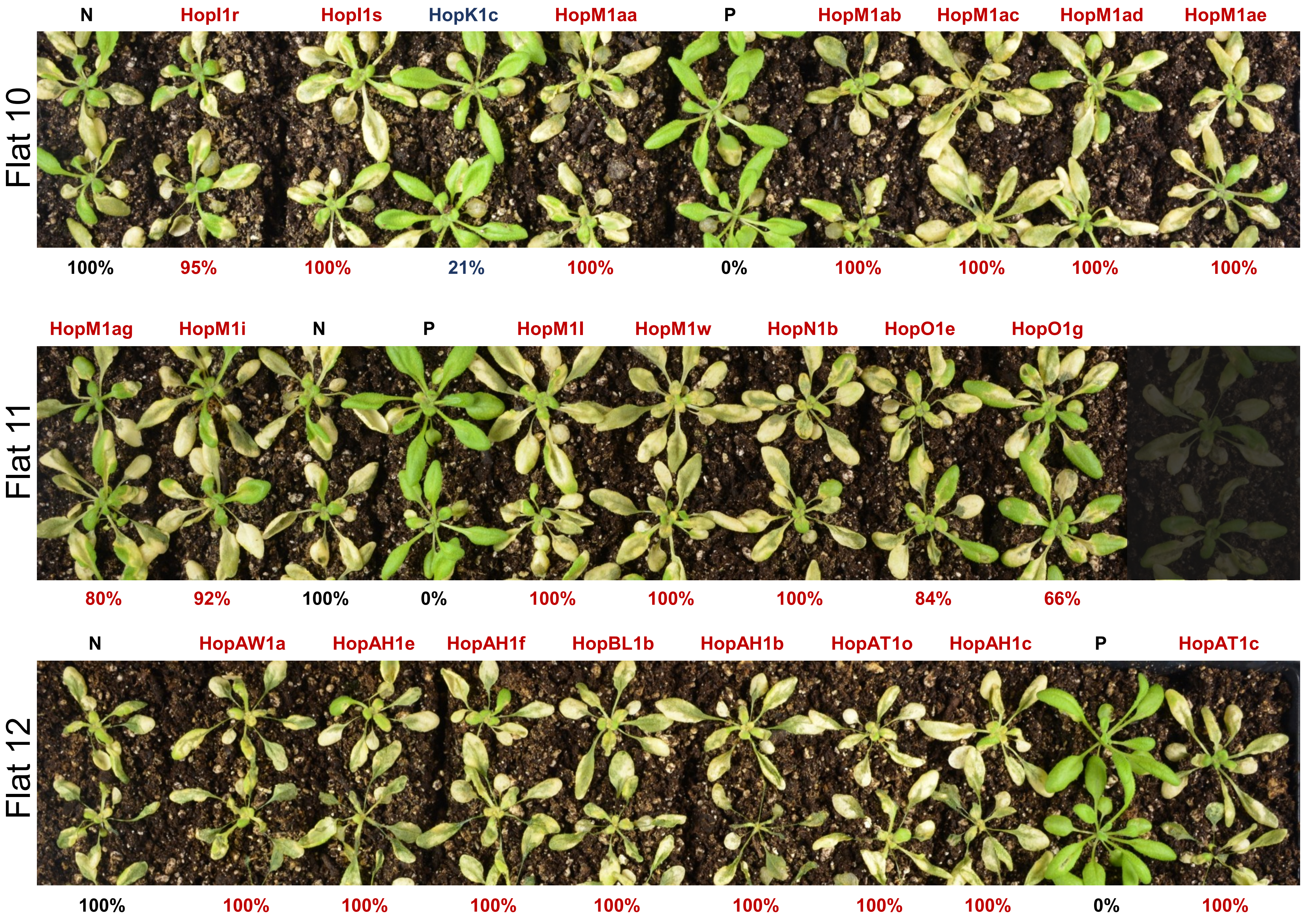
Flat 10
N
HopI1r
HopI1s
HopK1c
HopM1aa
P
HopM1ab
HopM1ac
HopM1ad
HopM1ae
100%
95%
100%
21%
100%
0%
100%
100%
100%
100%
Flat 11
HopM1ag
HopM1i
N
P
HopM1l
HopM1w
HopN1b
HopO1e
HopO1g
80%
92%
100%
0%
100%
100%
100%
84%
66%
Flat 12
N
HopAW1a
HopAH1e
HopAH1f
HopBL1b
HopAH1b
HopAT1o
HopAH1c
P
HopAT1c
100%
100%
100%
100%
100%
100%
100%
100%
0%
100%

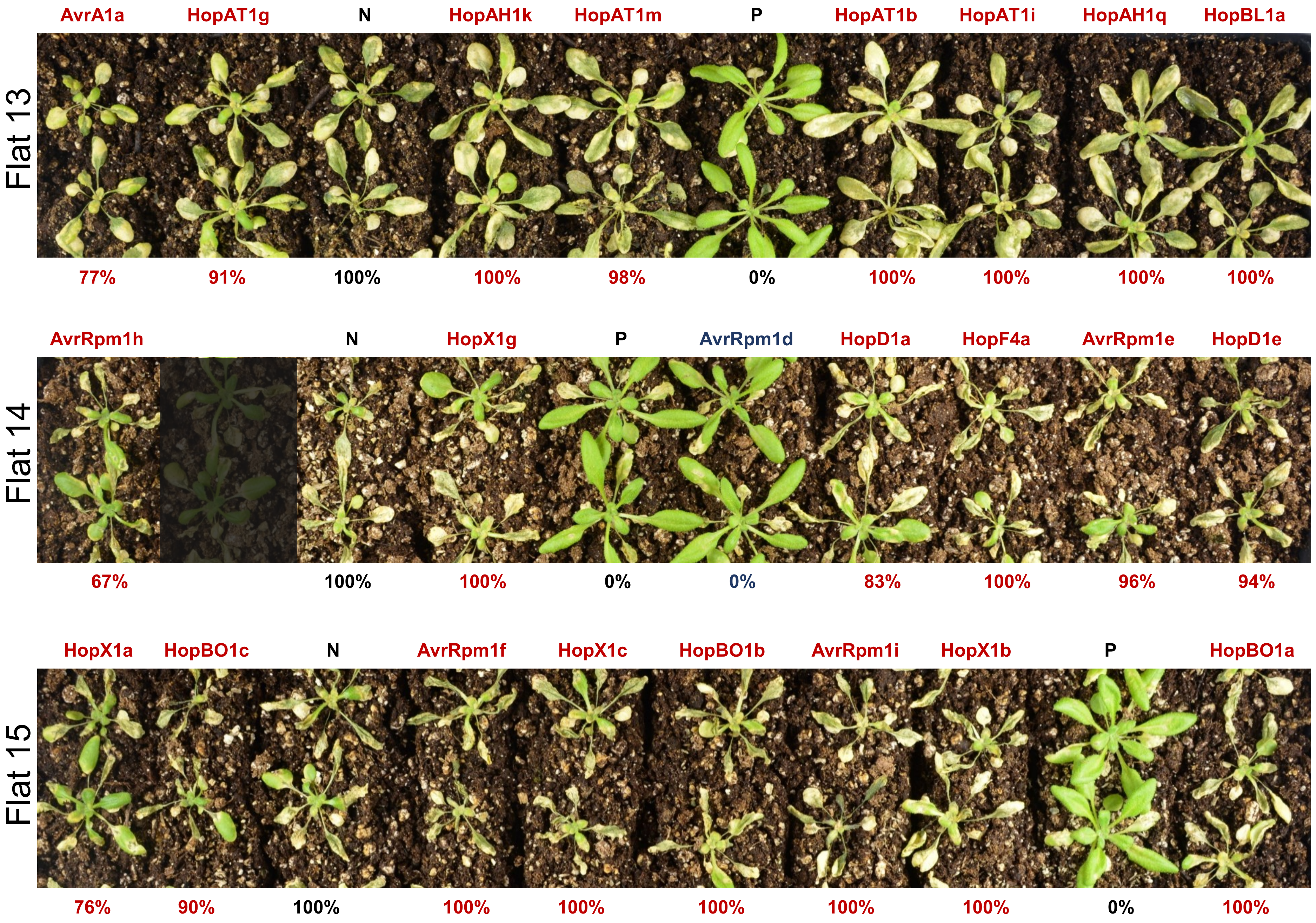
Flat 13
AvrA1a
HopAT1g
N
HopAH1k
HopAT1m
P
HopAT1b
HopAT1i
HopAH1q
HopBL1a
77%
91%
100%
100%
98%
0%
100%
100%
100%
100%
Flat 14
AvrRpm1h
N
HopX1g
P
AvrRpm1d
HopD1a
HopF4a
AvrRpm1e
HopD1e
67%
100%
100%
0%
0%
83%
100%
96%
94%
Flat 15
HopX1a
HopBO1c
N
AvrRpm1f
HopX1c
HopBO1b
AvrRpm1i
HopX1b
P
HopBO1a
76%
90%
100%
100%
100%
100%
100%
100%
0%
100%

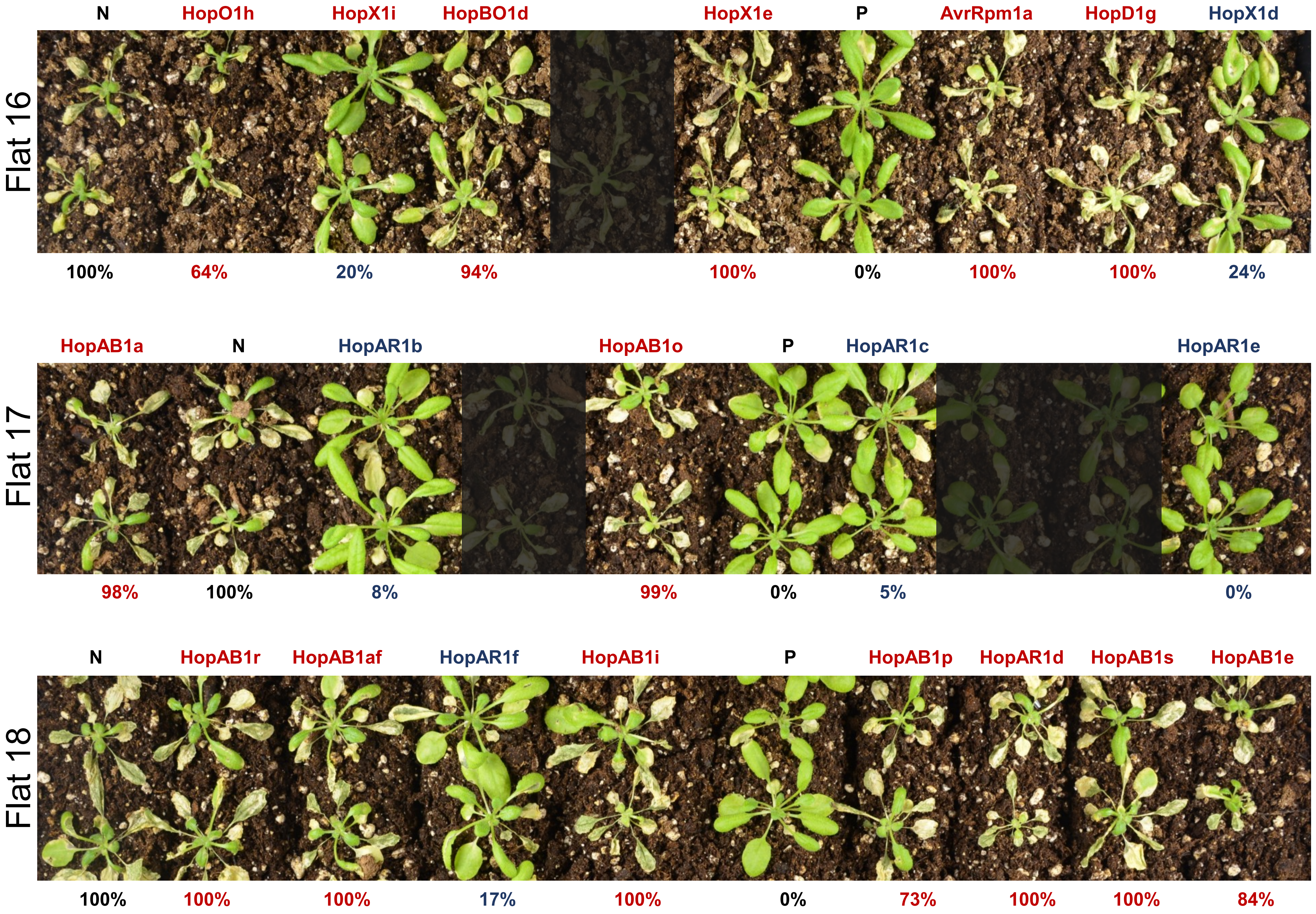

Flat 16
N
HopO1h
HopX1i
HopBO1d
HopX1e
P
AvrRpm1a
HopD1g
HopX1d
100%
64%
20%
94%
100%
0%
100%
100%
24%
Flat 17
HopAB1a
N
HopAR1b
HopAB1o
P
HopAR1c
HopAR1e
98%
100%
8%
99%
0%
5%
0%
Flat 18
N
HopAB1r
HopAB1af
HopAR1f
HopAB1i
P
HopAB1p
HopAR1d
HopAB1s
HopAB1e
100%
100%
100%
17%
100%
0%
73%
100%
100%
84%

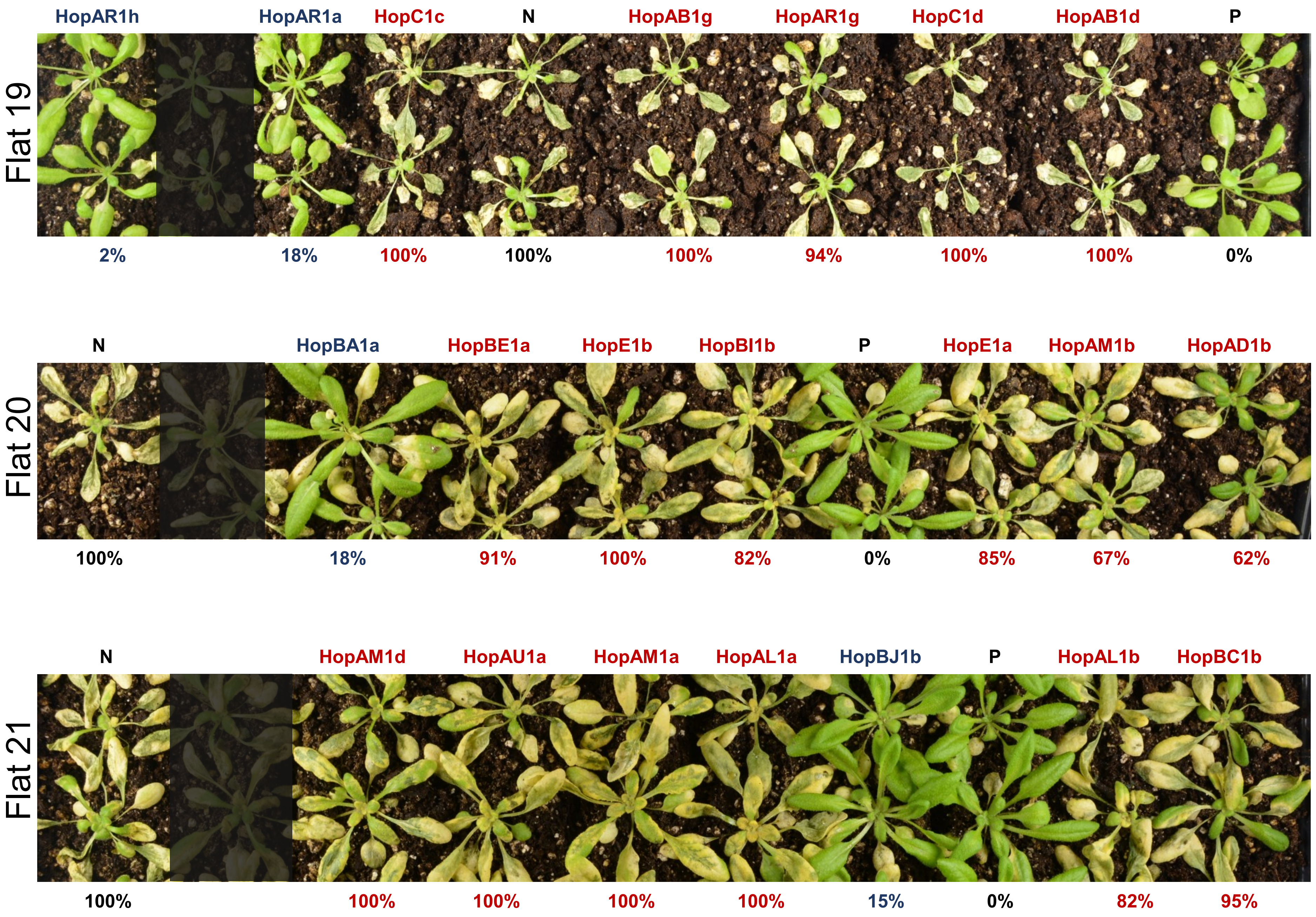
Flat 19
HopAR1h
HopAR1a
HopC1c
N
HopAB1g
HopAR1g
HopC1d
HopAB1d
P
2%
18%
100%
100%
100%
94%
100%
100%
0%
Flat 20
N
HopBA1a
HopBE1a
HopE1b
HopBI1b
P
HopE1a
HopAM1b
HopAD1b
100%
18%
91%
100%
82%
0%
85%
67%
62%
Flat 21
N
HopAM1d
HopAU1a
HopAM1a
HopAL1a
HopBJ1b
P
HopAL1b
HopBC1b
100%
100%
100%
100%
100%
15%
0%
82%
95%

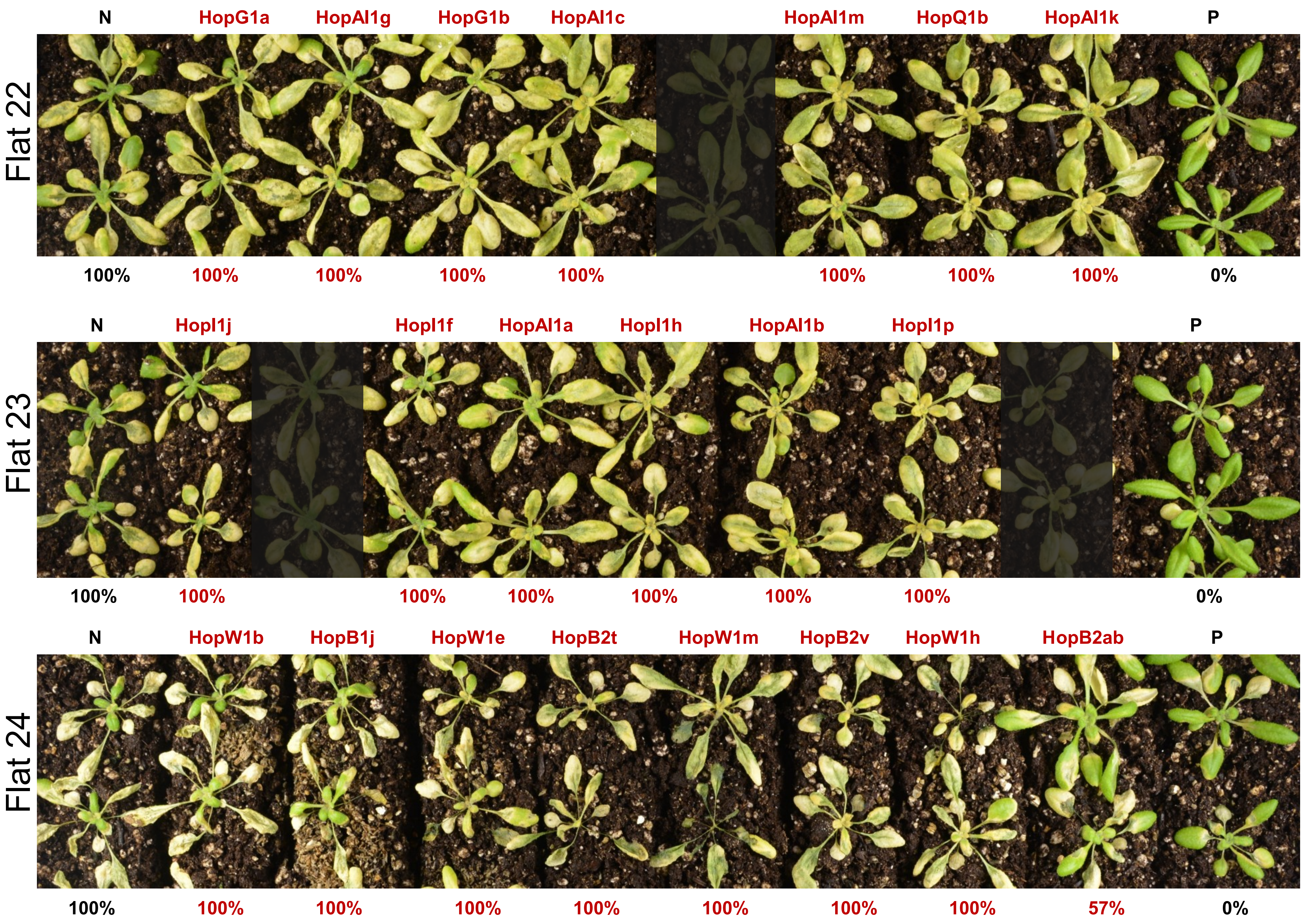

Flat 22
N
HopG1a
HopAl1g
HopG1b
HopAl1c
HopAl1m
HopQ1b
HopAl1k
P
100%
100%
100%
100%
100%
100%
100%
100%
0%
Flat 23
N
HopI1j
HopI1f
HopAl1a
HopI1h
HopAl1b
HopI1p
P
100%
100%
100%
100%
100%
100%
100%
0%
Flat 24
N
HopW1b
HopB1j
HopW1e
HopB2t
HopW1m
HopB2v
HopW1h
HopB2ab
P
100%
100%
100%
100%
100%
100%
100%
100%
57%
0%

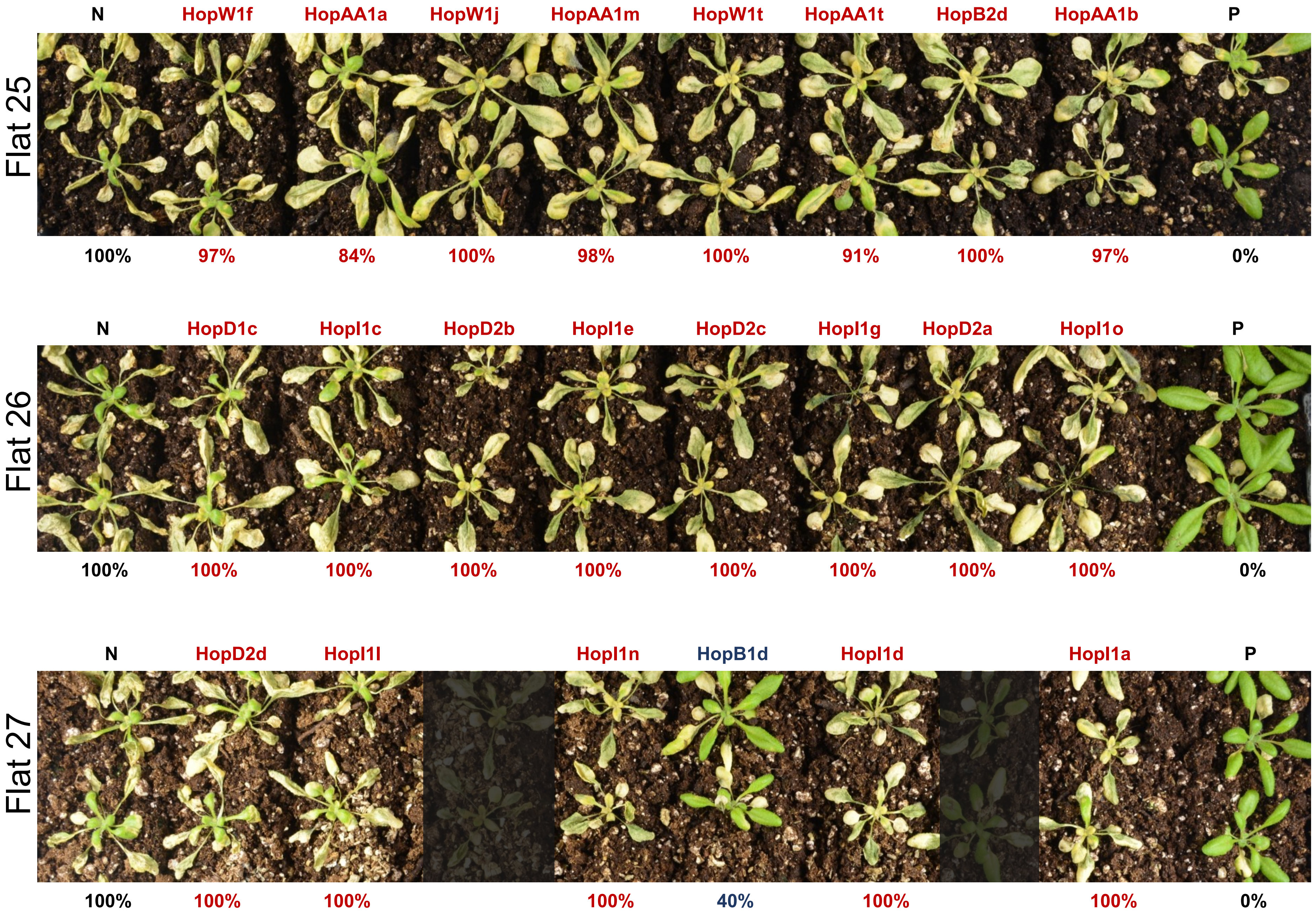

Flat 25
N
HopW1f
HopAA1a
HopW1j
HopAA1m
HopW1t
HopAA1t
HopB2d
HopAA1b
P
100%
97%
84%
100%
98%
100%
91%
100%
97%
0%
Flat 26
N
HopD1c
HopI1c
HopD2b
HopI1e
HopD2c
HopI1g
HopD2a
HopI1o
P
100%
100%
100%
100%
100%
100%
100%
100%
100%
0%
Flat 27
N
HopD2d
HopI1l
HopI1n
HopB1d
HopI1d
HopI1a
P
100%
100%
100%
100%
40%
100%
100%
0%

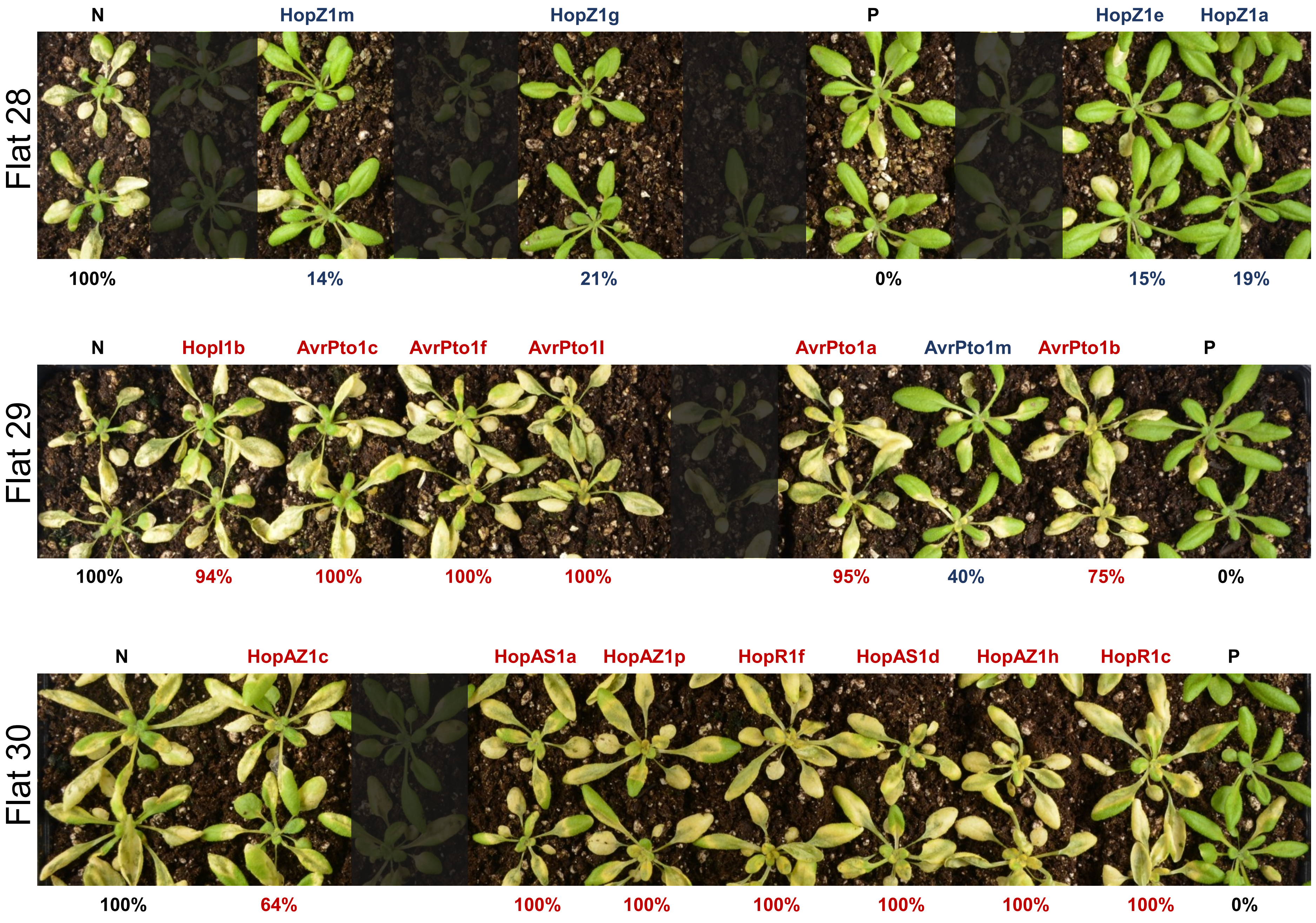

Flat 28
N
HopZ1m
HopZ1g
P
HopZ1e
HopZ1a
100%
14%
21%
0%
15%
19%
Flat 29
N
HopI1b
AvrPto1c
AvrPto1f
AvrPto1l
AvrPto1a
AvrPto1m
AvrPto1b
P
100%
94%
100%
100%
100%
95%
40%
75%
0%
Flat 30
N
HopAZ1c
HopAS1a
HopAZ1p
HopR1f
HopAS1d
HopAZ1h
HopR1c
P
100%
64%
100%
100%
100%
100%
100%
100%
0%

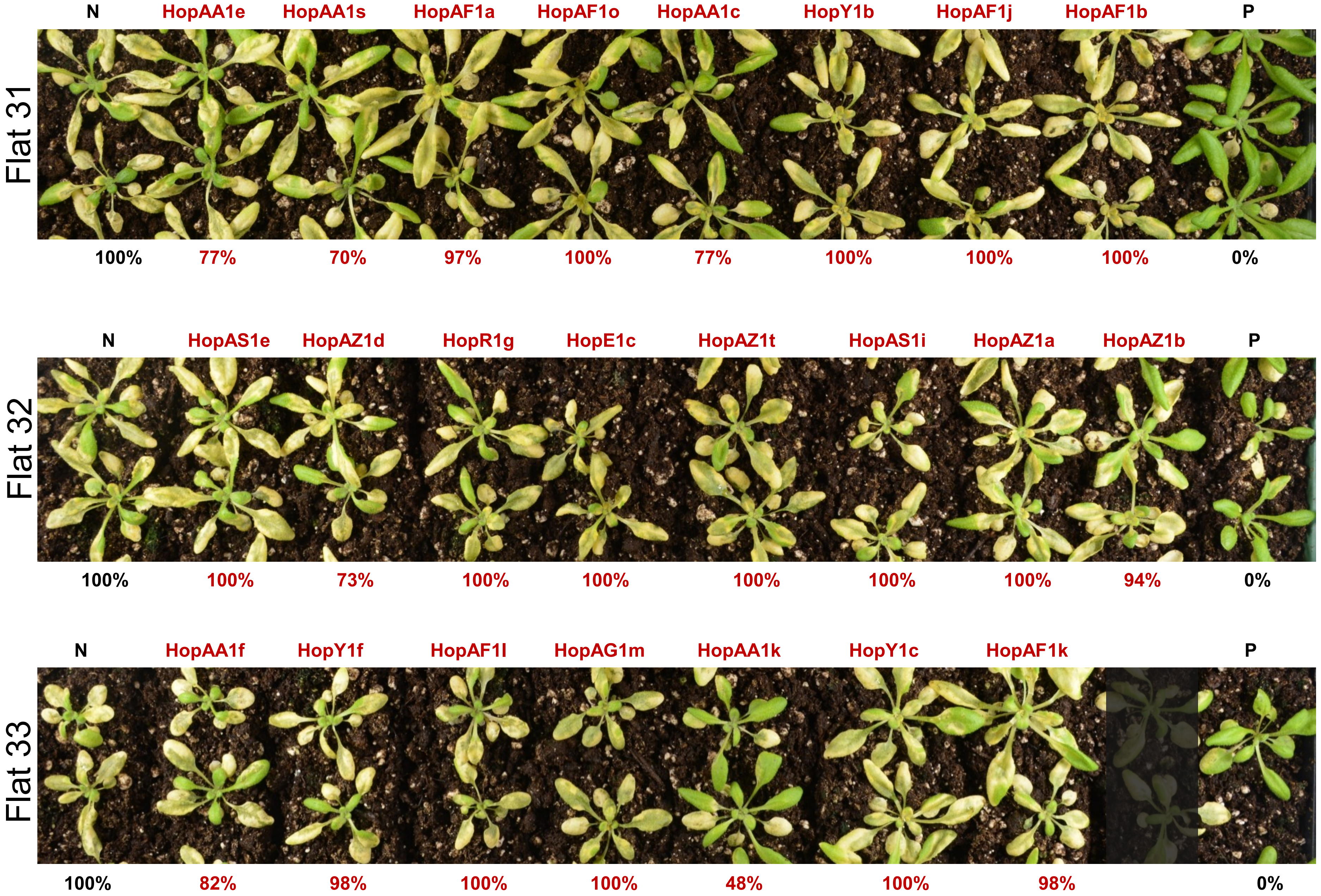
Flat 31
N
HopAA1e
HopAA1s
HopAF1a
HopAF1o
HopAA1c
HopY1b
HopAF1j
HopAF1b
P
100%
77%
70%
97%
100%
77%
100%
100%
100%
0%
Flat 32
N
HopAS1e
HopAZ1d
HopR1g
HopE1c
HopAZ1t
HopAS1i
HopAZ1a
HopAZ1b
P
100%
100%
73%
100%
100%
100%
100%
100%
94%
0%
Flat 33
N
HopAA1f
HopY1f
HopAF1l
HopAG1m
HopAA1k
HopY1c
HopAF1k
P
100%
82%
98%
100%
100%
48%
100%
98%
0%

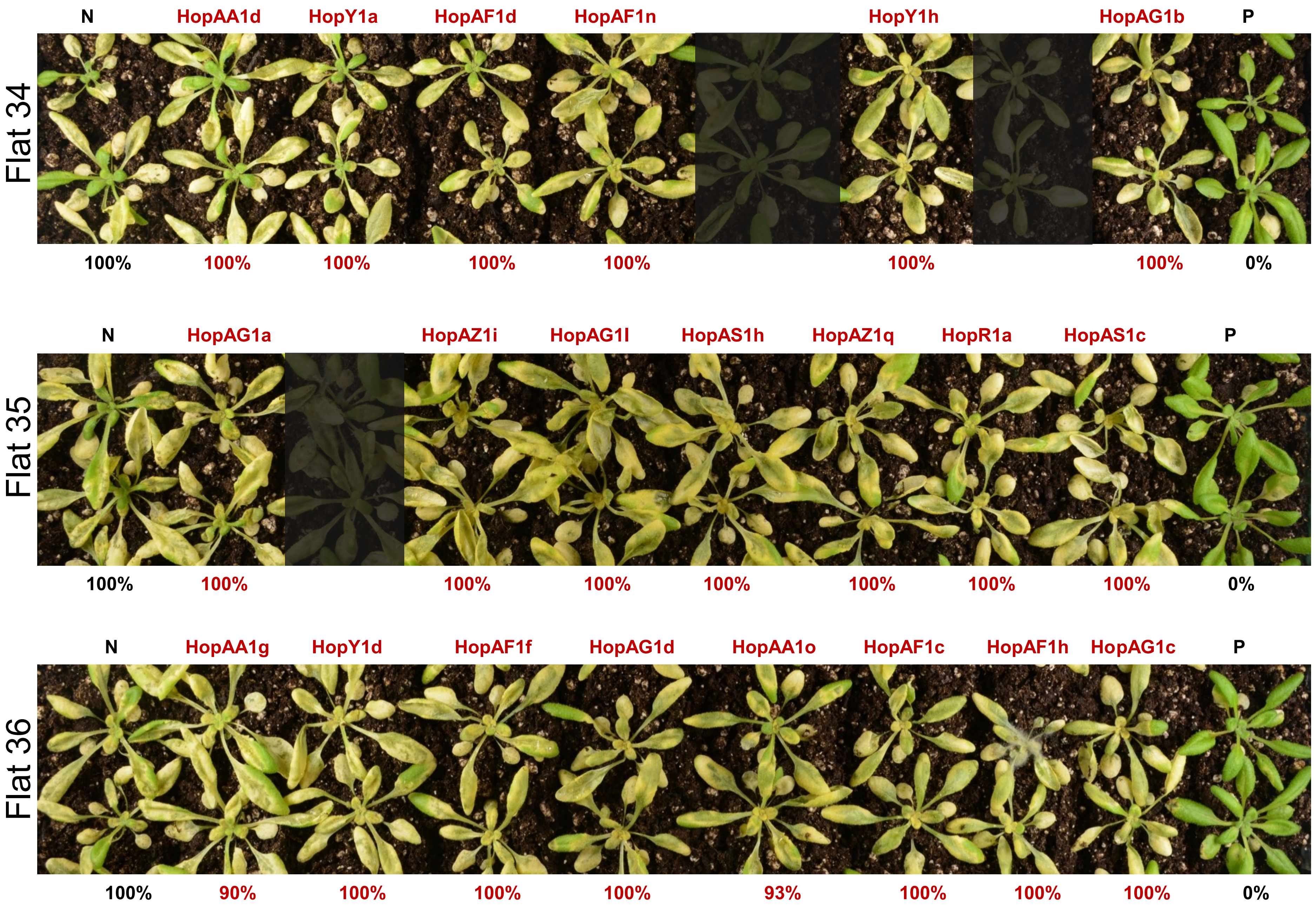
Flat 34
N
HopAA1d
HopY1a
HopAF1d
HopAF1n
HopY1h
HopAG1b
P
100%
100%
100%
100%
100%
100%
100%
0%
Flat 35
N
HopAG1a
HopAZ1i
HopAG1l
HopAS1h
HopAZ1q
HopR1a
HopAS1c
P
100%
100%
100%
100%
100%
100%
100%
100%
0%
Flat 36
N
HopAA1g
HopY1d
HopAF1f
HopAG1d
HopAA1o
HopAF1c
HopAF1h
HopAG1c
P
100%
90%
100%
100%
100%
93%
100%
100%
100%
0%

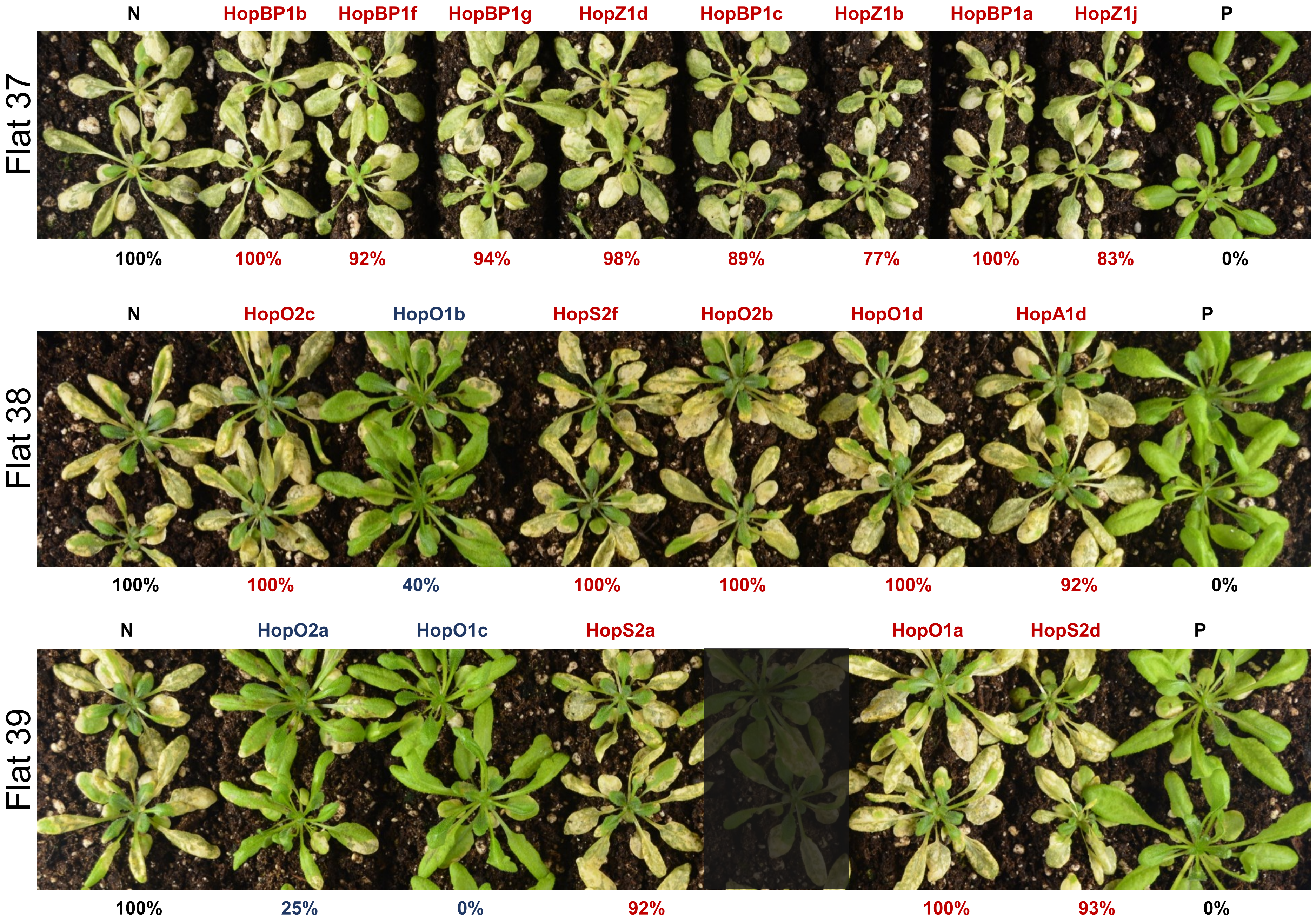

Flat 37
N
HopBP1b
HopBP1f
HopBP1g
HopZ1d
HopBP1c
HopZ1b
HopBP1a
HopZ1j
P
100%
100%
92%
94%
98%
89%
77%
100%
83%
0%
Flat 38
N
HopO2c
HopO1b
HopS2f
HopO2b
HopO1d
HopA1d
P
100%
100%
40%
100%
100%
100%
92%
0%
Flat 39
N
HopO2a
HopO1c
HopS2a
HopO1a
HopS2d
P
100%
25%
0%
92%
100%
93%
0%

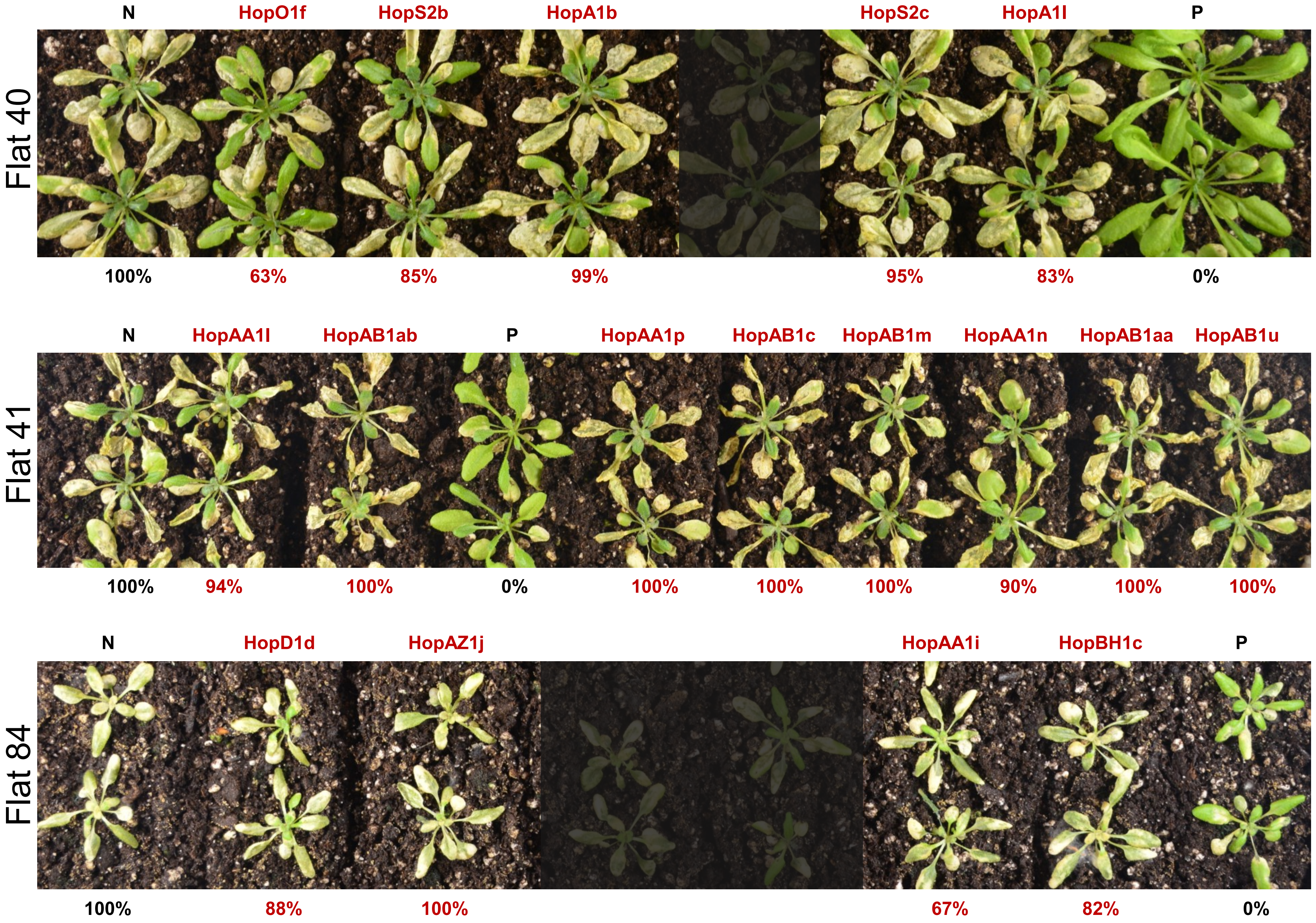
Flat 40
N
HopO1f
HopS2b
HopA1b
HopS2c
HopA1l
P
100%
63%
85%
99%
95%
83%
0%
Flat 41
N
HopAA1l
HopAB1ab
P
HopAA1p
HopAB1c
HopAB1m
HopAA1n
HopAB1aa
HopAB1u
100%
94%
100%
0%
100%
100%
100%
90%
100%
100%
Flat 84
N
HopD1d
HopAZ1j
HopAA1i
HopBH1c
P
100%
88%
100%
67%
82%
0%

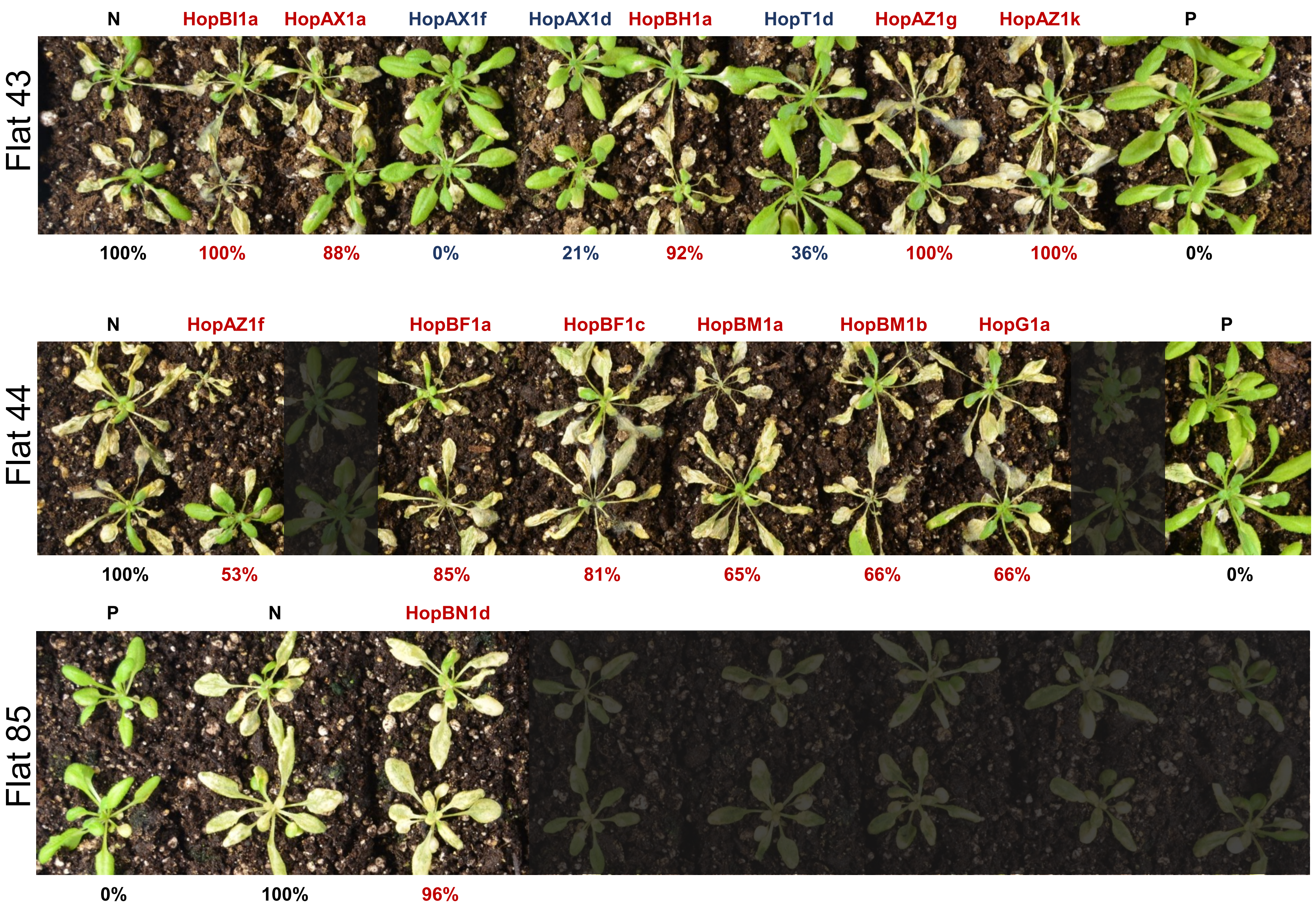
Flat 43
N
HopBI1a
HopAX1a
HopAX1f
HopAX1d
HopBH1a
HopT1d
HopAZ1g
HopAZ1k
P
100%
100%
88%
0%
21%
92%
36%
100%
100%
0%
Flat 44
N
HopAZ1f
HopBF1a
HopBF1c
HopBM1a
HopBM1b
HopG1a
P
100%
53%
85%
81%
65%
66%
66%
0%
Flat 85
P
N
HopBN1d
0%
100%
96%

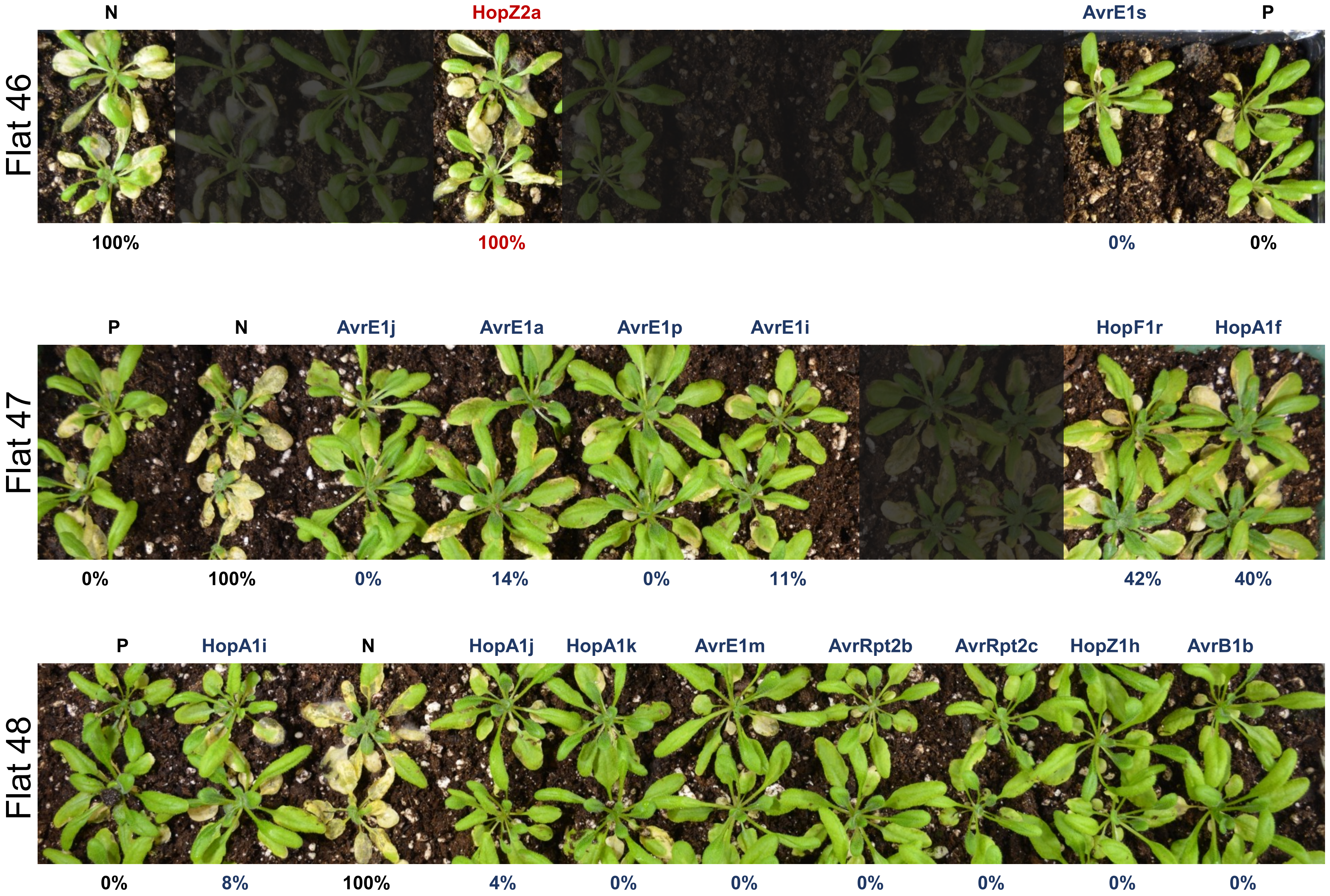
Flat 46
N
HopZ2a
AvrE1s
P
100%
100%
0%
0%
Flat 47
P
N
AvrE1j
AvrE1a
AvrE1p
AvrE1i
HopF1r
HopA1f
0%
100%
0%
14%
0%
11%
42%
40%
Flat 48
P
HopA1i
N
HopA1j
HopA1k
AvrE1m
AvrRpt2b
AvrRpt2c
HopZ1h
AvrB1b
0%
8%
100%
4%
0%
0%
0%
0%
0%
0%

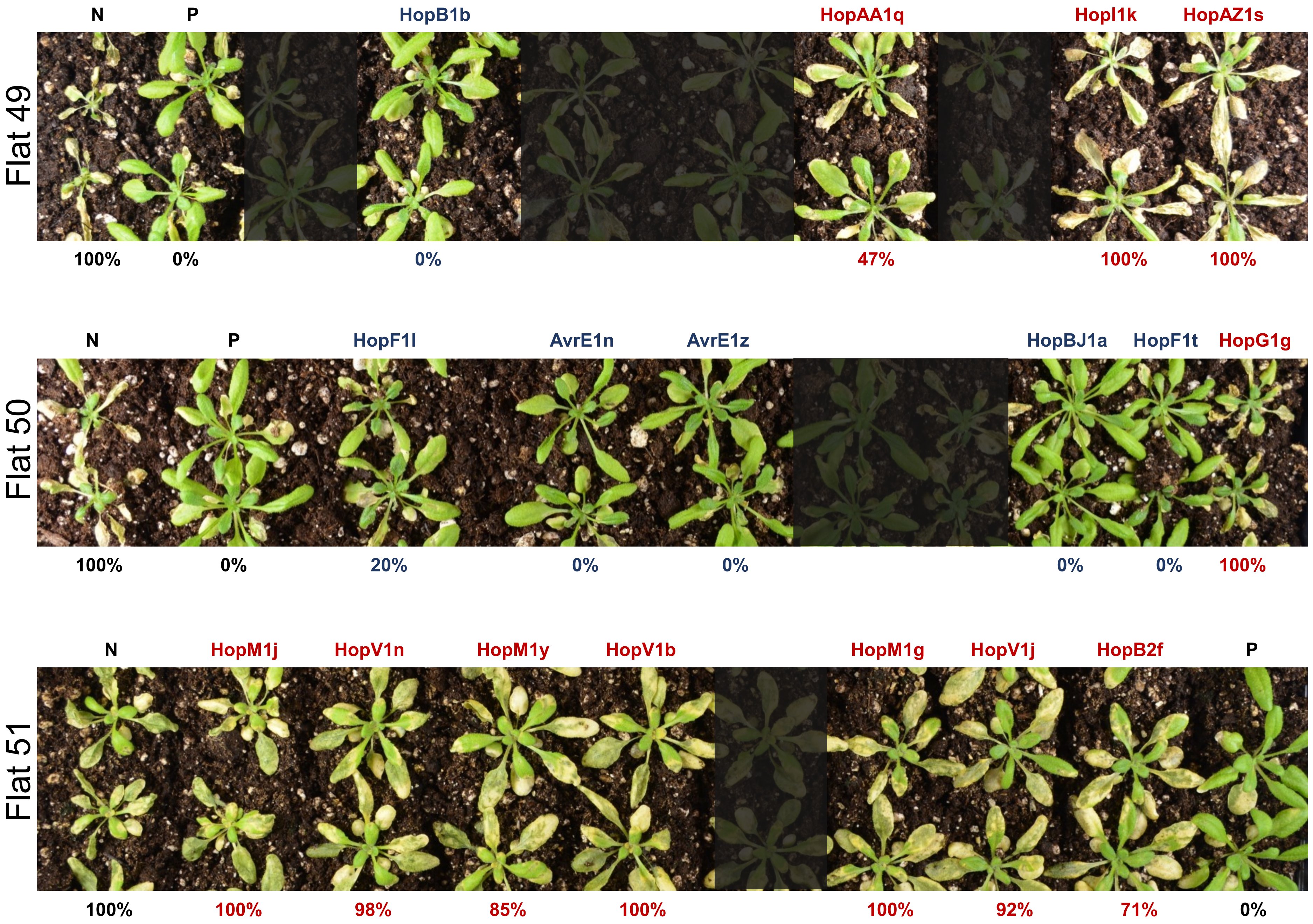

Flat 49
N
P
HopB1b
HopAA1q
HopI1k
HopAZ1s
100%
0%
0%
47%
100%
100%
Flat 50
N
P
HopF1l
AvrE1n
AvrE1z
HopBJ1a
HopF1t
HopG1g
100%
0%
20%
0%
0%
0%
0%
100%
Flat 51
N
HopM1j
HopV1n
HopM1y
HopV1b
HopM1g
HopV1j
HopB2f
P
100%
100%
98%
85%
100%
100%
92%
71%
0%

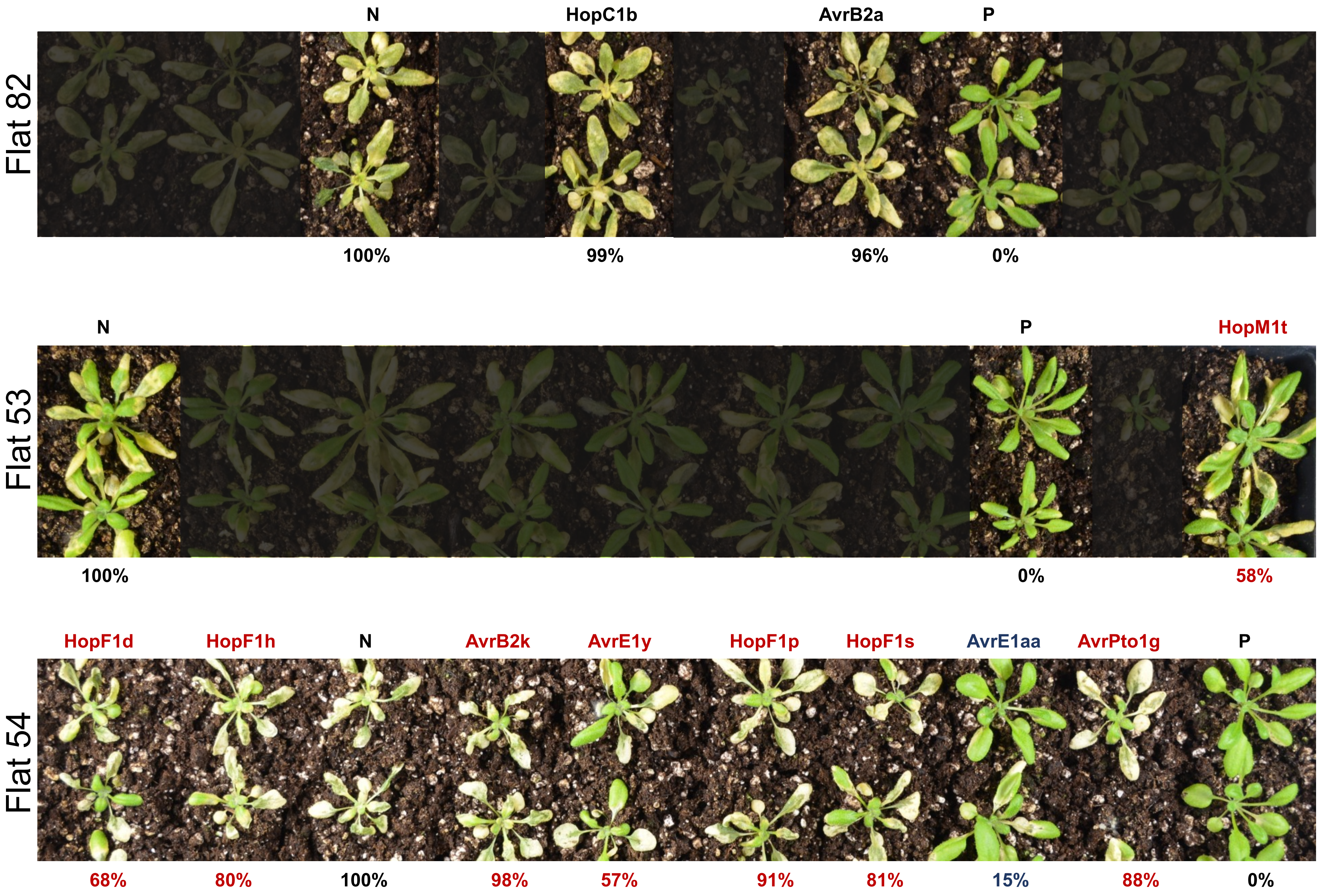
Flat 82
N
HopC1b
AvrB2a
P
100%
99%
96%
0%
Flat 53
N
P
HopM1t
100%
0%
58%
Flat 54
HopF1d
HopF1h
N
AvrB2k
AvrE1y
HopF1p
HopF1s
AvrE1aa
AvrPto1g
P
68%
80%
100%
98%
57%
91%
81%
15%
88%
0%

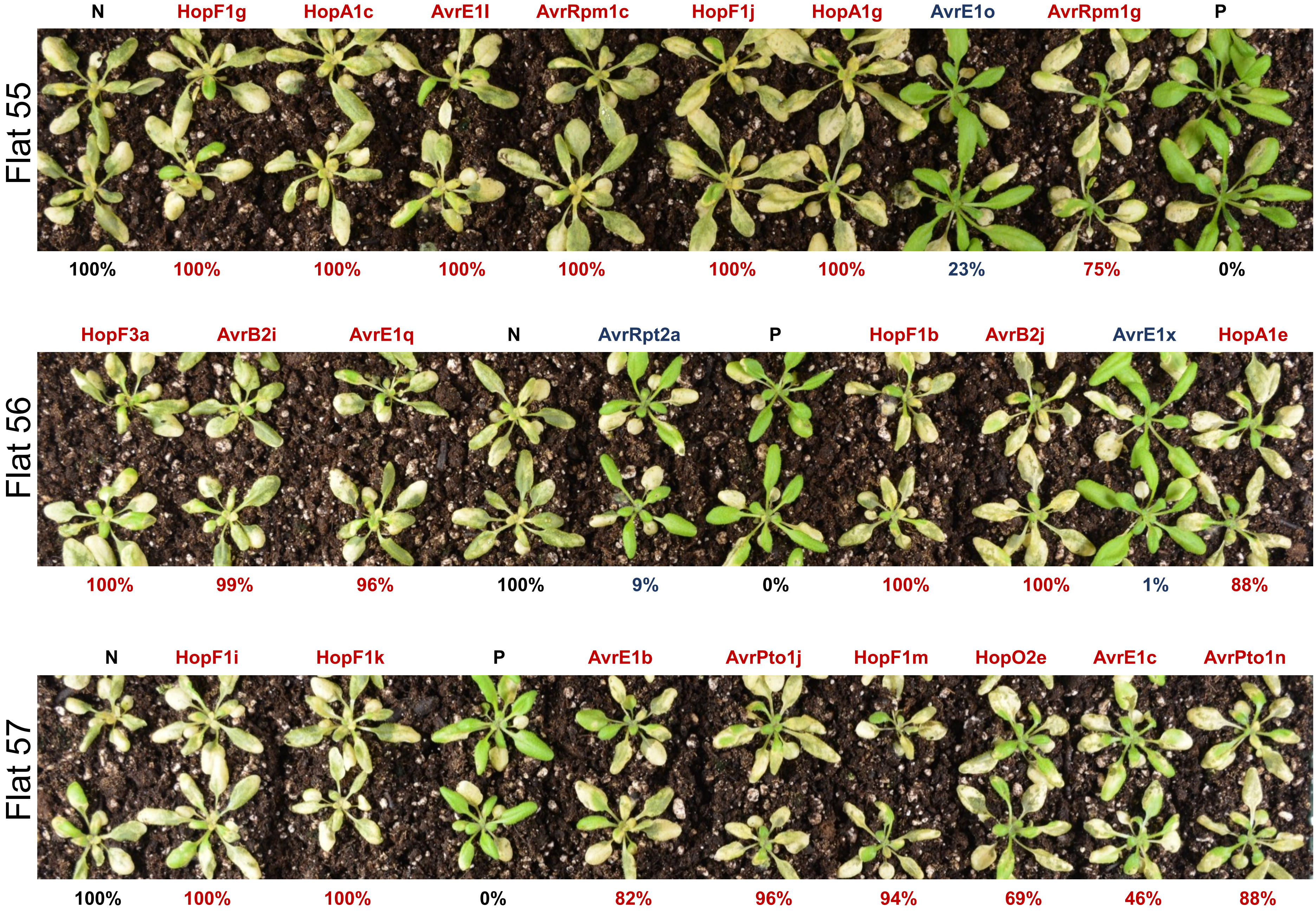
Flat 55
N
HopF1g
HopA1c
AvrE1l
AvrRpm1c
HopF1j
HopA1g
AvrE1o
AvrRpm1g
P
100%
100%
100%
100%
100%
100%
100%
23%
75%
0%
Flat 56
HopF3a
AvrB2i
AvrE1q
N
AvrRpt2a
P
HopF1b
AvrB2j
AvrE1x
HopA1e
100%
99%
96%
100%
9%
0%
100%
100%
1%
88%
Flat 57
N
HopF1i
HopF1k
P
AvrE1b
AvrPto1j
HopF1m
HopO2e
AvrE1c
AvrPto1n
100%
100%
100%
0%
82%
96%
94%
69%
46%
88%

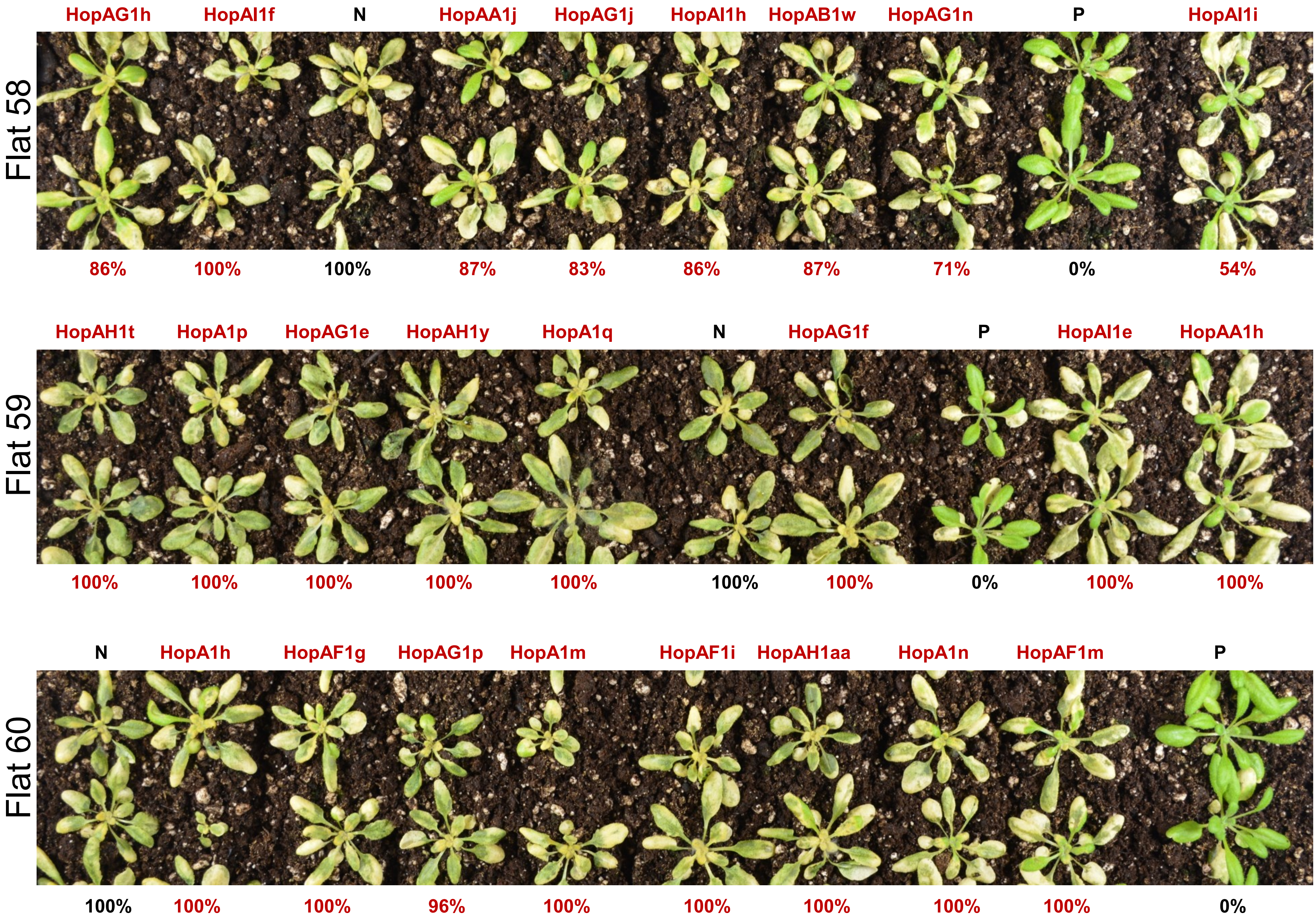
HopAG1h
HopAI1f
N
HopAA1j
HopAG1j
HopAI1h
HopAB1w
HopAG1n
P
HopAI1i
Flat 58
86%
100%
100%
87%
83%
86%
87%
71%
0%
54%
HopAH1t
HopA1p
HopAG1e
HopAH1y
HopA1q
N
HopAG1f
P
HopAI1e
HopAA1h
Flat 59
100%
100%
100%
100%
100%
100%
100%
0%
100%
100%
N
HopA1h
HopAF1g
HopAG1p
HopA1m
HopAF1i
HopAH1aa
HopA1n
HopAF1m
P
Flat 60
100%
100%
100%
96%
100%
100%
100%
100%
100%
0%

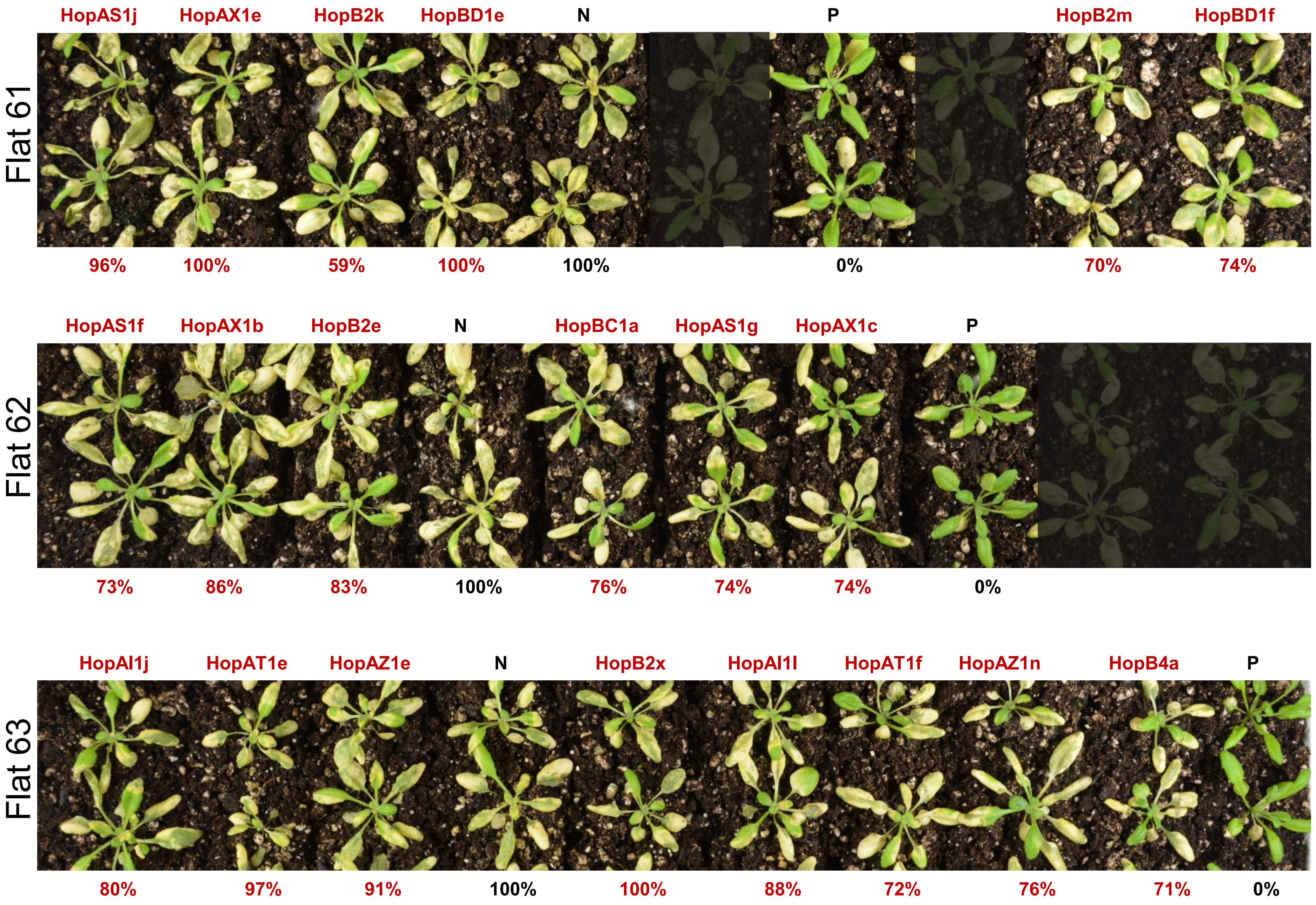
Flat 61
HopAS1j
HopAX1e
HopB2k
HopBD1e
N
P
HopB2m
HopBD1f
96%
100%
59%
100%
100%
0%
70%
74%
Flat 62
HopAS1f
HopAX1b
HopB2e
N
HopBC1a
HopAS1g
HopAX1c
P
73%
86%
83%
100%
76%
74%
74%
0%
Flat 63
HopAI1j
HopAT1e
HopAZ1e
N
HopB2x
HopAI1l
HopAT1f
HopAZ1n
HopB4a
P
80%
97%
91%
100%
100%
88%
72%
76%
71%
0%

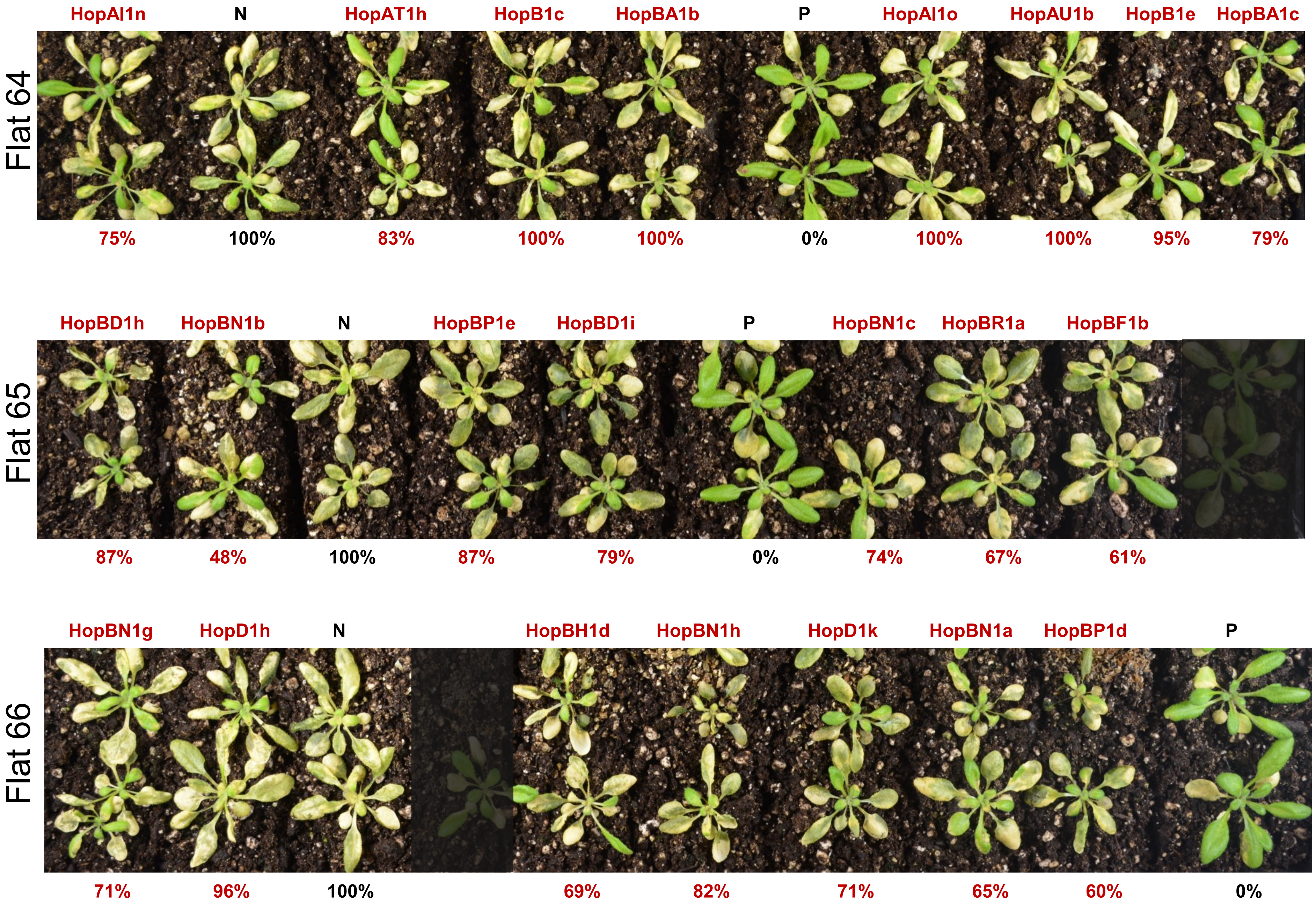
Flat 64
HopAI1n
N
HopAT1h
HopB1c
HopBA1b
P
HopAI1o
HopAU1b
HopB1e
HopBA1c
75%
100%
83%
100%
100%
0%
100%
100%
95%
79%
Flat 65
HopBD1h
HopBN1b
N
HopBP1e
HopBD1i
P
HopBN1c
HopBR1a
HopBF1b
87%
48%
100%
87%
79%
0%
74%
67%
61%
Flat 66
HopBN1g
HopD1h
N
HopBH1d
HopBN1h
HopD1k
HopBN1a
HopBP1d
P
71%
96%
100%
69%
82%
71%
65%
60%
0%

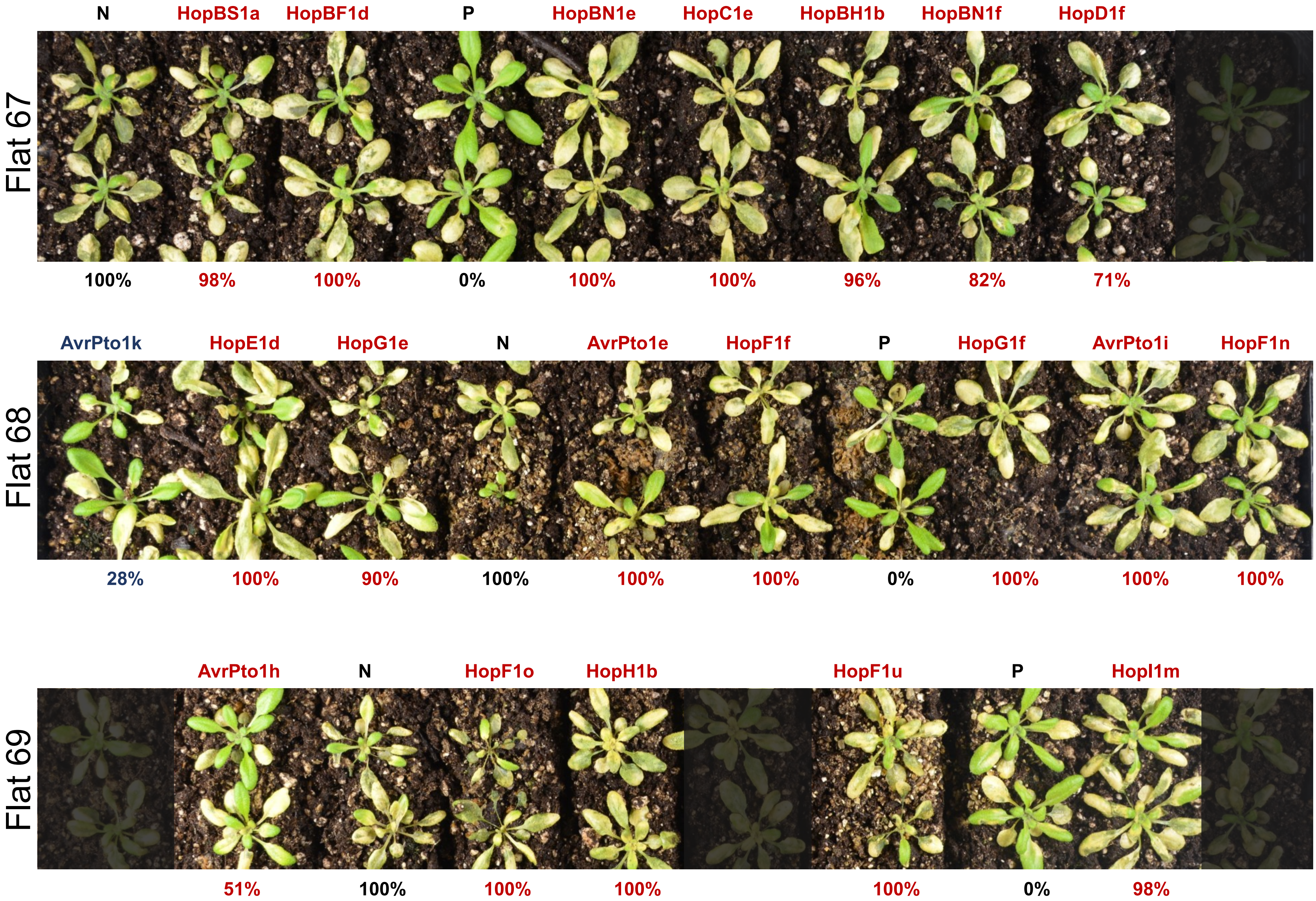
Flat 67
N
HopBS1a
HopBF1d
P
HopBN1e
HopC1e
HopBH1b
HopBN1f
HopD1f
100%
98%
100%
0%
100%
100%
96%
82%
71%
Flat 68
AvrPto1k
HopE1d
HopG1e
N
AvrPto1e
HopF1f
P
HopG1f
AvrPto1i
HopF1n
28%
100%
90%
100%
100%
100%
0%
100%
100%
100%
Flat 69
AvrPto1h
N
HopF1o
HopH1b
HopF1u
P
HopI1m
51%
100%
100%
100%
100%
0%
98%

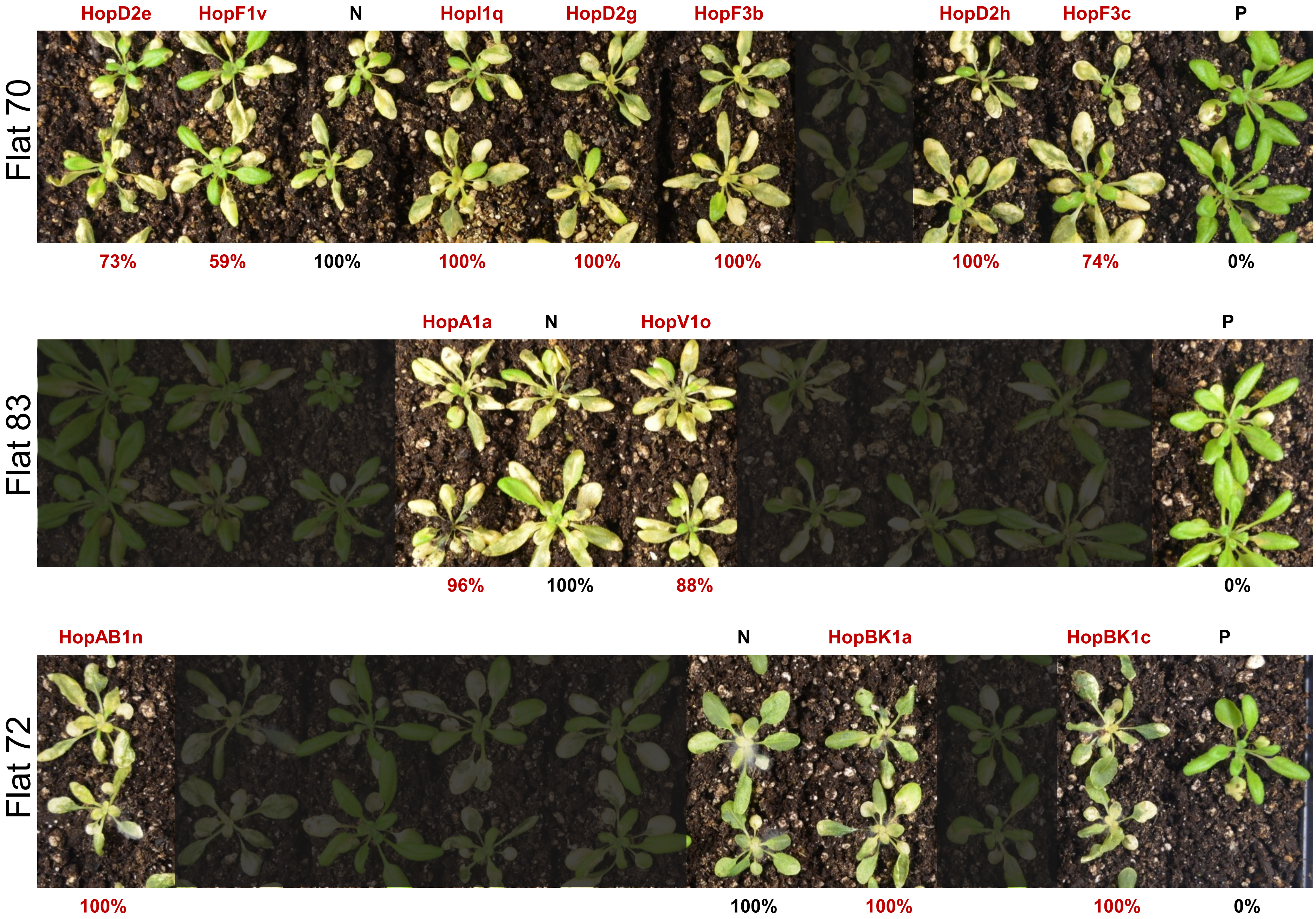
Flat 70
HopD2e
HopF1v
N
HopI1q
HopD2g
HopF3b
HopD2h
HopF3c
P
73%
59%
100%
100%
100%
100%
100%
74%
0%
Flat 83
HopA1a
N
HopV1o
P
96%
100%
88%
0%
Flat 72
HopAB1n
N
HopBK1a
HopBK1c
P
100%
100%
100%
100%
0%

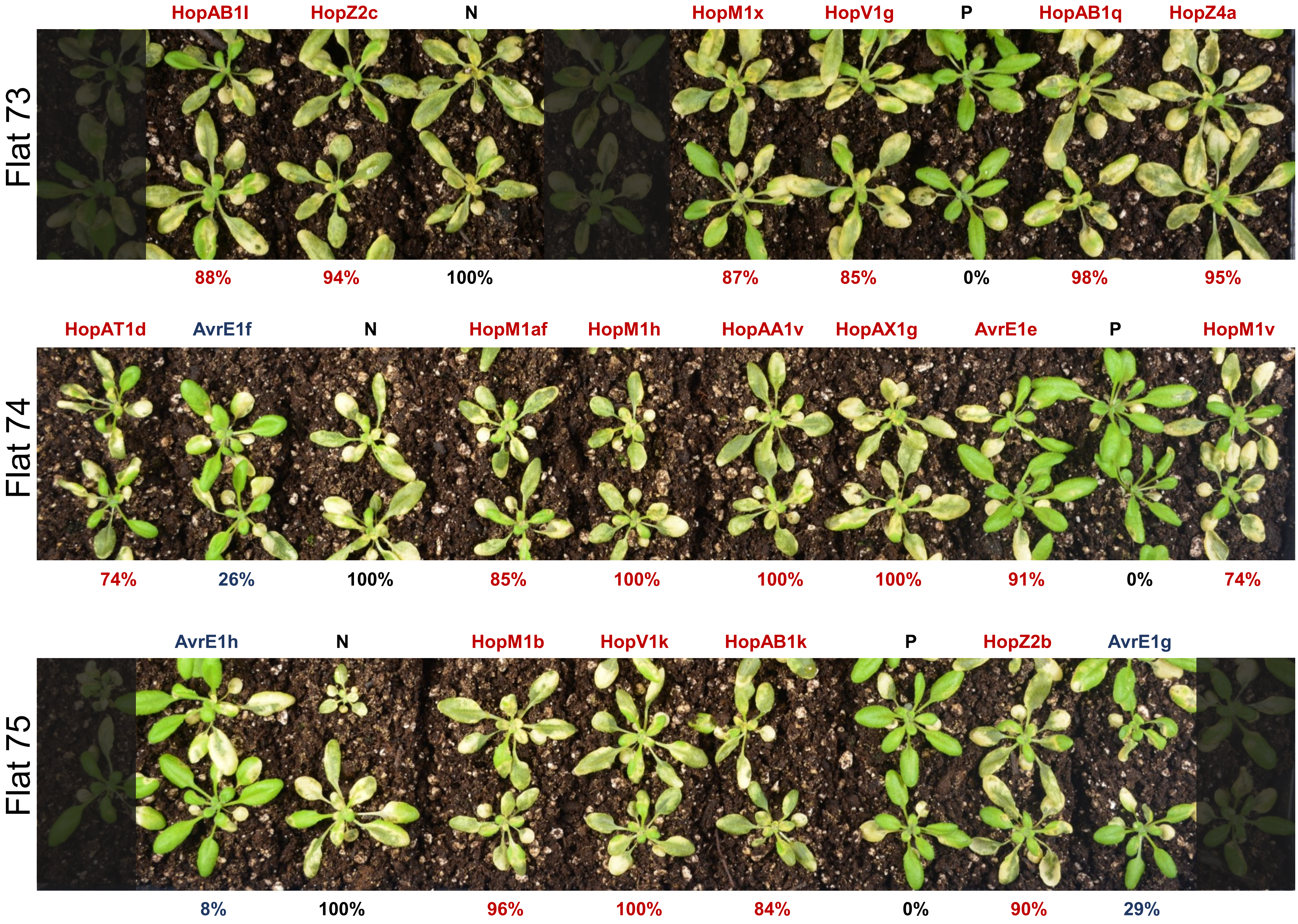
HopAB1l
HopZ2c
N
HopM1x
HopV1g
P
HopAB1q
HopZ4a
Flat 73
88%
94%
100%
87%
85%
0%
98%
95%
HopAT1d
AvrE1f
N
HopM1af
HopM1h
HopAA1v
HopAX1g
AvrE1e
P
HopM1v
Flat 74
74%
26%
100%
85%
100%
100%
100%
91%
0%
74%
AvrE1h
N
HopM1b
HopV1k
HopAB1k
P
HopZ2b
AvrE1g
Flat 75
8%
100%
96%
100%
84%
0%
90%
29%

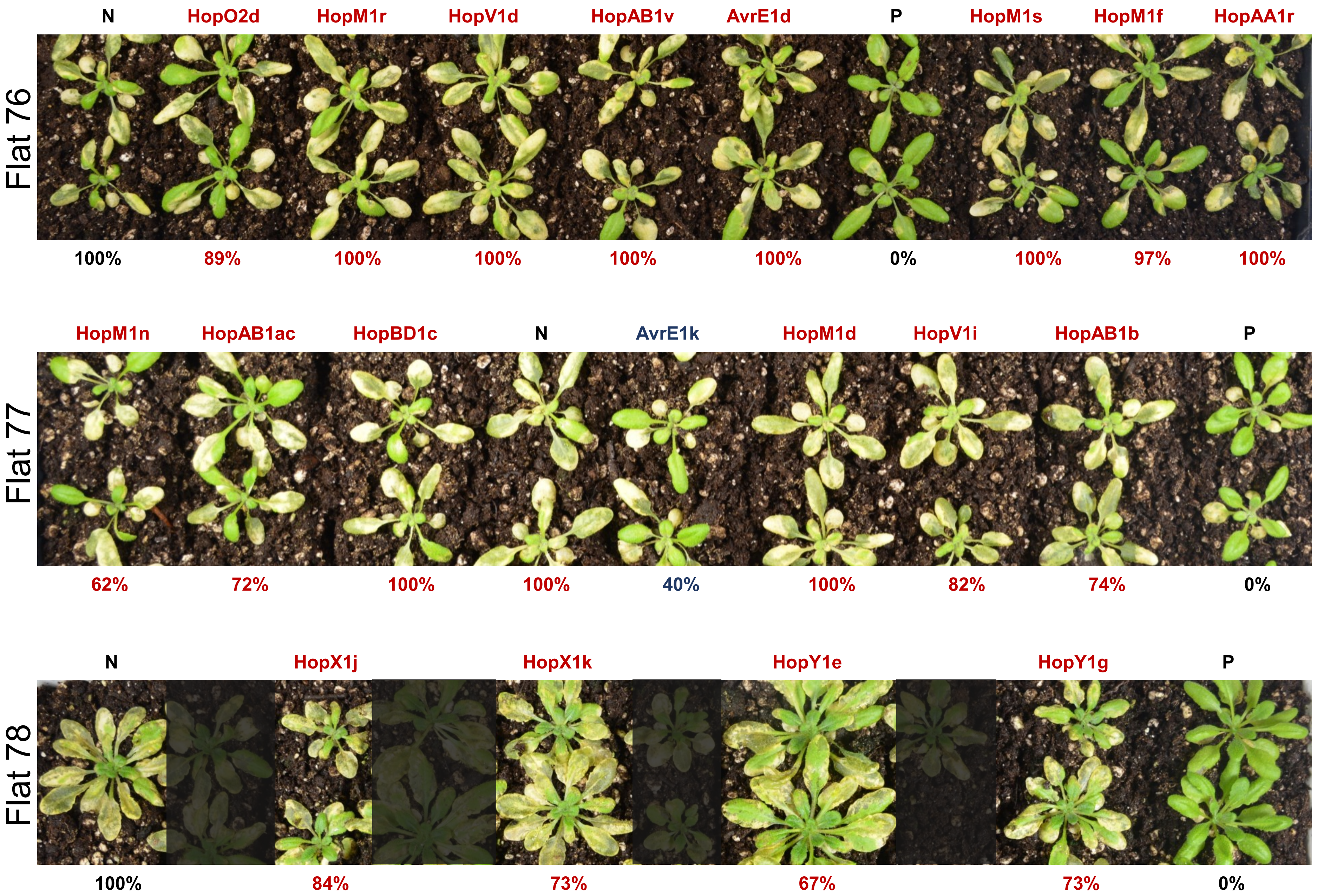

N HopO2d HopM1r HopV1d HopAB1v AvrE1d P HopM1s HopM1f HopAA1r
Flat 76
100% 89% 100% 100% 100% 100% 0% 100% 97% 100%
HopM1n HopAB1ac HopBD1c N AvrE1k HopM1d HopV1i HopAB1b P
Flat 77
62% 72% 100% 100% 40% 100% 82% 74% 0%
N HopX1j HopX1k HopY1e HopY1g P
Flat 78
100% 84% 73% 67% 73% 0%

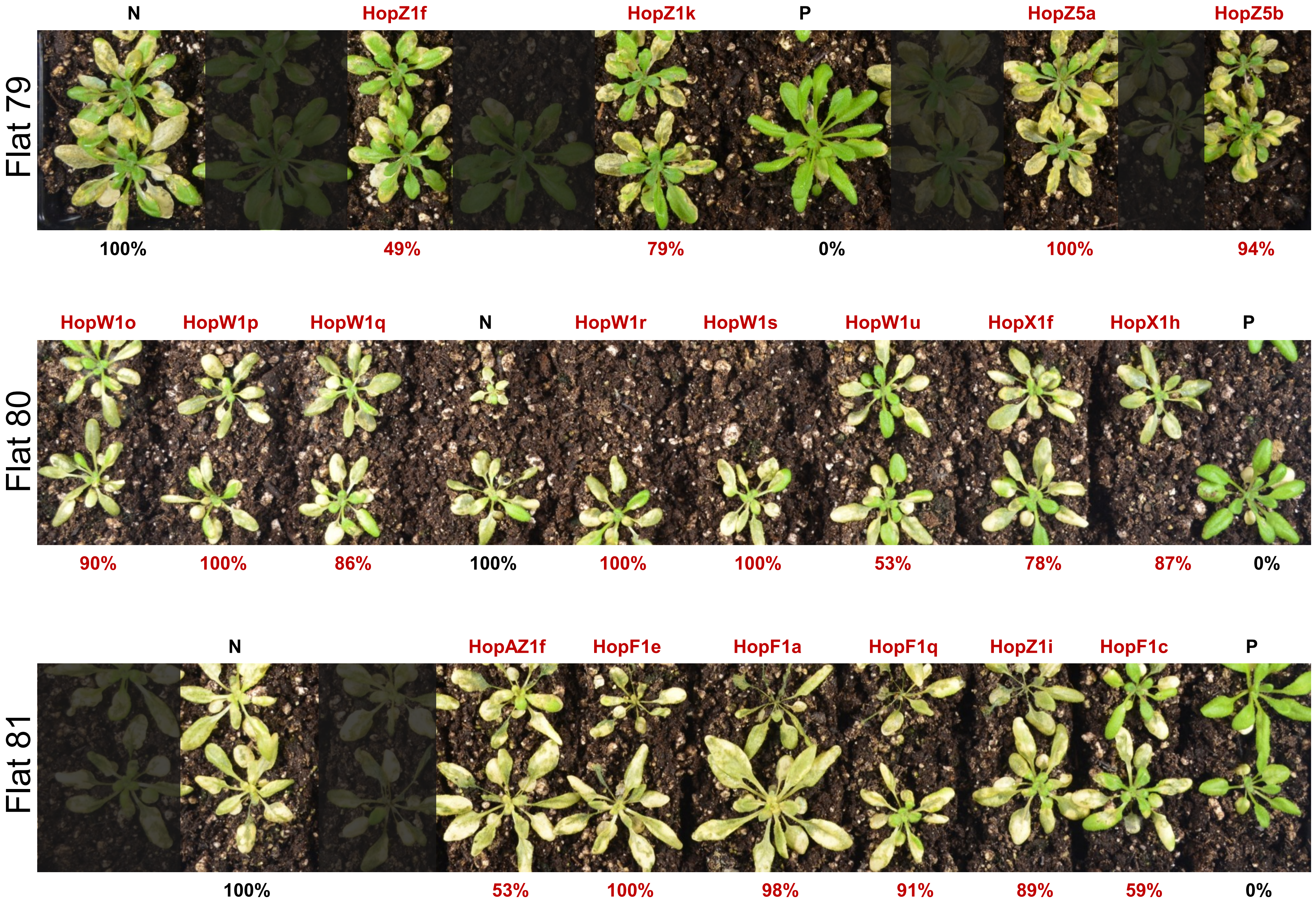
Flat 79
N
HopZ1f
HopZ1k
P
HopZ5a
HopZ5b
100%
49%
79%
0%
100%
94%
Flat 80
HopW1o
HopW1p
HopW1q
N
HopW1r
HopW1s
HopW1u
HopX1f
HopX1h
P
90%
100%
86%
100%
100%
100%
53%
78%
87%
0%
Flat 81
N
HopAZ1f
HopF1e
HopF1a
HopF1q
HopZ1i
HopF1c
P
100%
53%
100%
98%
91%
89%
59%
0%

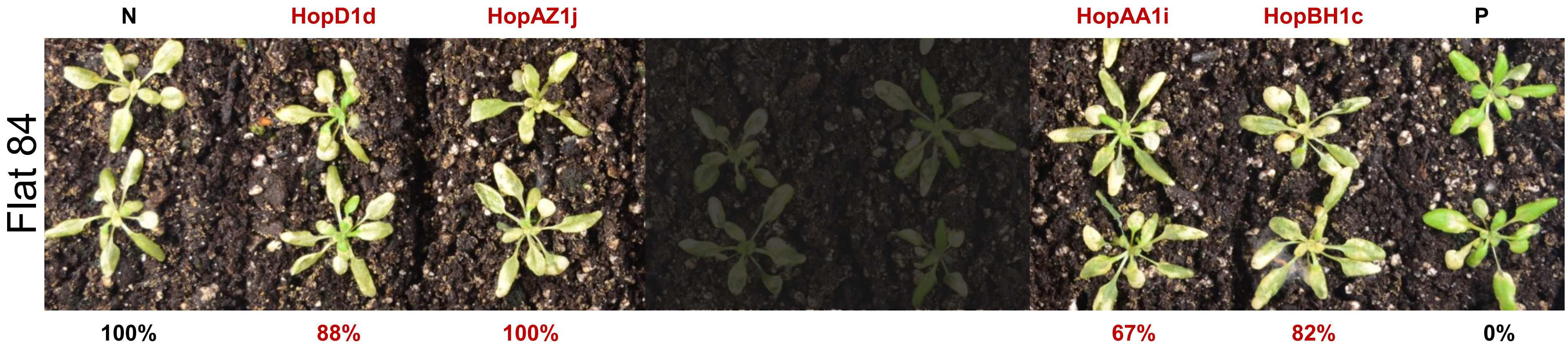

N
HopD1d
HopAZ1j
HopAA1i
HopBH1c
P
Flat 84
100%
88%
100%
67%
82%
0%
